# Gene-by-gene screen of the unknown proteins encoded on *P. falciparum* chromosome 3

**DOI:** 10.1101/2022.07.07.499005

**Authors:** Jessica Kimmel, Marius Schmitt, Alexej Sinner, Pascal Jansen, Sheila Mainye, Gala Ramón-Zamorano, Christa Geeke Toenhake, Jan Stephan Wichers, Jakob Cronshagen, Ricarda Sabitzki, Paolo Mesén-Ramírez, Hannah Michaela Behrens, Richárd Bártfai, Tobias Spielmann

## Abstract

Taxa-specific proteins are key determinants defining the biology of all organisms and represent prime drug targets in pathogens. However, lacking comparability with proteins in other lineages makes them particularly difficult to study. In malaria parasites this is exacerbated by technical limitations. Here, we analysed the cellular location, essentiality, function and, in selected cases, interactome of all unknown non-secretory proteins encoded on an entire *P. falciparum* chromosome. The nucleus was the most common localisation, indicating it is a hotspot of parasite-specific biology. More in-depth functional studies with four proteins revealed essential roles in DNA replication and mitosis. The novel mitosis proteins defined a possible orphan complex and a highly diverged complex needed for the spindle-kinetochore connection. Structure-function comparisons indicated that the taxa-specific proteins evolved by different mechanisms. This work demonstrates the feasibility of gene-by-gene screens to elucidate the biology of malaria parasites and reveal critical parasite-specific processes of interest as drug targets.

## INTRODUCTION

Malaria parasites are unicellular organisms with a complex life cycle. The intracellular development in red blood cells causes the symptoms of the disease that in the case of the *Plasmodium falciparum* parasite kills several hundred thousand people each year (World Health Organization, 2021). As eukaryotic organisms, the parasites share many core processes with the human host. However, one third of the parasite’s predicted proteins lacks detectable homology to proteins in other organisms and is still without annotated function. These taxa-specific proteins are of particular importance because they are likely to be involved in parasite-specific biology and may define traits specific to parasites and the genus. Due to their absence in the human host, taxa-specific essential proteins and functions are also of high interest because they represent potential drug targets that could be specifically inhibited without harming the host (Oberstaller et al., 2021; Rancati et al., 2018).

Proteins unique for a given taxonomic unit (often referred to as ‘orphans’) are responsible for species-specific traits and the evolution of major taxonomical branches (Bowles et al., 2020; Johnson and Tsutsui, 2011; Khalturin et al., 2008). Although these orphans are common in all taxa (Marsden, 2006), comparably few have been functionally studied so far (Tautz and Domazet-Lošo, 2011). There is a debate whether taxa-specific genes arose predominately de novo (from noncoding DNA regions) (Vakirlis et al., 2020) or whether the majority are simply orthologs that have evolved beyond recognition (Weisman et al., 2020). New functionalities can also arise through rearrangements of functional domains within and between genes (Cosby et al., 2021). The evolution beyond recognition scenario implies that function might still be assigned to ‘unknowns’ if prediction methods improve. For disease causing organisms, such as malaria parasites, it is even more important whether a potentially taxa-specific protein still has structural similarities with evolutionarily conserved proteins, as this would make it a less likely drug target. Hence, identifying parasite-specific proteins and functions, regardless of the evolutionary origin, is of high importance.

A further important factor is protein essentiality. To assign this on a large scale, genome-wide approaches have been carried out in *P. berghei*, and *P. falciparum* parasites (Bushell et al., 2017; Zhang et al., 2018). These approaches also permit functional and stage specific screens (Russell et al., 2021; Stanway et al., 2019; Thomas et al., 2016). However, they provide no data for the cellular location of individual proteins nor cell lines for their functional analysis. For instance, Zhang et al. (2018) provides a predicted score of essentiality for *P. falciparum* genes that – despite its usefulness – does not preclude experimental validation for individual targets of interest (Ishizaki et al., 2022). These latter issues are solved by gene-by-gene screens that however typically lack throughput in malaria parasites (Ishizaki et al., 2022; de Koning-Ward et al., 2015; Rancati et al., 2018). Taking advantage of the selection-linked integration (SLI) system for genomic modification (Birnbaum et al., 2017) we here carried out a proof-of-concept study covering all unknown ‘non-secretory’ proteins (i.e. all proteins not recruited into the ER or ER membrane) encoded on chromosome 3 of *P. falciparum* parasites. We established a resource of cell lines that for each target protein (i) provides a detailed localisation of the physiologically expressed protein, (ii) permitted us to conditionally inactivate it for functional studies and (iii) in selected cases was used to obtain interactomes using dimerisation-induced quantitative BioID (DiQ-BioID (Birnbaum et al., 2020)), all based essentially on one genome modification cell line per target. This non-biased approach indicated that ~50% of the target proteins were important for parasite growth and that the nucleus is a dominant site of taxa-specific proteins, highlighting it as a source of potential drug targets. In-depth functional analyses of four nuclear proteins revealed they had different functions in DNA replication and mitosis and indicated that taxa-specific proteins in malaria parasites arose by different means. This work shows the feasibility and usefulness of gene-by-gene screens in *P. falciparum* parasites and provides proof-of-principle to analyse all unknown proteins of this parasite.

## RESULTS

### Candidate selection and cellular localisation

We selected all proteins encoded by genes on chromosome 3 that at the start of this study were annotated as unknown (PlasmoDB v31) and for which there was evidence of expression in asexual blood stages (Le Roch et al., 2003) (Figure S1A). For technical reasons (due to the chosen inactivation approach) we excluded signal peptide- and transmembrane domain-containing proteins (‘secretory proteins’). BLAST searches were done to exclude proteins with homologies outside the taxa in cases where this was not reflected in the PlasmoDB (v31) annotation. Proteins with generic domains that did not give specific information on function (e.g., zinc fingers) were kept in the list. This resulted in 33 candidates termed Chr3C1-33 (Figure S1A; Table S1). To analyse their localisation and function, we C-terminally tagged the proteins by fusing the endogenous genes with the sequence encoding 2×FKBP-GFP-2×FKBP using SLI (Birnbaum et al., 2017) (Figure S1B). We obtained knock-in cell lines for 27 candidates, suggesting that six genes were refractory to C-terminal tagging (Figure S1C, D). The candidate proteins showed various subcellular localisations (Figure 1; Datafile 1). The largest group were nuclear proteins, either showing a uniform nuclear signal or defined foci (e.g., Chr3C7 and Chr3C16 which showed foci at the nuclear periphery) (Figure 1A). Four proteins were found in the vicinity of the nucleus that we classified as ‘nucleus proximal or Golgi’ (Figure 1A), of which co-localisation with GRASP confirmed a Golgi location for Chr3C3 and a Golgi proximal compartment for Chr3C18 (Figure S1E, F). Two proteins (Chr3C12 and Chr3C13) showed locations typical for functions in invasion (Figure 1A). Chr3C13 showed a focal localisation in schizonts and at the apical end of merozoites (Datafile 1), resembling apical organellar localisation. Co-expression with apical markers revealed no co-localisation with micronemes, rhoptry bulb or neck but a location even more apical than all these markers (Figure S1G), indicating that Chr3C13 is a novel apical protein located close to the apical tip. Chr3C12 was recently located at the inner membrane complex (IMC) (named PIC4) using this cell line (Wichers et al., 2021). Chr3C19 and Chr3C20 showed a localisation pattern reminiscent of the mitochondrion, however, only Chr3C19 (Figure 1A) co-localised with Mito-Tracker (Figure S1H, I). In total, 33% of the proteins were localised in the nucleus, 12% in the cytoplasm, 12% in proximity to the nucleus or Golgi, 3% comprised candidates at the IMC, the apical complex or the mitochondrion, and 21% showed unknown or complex localisation patterns (Figure 1B). Assessing expression of the endogenously tagged candidate proteins across blood stage development (Figure 1C), we observed varying patterns ranging from presence throughout the intraerythrocytic cycle to very narrow time windows (e.g., late-schizont specific expression of Chr3C25). In conclusion, this approach localised most of the non-secretory, unknown proteins of chromosome 3 in their physiological state through blood stage development and revealed an unexpectedly high proportion of nuclear proteins amongst them.

**Figure 1:**
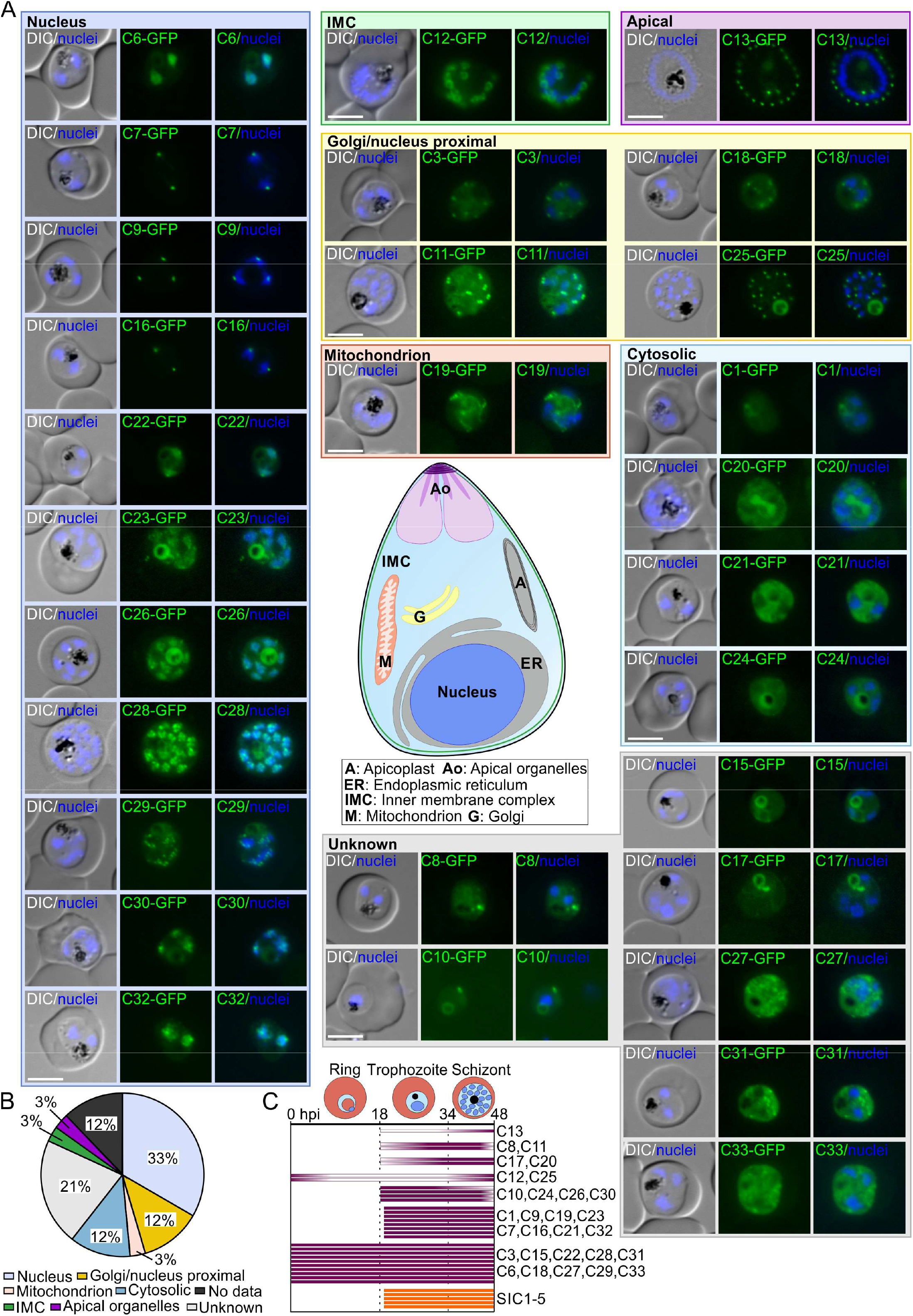
Subcellular localisations of unknown proteins encoded on chromosome 3. (A) Live cell fluorescence microscopy images showing the location of GFP-tagged candidates expressed from the endogenous locus (representative images of 3 independent microscopy sessions with at least 10 image series each). See Datafile 1 for full panels covering all asexual blood stages. Fluorescence patterns were assigned to subcellular structures of the parasite as indicated in scheme. (B) Pie chart displaying proportion of candidates with the indicated localisations. No cell lines were obtained for Chr3C2, Chr3C4, Chr3C5, Chr3C14 (‘no data’). (C) Expression timing of the physiological proteins (chromosome 3 screen candidates, purple; SICs, orange) determined from detected GFP signal during asexual blood stage development. Each bar represents one individual candidate.*DIC, differential interference contrast; nuclei were stained with DAPI; size bars, 5 μm; hpi, hours post invasion*.

### Importance for parasite survival (‘essentiality’)

To assess their essentiality and function, we conditionally inactivated all the tagged candidates (except for Chr3C12 already shown to be dispensable (Wichers et al., 2021)) using the knock sideways (KS) system (Birnbaum et al., 2017; Haruki et al., 2008; Robinson et al., 2010). Of the 26 candidates 22 were efficiently mislocalised upon induction of the KS, while the others showed no or only partial mislocalisation (Figure 2A, B; Figure S2A). Growth assays revealed an important function during blood stage development for eight of the proteins efficiently knocked aside (Figure 2A, C, D), whereas the others appeared to be dispensable (Figure S2A). For example, this revealed that Chr3C3 is a new essential Golgi protein, Chr3C29 is a likely nuclear pore protein with an important function and Chr3C7 and Chr3C16 are essential nuclear proteins. The novel apical tip protein Chr3C13 is required for invasion (Figure S2B, C).

**Figure 2:**
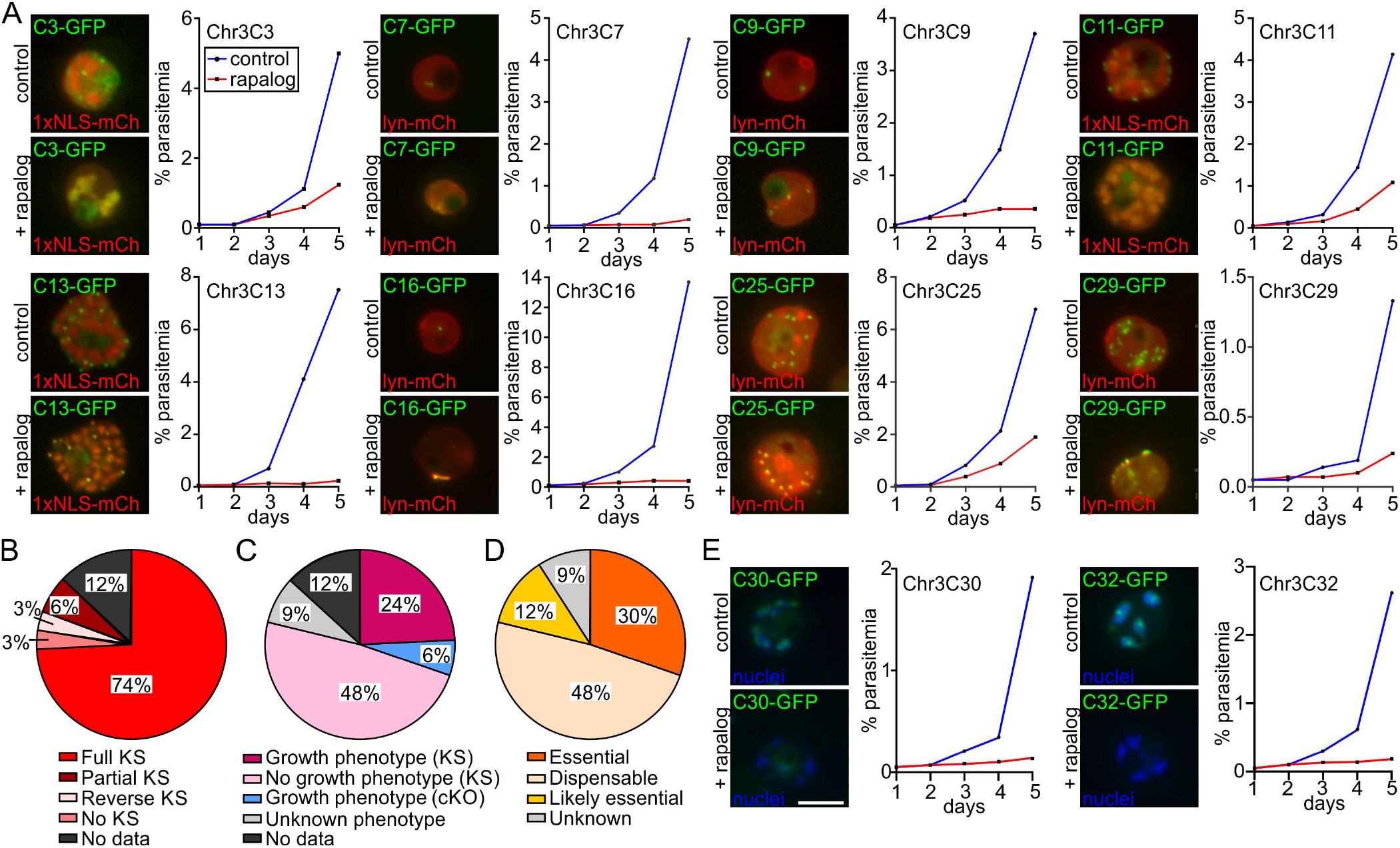
Conditional inactivation of candidates. (A) Live cell fluorescence microscopy images of KS with knock-in parasites episomally expressing either the lyn-FRB-mCherry (lyn-mCh) for mislocalisation to the parasite plasma membrane or 1×NLS-FRB-mCherry mislocaliser (1×NLS-mCh) for mislocalisation into the nucleus grown in absence (control) or presence of rapalog (+ rapalog, induced KS) for candidates where this resulted in a growth defect as indicated by the growth curves (parasitemia measured using flow cytometry). Images (17-24 h after addition of rapalog) are representatives of 3 independent microscopy sessions with at least 10 image series per session and condition; growth curves are representatives from 3 independent experiments (all replicates shown in Figure S2A). (B) Pie chart showing the KS efficiency of all tested candidates. Full mislocalisation, target protein undetectable at original localisation; no mislocalisation, no change in target protein localisation; partial mislocalisation, some of the target protein remained at original localisation in at least one replicate; reverse mislocalisation, re-localisation of the mislocaliser to the original site of the target protein; no data, no KS as not tagged. (C) Pie chart showing proportion of candidates with growth phenotypes by either KS or cKO. Unknown phenotype, candidates with partial or no mislocalisation; no data, candidates that could not be tagged. (D) Pie chart displaying proportion of essential and dispensable candidates based on KS or cKO results. Candidates that could not be C-terminally tagged were classified as likely essential. Unknown, Unknown phenotype, candidates with partial or no mislocalisation. (E) Live cell fluorescence microscopy images of Chr3C30 and Chr3C32 cKO parasites grown in absence (control) or presence of rapalog (+ rapalog, induced diCre based gene excision) for 24h are shown next to growth curves. Images and curves are representatives from 3 independent experiments (all growth experiments shown in Figure S3A, C) with at least 10 imaged parasites per imaging session and condition. *mCh, mCherry; size bars, 5 μm*.

The six candidates that could not be C-terminally tagged were also assumed as important for para-site growth as this was likely due to a detrimental influence of the tag on an essential function (Figure 2D). This assumption was confirmed for Chr3C30 and Chr3C32 using SLI for N-terminal tagging (Figure S1J, K) and diCre-based conditional knock-out (cKO) which resulted in a pro-found growth defect (Figure 2E; Figure S3A-C). Overall, this analysis revealed that 10 (30%) of the 33 candidates were essential for parasite survival, 4 (12%) likely essential, 16 (48%) dispensable and 3 (9%) could not be inactivated with the method used (Figure 2D, Table S1). Some of the candidates classified as dispensable may in principle still be important for parasite growth, considering limitations of the methodology (e.g., insufficient level of mislocalisation or site independent function). Due to the abundance of nuclear proteins, the nucleus may represent a site particularly rich in taxa-specific functions and we chose four of the essential nuclear candidates (Chr3C7, Chr3C16, Chr3C30 and Chr3C32) for an in-depth functional analysis.

### Chr3C30 and Chr3C32 function in DNA replication and mitosis

Chr3C30 and Chr3C32 showed a diffuse distribution in the nucleus with a focus in proximity of tu bulin accumulations at the nuclear periphery (Figure 3A, B), suggesting a spatial relationship with centromeres. The cKO of both Chr3C30 and ChrC32 resulted in a development arrest in schizonts (Figure 3C, D). The phenotype of Chr3C30 cKO included a severely disrupted nuclear morphology, misaligned microtubules, and a significantly reduced overall DNA content compared to control parasites of similar size (Figure 3E). Despite of this, we observed multiple small Hoechst foci in these parasites that usually were associated with a tubulin focus and contained less DNA than controls (Figure 3F, G), suggesting multiple division events despite omission of the S-phase. Chr3C30 is therefore likely needed for DNA replication. Interestingly, we also found multiple hemozoin clusters (Figure 3A, F) in Chr3C30 cKO parasites, indicating a fragmentation of the digestive vacuole which was unexpected for a nuclear protein and may be a downstream effect. HHPred (Gabler et al., 2020; Zimmermann et al., 2018) identified a PSF2 domain in Chr3C30, suggesting it is one of the four GINS complex subunits known from other taxa and in fact has recently been annotated as a GINS protein in PlasmoDB (v46; Figure 3H). Based on structural similarity of the Chr3C30 alpha fold model (Jumper et al., 2021; Varadi et al., 2022) with hsPSF2, we propose that Chr3C30 (PSF2 subunit) along with PF3D7_0523600, PF3D7_1335500, PF3D7_1303900 and PF3D7_0914800 (also predicted to contain GINS domains using HHPred) are the *P. falciparum* equivalents of the GINS complex where at least one of the subunits appears to have duplicated (Figure 3I; S3H). Our functional data for Chr3C30 is congruent with the role of the GINS complex in DNA replication and segregation in model organisms (Huang et al., 2005; MacNeill, 2010), indicating that Chr3C30 is an example of improved annotation, revealing the correct orthology.

**Figure 3:**
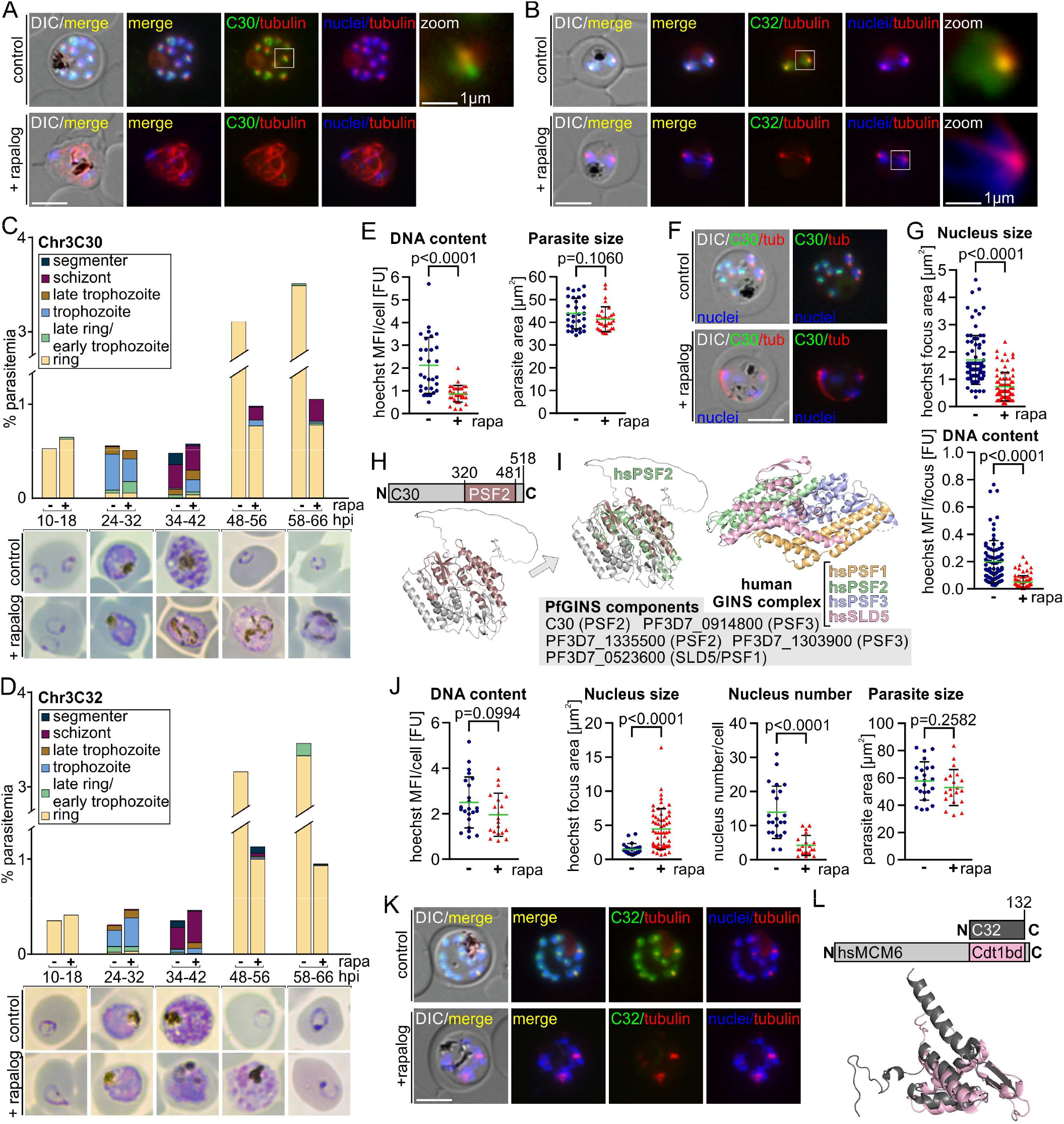
Functional analysis of Chr3C30 and Chr3C32. (A) Live cell fluorescence microscopy images (representatives of 3 independent microscopy sessions with at least 10 image series per session and condition) of Chr3C30 cKO late stage schizonts grown in presence (+ rapalog, induced diCre gene excision) or absence (control) of rapalog and co-stained with Tubulin Tracker Deep Red. White box enlarged section (zoom). See Figure S3D for full panels. (B) Live cell fluorescence microscopy images (representatives of 3 independent microscopy sessions with at least 10 image series per session and condition) of Chr3C32 cKO parasites after the first nuclear division after growth in presence (+ rapa, induced diCre gene excision) or absence (control) of rapalog and co-stained with Tubulin Tracker Deep Red. White box, enlarged section (zoom). See Figure S3E for full panels. (C, D) Stages and growth based on Giemsa smears (example images shown beneath the graph) in synchronised (0-8 h synchronisation window) in Chr3C30 cKO (C) or Chr3C32 cKO (D) parasites grown in presence (+ rapa, induced diCre gene excision) and absence (-rapa) of rapalog at the indicated time points (second biological replicate shown in Figure S3F, G). (E) Quantification of DNA content (left) in early stage Chr3C30 schizonts after cKO (+ rapa) compared to control (-rapa). Parasites of similar size (right, size based on DIC images) from four independent experiments were used (+ rapa n=30, -rapa n=32). Unpaired two-tailed t-test, P values are indicated, error bars are SD. (F) Example fluorescence images of Chr3C30 cKO parasites from analysis in (E) and (G). See Figure S3D for full panels. (G) Quantification of nucleus size (top) and DNA content per nucleus (bottom) in 10 parasites from (E); + rapa, induced diCre gene excision; - rapa, control. Unpaired two-tailed t-test, P values are indicated, error bars are SD. (H) Scheme (top; not to scale) and predicted protein structure by alphafold (bottom) of ChrC30 with PSF2 domain indicated. Numbers, aa positions. (I) Alignment of Chr3C30 protein structure from (J) with hsPSF2 protein structure (2Q9Q; RMSD=2.8 over 168 residues) and possible assignment to human GINS complex (2Q9Q; structure shown to the right). Other subunits of possible *P. falciparum* GINS subunits are indicated (Top hit for domains identified by HHPred indicated in brackets). (J) Quantification of DNA content, nucleus size and number of nuclei in Chr3C32 schizonts after cKO (+ rapa) compared to control (-rapa). Parasites of similar size (graph on the right, size based on DIC images) from two independent experiments (+ rapa n=23, - rapa n=20). Unpaired two-tailed t-test, P values are indicated, error bars are SD. (K) Example fluorescence images of Chr3C32 cKO parasites from analysis in (L). See Figure S3E for full panels. (L) Scheme (top; not to scale) of Chr3C32 which corresponds to the C-terminal CDT1 binding domain (CDT1 bd) of human MCM6. Alignment (bottom) of Chr3C32 predicted protein structure (by alphafold) with protein structure of CDT1 binding domain of human MCM6 (2KLQ; RMSD=5.3 over 88 residues). *DIC, differential interference contrast; tubulin, stained with Tubulin Tracker Deep red; nuclei stained with Hoechst 33342; merge, merged green (GFP), red (Cy5) and blue (Hoechst) channels; Size bars, 5 μm (if not indicated otherwise); hpi, hours post invasion; MFI, mean fluorescence intensity; FU, arbitrary fluorescence units*.

In contrast to Chr3C30, the cKO of Chr3C32 resulted in a lower number of nuclei that were of significantly larger size but the total DNA content per parasite was not affected in late schizonts of similar size (Figure 3J, K), suggesting a nuclear division defect. HHPred indicated that Chr3C32 corresponds to the CDT1 binding domain of the DNA replication licensing factor MCM6 (Figure 3L). However, a ‘full-size’ MCM6 (PF3D7_1355100) already exists in *P. falciparum*, indicating that Chr3C32 is a novel protein corresponding to the CDT1 binding domain alone that may have arisen from a duplication. As our functional data revealed an important function of Chr3C32 in mitosis rather than DNA replication (the function Ctd1 and MCM6 typically are associated with), Chr3C32 may be an example of a domain that acquired a new function.

### Chr3C7 is a potential orthologue of SKA1 or SKA3

The detailed characterisation of the growth defect after KS-based conditional inactivation of Chr3C7 revealed no obvious negative impact on ring and trophozoite development, but the parasites failed to complete schizogony (Figure 4A). Analysis of schizonts with inactivated Chr3C7 showed an amorphous nuclear mass without clearly separated nuclei and elongated and misaligned micro-tubules (Figure 4B). The number of discernible nuclei per cell was lower and they were of a larger size compared to the control while overall DNA content was not affected, indicating a nuclear division defect (Figure 4C). To obtain more detailed functional insight, we assessed the first nuclear division that was reported to be the longest of all division rounds (Simon et al., 2021) and is not obscured by the presence of multiple nuclei. With on-set of nuclear division, the Chr3C7 focus co-localised with accumulated tubulin signal (Figure 4D), likely associated with the DNA-free region at the nuclear periphery (Simon et al., 2021) and was located beneath the extranuclear centriolar plaques (CPs) represented by CEN3 (Simon et al., 2021) at the nuclear periphery (Figure 4E). As mitosis proceeded, Chr3C7 was associated with the typical mitotic microtubular structures (Figure 4D) and duplicated Chr3C7 foci were found between duplicated CPs (Figure 4E). Inactivation of Chr3C7 resulted in aberrant mitotic spindles and interpolar spindle microtubules while the nuclei remained connected after anaphase (Figure 4F), resembling previously observed anaphase chromatin bridges (Liffner and Absalon, 2021). Thereafter, the para-sites lacking Chr3C7 were unable to equally segregate DNA into daughter nuclei and the microtubule ends became misaligned (Figure 4F). While the localisation of CPs was not affected during the first mitosis after inactivation of Chr3C7 (Figure 4G), they became clustered in the parasite periphery during progressing schizogony. This indicates that CP duplication continues, leading to accumulations not connected to the nuclei (Figure 4H). While in the control Chr3C7 colocalised with the kinetochore marker NDC80 (Zeeshan et al., 2020) during schizogony, absence of Chr3C7 resulted in disorganised kinetochores that became distributed along the microtubules (Figure 4I). Overall, these results indicate that Chr3C7 facilitates proper connection of microtubules to chromosomes and is required for chromosome segregation during mitosis in blood stage schizogony. A HHPred search with Chr3C7 (Figure 4J) identified two regions with homologies to spindle and kinetochore-associated proteins (SKA) as the top hits and this protein has in the meantime been annotated as SKA2 in PlasmoDB (v46). In human cells, the SKA complex (consisting of SKA1, SKA2 and SKA3) connects spindle microtubules with the kinetochores and loss of SKA resulted in delayed chromosome separation (Hanisch et al., 2006; Jeyaprakash et al., 2012; Welburn et al., 2009), reminiscent of the phenotype observed here. A structural alignment with the Chr3C7 protein structure predicted by alphafold revealed that the N-terminal SKA domain of Chr3C7 shows similarity with the conserved N-terminal region present in all human SKA proteins (Figure 4J, K). The C-terminal winged helix domain of Chr3C7 overlaps with the hsSKA1 microtubule-binding domain (MTBD) (Figure 4J, K) that is also present in hsSKA3 but not in hsSKA2. Based on this and the functional data, we therefore conclude that Chr3C7 likely is a functional orthologue of SKA1 or SKA3 (henceforth referred to SKA1/3).

**Figure 4:**
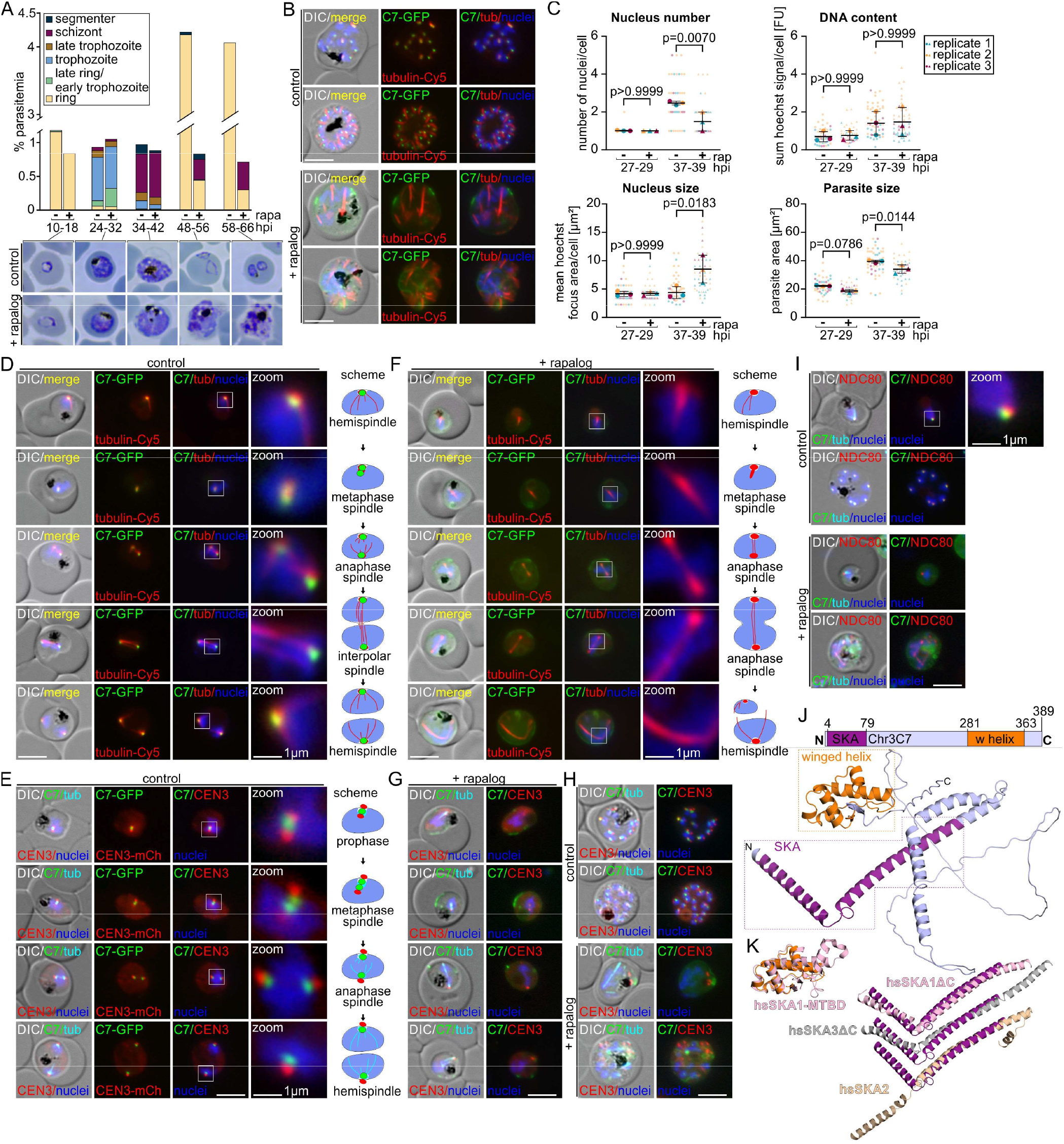
Functional analysis of Chr3C7. (A) Stages and growth based on Giemsa smears (example images shown beneath the graph) in synchronised (0-8 h synchronisation window) Chr3C7 KS parasites grown in presence (+ rapa, to induce Chr3C7 KS) and absence of rapalog (-rapa) at the indicated time points (second biological replicate shown in Figure S4A). (B) Live cell fluorescence microscopy images (representative images of 3 independent microscopy sessions with each at least 10 image series per condition) of Chr3C7 KS late stage schizonts grown in absence (control) and presence of rapalog (+ rapalog, to induce Chr3C7 KS) co-stained with Tubulin Tracker Deep Red. See Figure S4B for full panel. (C) Quantification of nuclei number, nucleus size, DNA content and parasite size in synchronised (0-2 h synchronisation window) Chr3C7 parasites after KS (+ rapa) compared to control (-rapa) before (27-29 hpi) and after (37-39 hpi) first nuclear division from 3 independent experiments. See Figure S4C for graph containing all time points and n numbers. Multiple paired t-tests (matched repeated-measures one-way ANOVA with assumed sphericity) and Bonferroni correction for multiple testing, P values indicated; error bars are SD. (D and F) Live cell fluorescence microscopy images (representative images of 3 independent microscopy sessions with at least 10 image series per session and condition) of synchronised (0-2 h synchronisation window) Chr3C7 KS parasites grown in absence (D, control) and presence (F, + rapalog, induced Chr3C7 KS) of rapalog and co-stained with Tubulin Tracker Deep Red during first nuclear division. Typical spindle types and duplication of Chr3C7 focus (D) and elongated anaphase spindles and incomplete nuclear segregation (F) were observed (zoom and indicated in the schemes). See Figure S4B for full panels. (E and G) Live cell fluorescence microscopy images (representative images of 3 independent microscopy sessions with at least 10 image series per session and condition) showing co-localisation with centriolar plaque marker CEN3-mCherry in synchronised (0-2 h synchronisation window) Chr3C7 KS parasites grown in absence (E, control) and presence of rapalog (G, + rapalog, to induce Chr3C7 KS) and co-stained with Tubulin Tracker Deep Red during first nuclear division. CEN3 focus duplication occurred before Chr3C7 focus duplication (E, zoom and indicated in the scheme) and became dissociated from nucleus after Chr3C7 KS (G). See Figure S4D for full panels. (H and I) Live cell fluorescence microscopy images (representative images of 3 independent microscopy sessions with at least 10 image series per session and condition) of selected cells from asynchronous parasites showing co-localisation with centriolar plaque marker CEN3-mCherry (H) or kinetochore marker NDC80-mCherry (I) in Chr3C7 KS schizonts grown in absence (control) and presence of rapalog (+ rapalog, to induce Chr3C7 KS) and co-stained with Tubulin Tracker Deep Red. See Figure S4D for full panels. (J) Scheme (top, not to scale) and predicted protein structure by alphafold (bottom) showing Chr3C7 with SKA and winged helix domain (w helix); numbers indicate aa positions. (K) Alignment of Chr3C7 SKA domain (left) from (J) with structure of SKA domain of hsSKA1ΔC (4AJ5; RMSD=4.8 over 72 residues), hsSKA2 (4AJ5; RMSD=3.9 over 56 residues) and hsSKA3ΔC (4AJ5; RMSD=4.32 over 72 residues) and Chr3C7 winged helix domain (right) with microtubule-binding domain (MTBD) of hsSKA1 (4C9Y; RMSD=3.0 over 80 residues). *Size bars, 5 μm (if not indicated otherwise); DIC, differential interference contrast; tub/tubulin, stained with Tubulin Tracker Deep red; nuclei stained with Hoechst 33342; hpi, hours post invasion; FU, arbitrary fluorescence units*.

### Interactome of Chr3C7 defines a highly diverged SKA complex

To further clarify the role of Chr3C7 as the potential SKA1/3 in *P. falciparum* parasites, we carried out DiQ-BioID (Birnbaum et al., 2020; Kimmel et al., 2021) to identify interactors and compartment neighbours. This resulted in a list of candidates including many known and suspected nuclear residents and nuclear division proteins such as NDC80 (Zeeshan et al., 2020), kinesins (Zeeshan et al., 2019), and NUF2 (Zeeshan et al., 2020) (Figure 5A, B, S5 A-H) but also many high confidence hits annotated as unknown in PlasmoDB (v46). For validation, we selected the top hits including NDC80 and five unknown proteins termed SKA1/3-interacting candidates (SIC1-5) and analysed their localisation and function as done for the candidates of the screen (Figure 5; Table S1). Conditional inactivation of endogenously tagged NDC80 as well as SIC1-5 using KS revealed NDC80 and all SICs, except for SIC3, to be essen-tial for blood stage growth and resulted in a similar phenotype to that observed in schizonts lacking functional SKA1/3 (Figure 5C). NDC80, SIC1, SIC2, SIC4 and SIC5 co-localised with SKA1/3 and tubulin in the nucleus, confirming these SICs as interactors or compartment neighbours (Figure 5D). In contrast, while SIC3 was found in the nucleus, it was dispersed and not accumulated in SKA foci (Figure 5D), indicating it may be a false positive. Accordingly, SIC3 was the only tested candidate not consistently detected in all DiQ-Bi-oID replicates (Figure 5B; Figure S5D-H). Further supporting a direct functional connection, the inactivation of SIC1, SIC2, SIC4 and NDC80 resulted in elongated microtubules that were disconnected from nuclei and SKA1/3 appeared to be stuck at the misaligned microtubules, indicating a disconnection of SKA1/3 from the chromosomes (Figure 5D, E). Overall, this indicated an important contribution of these proteins in the SKA-dependent connection of microtubules to the chromosomes during nuclear division. Interestingly, SKA1/3 clustered with SIC1 and SIC2 as the most enriched proteins in all DiQ-BioID replicates (Figure 5A, B; Figure S5D-H). HHPred searches revealed homology of a region of SIC1 to human SKA proteins as the top hit while SIC2 did not reveal any high confidence hits. However, alphafold predicted very long α-helices similar to those in human SKA proteins (Jeyaprakash et al., 2012) in both SIC1 and SIC2, although the reliability of these predictions was rather low (Figure S5J). Along with the functional data, this indicates that SIC1 and SIC2 may correspond to the other two SKA subunits that together with SKA1/3 correspond to a highly diverged SKA complex. The hypothesis of these three proteins forming a complex was further supported by co-mislocalisation of episomal SKA1/3 after KS induction of endogenously tagged SIC1 or SIC2 (Figure 5F). The SKA1/3 phenotype can nevertheless be considered specific, as complementation with an episomally functional copy of Chr3C7 rescued the KS (Figure S4F). These results indicate a model of the kinetochore-microtubule interface in *P. falciparum* parasites that involves the SKA complex with a similar function to human cells (Figure 5G) but evolved in a unique way and also contains parasite-specific proteins.

**Figure 5:**
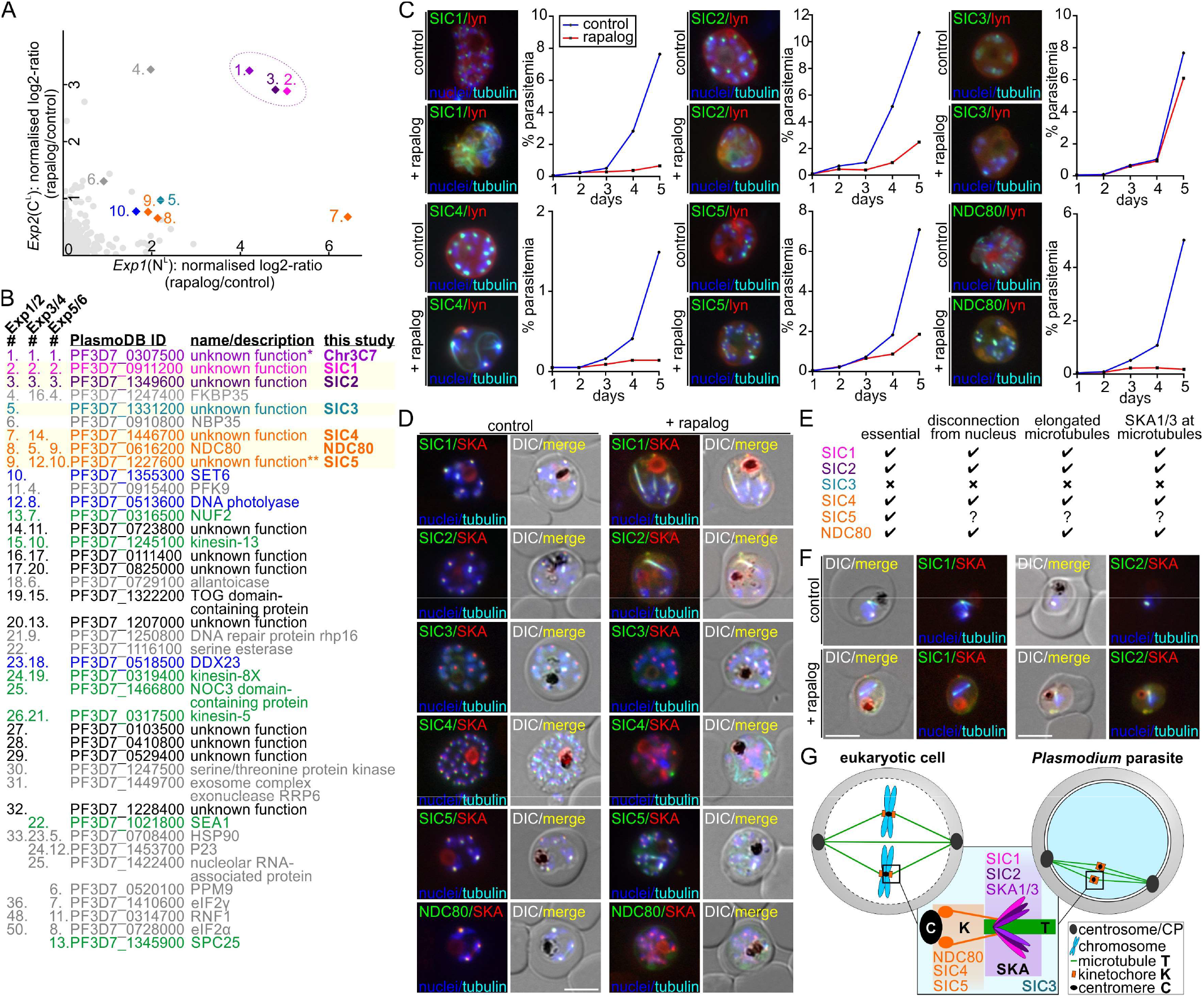
Chr3C7 interactome reveals SKA complex. (A) Top-right quadrant of scatter plot of Chr3C7 DiQ-BioID, showing proteins enriched (log2 ratio) on rapalog (biotinyliser on target) compared with control (biotinyliser in cytoplasm). Significantly enriched proteins are indicated (details in B). Purple circle highlights cluster of SKA1/3, SIC1 and SIC2. See Figure S5D-H for full plot and replicates. (B) Rank number according to enrichment, PlasmoDB ID, name and description, and name in this study of DiQ-BioID hits from 6 independent experiments shown in A and Figure S5D-H. Common DiQ-BioID contaminants (Birnbaum et al., 2020) are indicated in grey, top six candidates chosen for validation are highlighted (SICs, NDC80, colour code as in (A)). Colour code of further hits: proteins expected to be involved in mitosis, green; nuclear resident proteins, blue. *, annotated as SKA2 in PlasmoDB 46 and here designated SKA1/3; ** annotated as SPC24 (Zeeshan et al., 2020). (C) Representative fluorescence microscopy images of KS with SIC1-5 and NDC80 knock-in parasites episomally expressing the lyn-FRB-mCherry mislocaliser (lyn) grown in absence (control) or presence (+ rapalog, induced KS of SIC1-5/NDC80) of rapalog for 17-24 h and stained with Tubulin Tracker Deep Red are shown next to growth curves (parasitemia determined using flow cytometry). Images and curves are representatives from 3 independent experiments (see Figure S5I for full panels and Figure S2D for all growth experiments) with at least 10 image series per session and condition. (D) Live cell fluorescence microscopy images (representative images of 3 independent microscopy sessions with at least 10 image series per session and condition) showing co-localisation of SKA1/3 in SIC1-5 and NDC80 KS late schizonts grown in absence (control) and presence (+ rapalog, induced KS of SIC1-5/NDC80) of rapalog and co-stained with Tubulin Tracker Deep Red. See Figure S5I for full panels. (E) Effects on microtubules and SKA1/3 localisation after KS of SIC1-5 and NDC80 as evident from images in D and Figure S5I. Note that SIC5 KS resulted in reverse mislocalisation, preventing unequivocal assessment of the phenotype. (F) Live cell fluorescence microscopy images (representative images of 3 independent microscopy sessions with at least 10 image series per session and condition) showing colocalisation of SKA1/3 in SIC1 KS (left) or SIC2 KS (right) parasites grown in absence (control) and presence (+ rapalog, induced SIC1 or SIC2 KS) of rapalog and co-stained with Tubulin Tracker Deep Red. See Figure S5I for full panels. (G) Schematic representation of microtubule-kinetochore interface during metaphase in eukaryotic model organisms (closed mitosis, chromosome condensation) and *Plasmodium* parasites (open mitosis, uncondensed chromosomes). According to interactome (A, B) and functional data (D, E), SKA1/3, SIC1 and SIC2 in *Plasmodium* parasites correspond to SKA complex (purple) in other eukaryotes. SIC4 is a likely new kinetochore protein and kinetochore function of SIC5 (SPC24) and NDC80 was confirmed in this study. SIC3 is not connected to this functional site. *Size bars, 5μm; DIC, differential interference contrast; tubulin, stained with Tubulin Tracker Deep red; nuclei stained with Hoechst 33342; merge, merged GFP (green), mCherry (red), Cy5 (cyan) and Hoechst (blue) channels*.

### Chr3C16 is a novel, taxa-specific protein involved in mitosis

Analysis of the parasites after KS-based Chr3C16 inactivation showed that the growth phenotype was due to a defect in schizont development (Figure 6A). Growth was rescued by episomal complementation (Figure S6J). Schizonts lacking Chr3C16 contained a lower number of nuclei than controls (Figure 6B, C), suggesting a role of this protein in nuclear division. Tracking the first mitosis event showed that Chr3C16 appeared as a bell-shaped focus that was associated but not overlapping with tubulin accumulations and duplicated before separation of the nuclei (Figure 6D, F). After conditional inactivation of Chr3C16, only hemi-spindles and tubulin accumulations were visible in the time frame of the first nuclear division (Figure 6E, F), resulting in only one nucleus that had doubled in size (Figure 6G), indicating a mitosis defect that with further development manifested as a mitotic delay. Cell growth was not halted, and S-phase was not affected (Figure 6G), pinning the function of Chr3C16 to early mitosis between prophase and metaphase. Chr3C16 foci doubled before SKA1/3 (Figure 6H) and inactivation of Chr3C16 did not appear to affect SKA1/3, but fewer SKA1/3 foci were visible in later stages, resulting from the mitotic delay phenotype (Figure 6I, J). The Chr3C16 bell structure was always located at the side of CEN3 foci facing the DNA and the duplication of Chr3C16 and CPs occurred at the same time during mitosis (Figure 6K). Whereas the inactivation of Chr3C16 had no effect on CP duplication during the first mitosis (Figure 6L), we observed large unseparated nuclei with multiple CP foci in later stages (Figure 6M), indicating that the function of Chr3C16 is independent from CEN3. Overall, these results indicated an important function of Chr3C16 early in mitosis that could not be connected to any process known from model organisms. We therefore carried out DiQ-BioID to obtain information to which process it might belong to from interactors or compartment neighbours (Figure S6H-J). This resulted in a list of 14 significantly enriched proteins (99% percentile) that (apart from the likely false positives) intriguingly had no annotated function (Figure 5N). Of note, PF3D7_0724600 is a putative protein kinase related to Ark kinases that is of unknown function (Adderley et al., 2021). The absence of defining similarities indicated that Chr3C16 is a mitosis protein that with its interactors either evolved beyond recognition or represents a truly orphan functionality in this process.

**Figure 6:**
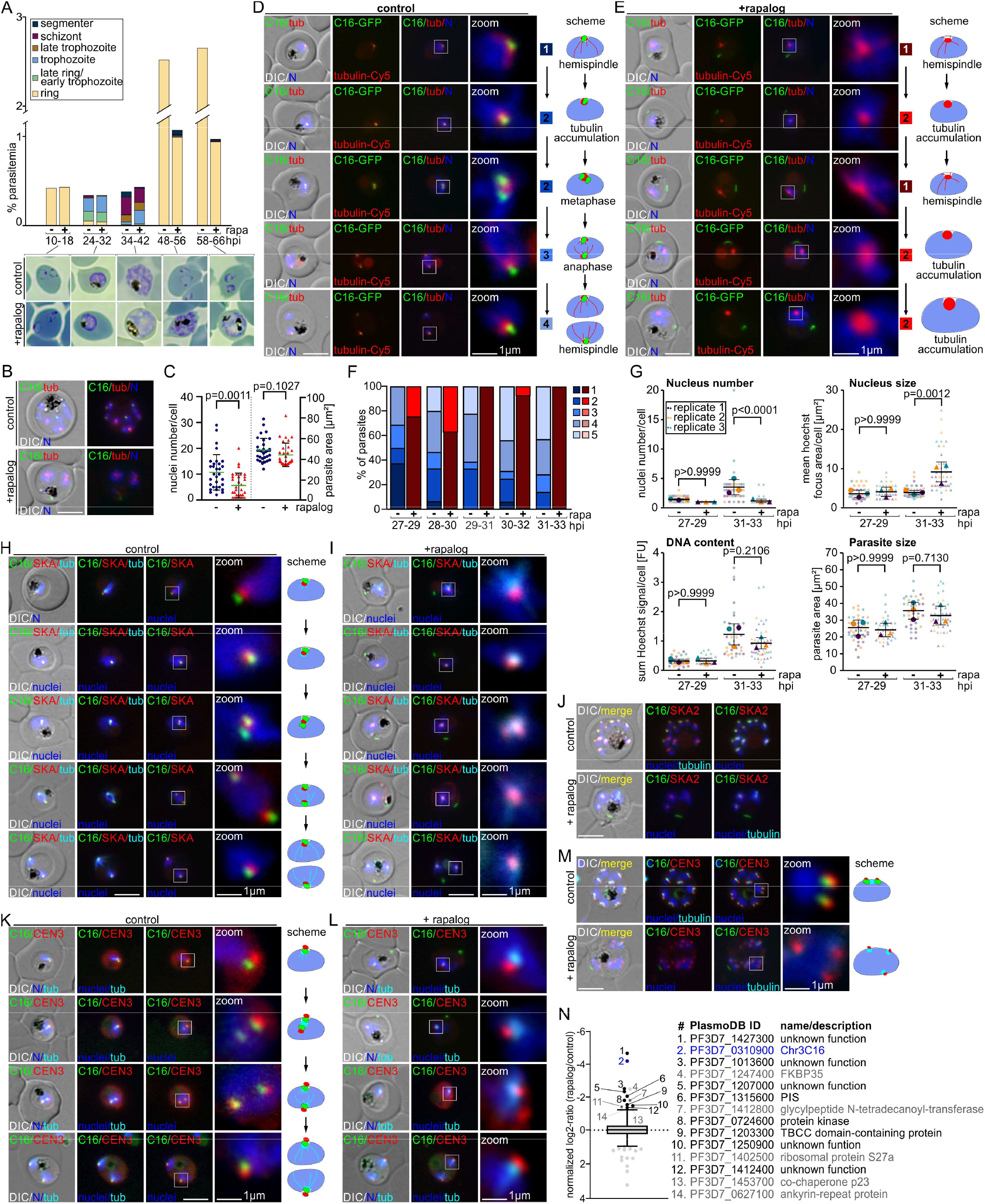
Functional analysis of Chr3C16. (A) Stages and growth based on Giemsa smears (example images shown beneath the graph) in synchronised (0-8 h synchronisation window) Chr3C16 KS parasites grown in presence (+ rapa, induced Chr3C16 KS) and absence (-rapa) of rapalog at the indicated time points (second biological replicate shown in Figure S6A). (B) Representative fluorescence microscopy images of Chr3C16 KS late stage schizonts grown in absence (control) and presence (+rapalog, induced Chr3C16 KS) of rapalog and co-stained with Tubulin Tracker Deep Red. See Figure S6B for full panels. (C) Quantification of nuclei number in Chr3C16 KS parasites after KS (+ rapa) compared to control (-rapa). Parasites of similar size (right, size based on DIC images) from 5 independent experiments were used (+ rapa n=38, - rapa n=37). Unpaired two-tailed t-test, P values are indicated, error bars are SD. (D, E) Live cell fluorescence microscopy images (representative images of 3 independent microscopy sessions with at least 10 image series per session and condition) of synchronised (0-2 h synchronisation window) Chr3C16 KS parasites grown in absence (D, control) and presence (E, + rapalog, induced Chr3C16 KS) of rapalog and co-stained with Tubulin Tracker Deep Red during the first nuclear division. Schematic representation next to zoom images indicate typical spindle types (red), duplication of Chr3C16 focus (green) and nuclear segregation (blue) that were observed in controls (D) whereas Chr3C16 KS resulted in alternating hemispindles and tubulin foci without nuclear division (E). See Figure S6C for full panels. According to these images, parasites were classified into five stage (1-5) of tubulin phenotypes during mitosis. (F) Quantification of synchronised parasites (0-2 hours synchronisation window) grown with (+ rapalog, induced Chr3C16 KS) or without rapalog (-rapalog) showing the 5 phases defined in (D, E) at the indicated time points during the first nuclear division. Stage 5 parasites were defined as all parasites with more than two nuclei or with two nuclei where a new round had been started. One representative of 3 independent experiments shown in Figure S6D. (G) Quantification of nuclei number, nucleus size, DNA content and parasite size in synchronised (0-2 h synchronisation window) Chr3C16 parasites after KS (+ rapalog) compared to control (-rapalog) at time points before (27-29 hpi) and after (31-33 hpi) the first nuclear division from the same 3 independent experiments from D, E, F. See Figure S6E for graph containing all time points and n numbers. Multiple paired t tests (matched repeated-measures one-way ANOVA with assumed sphericity and Bonferroni correction for multiple testing), P values are indicated, error bars are SD. (H and I) Live cell fluorescence microscopy images (representative images of 3 independent microscopy sessions with at least 10 image series per session and condition) showing co-localisation with SKA1/3-mCherry in synchronised (0-2 synchronisation window) Chr3C16 KS parasites grown in absence (H, control) and presence (I, + rapalog, induced Chr3C16 KS) of rapalog and co-stained with Tubulin Tracker Deep Red during first nuclear division. Zoom and schematic representation showing Chr3C16 focus duplication prior to SKA1/3 focus duplication (H) and effect on SKA1/3 of mitotic delay after Chr3C16 KS (I). See Figure S6F for full panels. (J) Live cell fluorescence microscopy images (representative images of 3 independent microscopy sessions with at least 10 image series per session and condition) showing co-localisation with SKA1/3-mCherry in Chr3C16 KS late schizonts grown in absence (control) and presence (+ rapalog, induced Chr3C16 KS) of rapalog and co-stained with Tubulin Tracker Deep Red. See Figure S6F for full panels. (K and L) Live cell fluorescence microscopy images (representative images of 3 independent microscopy sessions with at least 10 image series per session and condition) showing co-localisation with centriolar plaque marker CEN3-mCherry in synchronised (0-2 h synchronisation window) Chr3C16 KS parasites grown in absence (K, control) and presence (L, + rapalog, induced Chr3C16 KS) of rapalog and co-stained with Tubulin Tracker Deep Red during first nuclear division. Chr3C16 focus and CEN3 focus duplicate simultaneously (K, zoom and scheme) and CEN3 duplication was not affected after Chr3C26 KS (L, zoom and scheme). See Figure S6G for full panels. (M) Live cell fluorescence microscopy images (representative images of 3 independent microscopy sessions with at least 10 image series per session and condition) showing co-localisation centriolar plaque marker CEN3-mCherry in Chr3C16 KS late schizonts grown in absence (control) and presence (+ rapalog, induced Chr3C16 KS) of rapalog and co-stained with Tubulin Tracker Deep Red. Zoom and schematic representation indicates association of Chr3C16 and CEN3 at the centriolar plaque (control) and multiple CEN3 foci at nuclei after Chr3C16 KS (+ rapalog). See Figure S6G for full panels. (N) Box plot of Chr3C16 DiQ-BioID showing proteins enriched (log2 ratio) on rapalog (biotinyliser on target) compared with control (biotinyliser in cytoplasm) from one experiment. Significantly enriched proteins (99% percentile) are indicated with numbers (graph) and PlasmoDB IDs and annotation in table. Likely false positives (including known DiQ-BioID false positives (Birnbaum et al., 2020)) are in grey. *Size bars, 5 μm (if not indicated otherwise); DIC, differential interference contrast; tubulin, stained with Tubulin Tracker Deep red; nuclei stained with Hoechst 33342; merge, merged green, red and blue channels; hpi, hours post invasion; FU, arbitrary fluorescence units*.

## DISCUSSION

Here we show that gene-by-gene analysis of all unknown non-secretory proteins of an entire *P. falciparum* chromosome is feasible, indicating that the technical capacity for a similar analysis of all unknown proteins of this parasite is coming into reach. The resulting cell lines provide versatile options to study the localisation and expression timing of the physiologically expressed target proteins, their function by conditional inactivation and their interactome using DiQ-BioID. As with any system there are some drawbacks, e.g. that the KS levels were insufficient for functional analysis of some targets. While diCre is an alternative that can be used while still taking advantage of SLI (Birnbaum et al., 2017) and could include also secretory proteins, the need for synthesis of re-codonised gene fragments increases the cost. KS also has the advantage to be a very fast acting system by directly targeting the protein, reducing secondary effects that obscure phenotypic assessment (Birnbaum et al., 2017, 2020; Hoeijmakers et al., 2019; Jonscher et al., 2019; Robinson et al., 2010).

Here this was harnessed to obtain mechanistic insight into cell division of the parasite through the analysis of two newly identified protein groups involved in mitosis. We identified the three likely equivalents of SKA complex subunits (SKA1-3) and thereby found that malaria parasites use a highly taxa specific SKA complex for segregation of chromosomes during schizogony, instead of the functional analogous DASH/Dam1 complex known from e.g. yeast (van Hooff et al., 2017). We assigned other previously unknown proteins to a function with SKA1-3 at the kinetochore-microtubule interface, as evident from SKA1/3 remaining associated with the elongated microtubules that lost connection to chromosomes after inactivation of these proteins. While not all of the possible interaction candidates were validated, the high rate of confirmed and plausible hits (e.g. SICs, NDC80 (Zeeshan et al., 2020) and kinesins (Zeeshan et al., 2019)) indicates that further unknown proteins in this list are connected to this process. In contrast to the SKA complex which appears to be a highly adapted functionality that partially resembles that in other organisms, Chr3C16 had a function in mitosis upstream of SKA and neither through its sequence nor its DiQ-BioID interactome could be assigned to any known process. This group of proteins may therefore correspond to true orphans with a taxa-specific function. We show that Chr3C30 and Chr3C32 function in DNA replication and mitosis, respectively, with Chr3C30 being a probable component of the GINS complex while Chr3C32 consists of a single domain that has separated from its original function as a part of the larger protein MCM6. Hence, this work indicates that in malaria parasites the proteins originally annotated as unknown have different evolutionary origins: proteins that strongly diverged from evolutionarily related proteins but retained a similar function (SKA complex, GINS), proteins with new domain combinations (Chr3C32) and potential true orphans (Chr3C16 complex).

This screen revealed a high proportion of nuclear proteins, highlighting the nucleus as a major site of taxa-specific processes. Essential nuclear proteins may therefore represent a key vulnerability of the parasite that could be exploited for the design of novel drugs. Assuming chromosome 3 is representative, a similar approach with the entire genome is expected to identify another ~240 novel nuclear proteins. The nuclear proteins identified here showed a partial overlap with nuclear proteins from a targeted approach (Oehring et al., 2012) (Table S1). Our non-discriminatory gene-by-gene screen to analyse the localisation and function of unknown proteins therefore complements targeted approaches such as those based on organelle fractionations. Gene-by-gene screens also generate unbiased markers that might improve computational assignment of protein localisation obtained from global experimental and *in silico* approaches.

This work shows the power of combining localisation, functional analysis and interactomes with highly sensitive *in silico* methods such as HHPred or comparison with alpha fold structures to arrive at functions of individual or entire groups of proteins. Hence, using these *in silico* predictions alone will further reduce the number of unknown proteins in this parasite but only functional studies will give definitive support for a predicted function.

## Supporting information

Datafile 1

## ACKNOWLEDGEMENTS

We thank Bärbel Bergmann (BNITM) for help with DiQ-BioID extract preparation, the FACS core facility (BNITM) for providing flow cytometry equipment and service, Tim Gilberger (CSSB) for providing apical marker plasmids and PlasmoDB for their *Plasmodium* Informatics Resources. We thank Jacobus Pharmaceuticals for providing WR99210. DSM1 (MRA-1161) was obtained from MR4/BEI Resources (NIAID, NIH). J.K. and M.S. thank the Jürgen Manchot Stiftung and J.C. the Joachim Herz Stiftung for funding. S.M., G.R.-Z., R. B. and T.S. acknowledge funding by the Leibniz Association (K328/2020). H.M.B. was supported by a Ortrud Mührer Fellowship of the Vereinigung der Freunde des Tropeninstituts Hamburg e.V. P.J. and the proteomic facility (Department of Molecular Biology, Radboud University) are supported by the Oncode Institute and the Dutch Cancer Society.

## AUTHOR CONTRIBUTIONS

Conceptualization, T.S.; Methodology, J.K., M.S., P.J., S.M., G.R.-Z., J.S.W., J.C., R.B., and T.S.; Validation, J.K., M.S., A.S., P.J., S.M., G.R.-Z., J.S.W., C.G.T., J.C., P.M.-R., and H.B.; Formal Analysis, J.K., M.S., P.J., S.M., G.R.-Z., J.S.W., C.G.T., H.B., and T.S.; Investigation, J.K., M.S., A.S., P.J., S.M., G.R.-Z., J.S.W., C.G.T., J.C., P.M.-R., and H.B.; Resources, J.S.W. and R.S.; Writing – Original Draft, J.K. and T.S.; Writing – Review & Editing, J.K., M.S., S.M., G.R-Z., J.S.W., H.B., R.B., and T.S.; Visualization, J.K., M.S., A.S., S.M., J.S.W., and T.S.; Supervision, R.B., and T. S.; Funding Acquisition, R.B. and T.S. All authors read and approved the manuscript.

## DATA AVAILABILITY

All data are available in the manuscript or the supplementary materials. Further information, any additional information required to reanalyse the data reported in this paper and requests for cell lines, plasmids, resources, and reagents should be directed to and will be fulfilled by the lead contact, Tobias Spielmann.

## METHODS

### *P. falciparum* parasite culture

*P. falciparum* parasites (clone 3D7) (Walliker et al., 1987) were cultured in RPMI1640 containing 0,5% Albumax (Life Technologies) with a haematocrit of 5% in human 0+ erythrocytes at 37°C (in an atmosphere consisting of 5% O_2_, 5% CO_2_, 90% N_2_) according to standard procedures (Trager and Jensen, 1976).

### Red blood cells

Human transfusion blood concentrates (0+) were commercially purchased from Universitätsklinikum Hamburg-Eppendorf (Approval number 10569a/96-1). One ml of transfusion blood concentrate contained 0.5-0.7 ml of human erythrocytes, 0.28-0.44 ml SAG-M (consisting of 9.0 mg glucose monohydrate, 8.77 mg NaCl, 5.25 mg mannitol and 0.17 mg adenine per ml dH2O), 0.015 - 0.05 ml of human plasma, and 0.005 - 0.01 ml stabilizing solution (consisting of 26.3 mg sodium citrate, 25.5 mg glucose monohydrate, 3.27 mg citric acid monohydrate and 2.51 mg sodium dihydrogen phosphate dihydrate per ml dH2O). Blood concentrates are anonymous, and age or sex of blood donors were not known.

## METHOD DETAILS

### Cloning of plasmid constructs

To generate the plasmid for the integration cell lines of Chr3C2, Chr3C4, Chr3C6, Chr3C7, Chr3C9, Chr3C14, Chr3C21, Chr3C33, SICs and PfNDC80 (PF3D7_0616200) the C-terminal fragments before the stop codon (targeting regions) were amplified by PCR (length of targeting regions and primers in Figure S2) and cloned into pSLI-sandwich (Birnbaum et al., 2017) using NotI/AvrII (NEB). For the remaining candidate genes, the targeting regions (Figure S2) were synthesised and cloned into the pSLI-sandwich by Life Technologies GmbH (Darmstadt, Germany). For N-terminal SLI of Chr3C30 and Chr3C32, an N-terminal targeting region of Chr3C30 and Chr3C32 genomic sequence was amplified by PCR (primers in Figure S2) and inserted into the p-N-SLI-sandwich-K13-loxP plasmid (Birnbaum et al., 2017) using with NotI/PmeI (NEB), replacing the *kelch13* sequence. A functional and codon-changed version of Chr3C30 and Chr3C32 was synthesised (Genscript), amplified by PCR (primers in Figure S2) and cloned into the construct using AvrII/StuI (NEB). For KS, the 1×NLS-FRB-mCherry-DHODH^nmd3^ and lyn-FRB-mCherry-DHODH^nmd3^ mislocaliser plasmids (Birnbaum et al., 2017) and for di-Cre-based excision the pSkipFlox plasmid (Birnbaum et al., 2017) were used. Some KS experiments with Chr3C7 were carried out using the modified mislocaliser plasmid pLyn-FRB-T2A-mCherry-yDHODH^nmd3^ (Hoeijmakers et al., 2019). For co-localisation experiments of selected candidates, the Golgi-marker plasmid pGRASP-mCherry-BSD^nmd3^ (Birnbaum et al., 2020) and the apical marker (Cabrera et al., 2012; Geiger et al., 2020; Healer et al., 2002; Ito et al., 2019; Knuepfer et al., 2014; Peterson et al., 1989) plasmids pARL-^ama1^ARO-mCherry-yDHODH, pARL-^ama1^RON12-mCherry-yDHODH and pARL-^ama1^AMA1-mCherry-yDHODH (Wichers et al., 2021) were used. To obtain plasmids for co-localisation experiments while simultaneously carrying out KS of nuclear proteins as well as for complementation of Chr3C7 and Chr3C16 KS, the 1×NLS sequence of the marker-mCherry^sf3a2^-1xNLS-FRB-yDHODH^nmd3^-T2A plasmid was replaced by the sequence encoding MGCIKSKGKDSAGA (lyn, (Tóth et al., 2012). Subsequently, the full-length markers PfCEN3 (PF3D7_1027700), PfNDC80 (PF3D7_0616200), Chr3C7/SKA and Chr3C16 were amplified from 3D7 gDNA (primer sequences in Figure S2) and cloned into the construct, resulting in plasmids expressing CEN3-mCherry^sf3a2^-lyn-FRB-yDHO-DH^nmd3^-T2A, NDC80-mCherry^sf3a2^-lyn-FRB-yDHO-DH^nmd3^-T2A, Chr3C7/SKA-mCherry^sf3a2^-lyn-FRB-yDHODH^nmd3^-T2A and Chr3C16-mCherry^sf3a2^-lyn-FRB-yDHODH^nmd3^-T2A, respectively. For DiQ-Bi-oID, the BirA*-N^L^ and BirA*-C^L^ constructs (Birnbaum et al., 2020) were used. All constructs were cloned using Gibson assembly (Gibson et al., 2009) and the absence of unwanted mutations in inserts was verified by sequencing.

### *P. falciparum* parasite transfection, SLI and confirmation of verification of correct integration

Percoll-purified (Rivadeneira et al., 1983) late schizont stage parasites were transfected by electroporation with 50 μg of purified plasmid DNA (Qiagen) using the Amaxa system (Lonza Nucleofector II AAD-1001N, program U-033) as previously described (Moon et al., 2013). Transfectants were selected with 4 nM WR99210 (Jacobus Pharmaceuticals), 0.9 μM DSM1 (BEI resources) or 2 μg/μl blasticidin S (Life Technologies). For the selection of integrant parasites, SLI was done as described (Birnbaum et al., 2017) by adding G418 (ThermoFisher) at a final concentration of 400 μg/mL (C-terminal modification) or 0.9 μM DSM1 (N-terminal modification) to the culture after reappearance of transfectants carrying the episomal plasmid. After parasitemia recovered under G418 selection, correct integration was assessed using genomic DNA and PCR across the integration junctions and the absence of unmodified gene locus in the knock-in parasites was verified by PCR as previously described (Birnbaum et al., 2017).

### Induction of knock sideways and diCre excision

As previously described (Birnbaum et al., 2017), KS or diCre-based gene excision (cKO) was induced by splitting the culture into two dishes and addition of 250 nM rapalog (AP21967 (Clontech) diluted 1:20 in RPMI from a 500 mM stock dissolved in ethanol; Birnbaum et al., 2017) to the culture medium of one dish (+rapalog) while the other dish served as control. If not indicated otherwise, mislocalisation after KS of the candidates or absence of Chr3C30 and Chr3C32 after diCre excision (as compared with the control culture) was confirmed microscopically 17-24 h after addition of rapalog.

### Parasite growth assays

To test the essentiality of the candidates, the growth (parasitemia) of KS or diCre-based gene excision parasite cultures compared to controls was monitored every 24 h over 5 days using a previously established flow cytometry based assay (Birnbaum et al., 2017). For each flow cytometry measurement on day 1-5, the culture was resuspended, 20 μl was removed and mixed with 80 μl RPMI containing Hoechst 33342 (Cayman) and dihydroethidium (Cayman) at a final concentration of 4.5 μg/ml and 5 μg/ml, respectively. The sample was incubated for 20 minutes at room temperature followed by adding 400 μl of 0.003% glutaraldehyde in cold RPMI. The parasitemia was measured with a LSRII flow cytometer by counting 100,000 events using the FACSDiva software (BD Biosciences). On day 1, the parasitemia of an asynchronous culture was measured and adjusted to 0.05% or 0.1% parasitemia, split into two 2 ml dishes and KS or diCre-based gene excision was induced in one dish while the other served as a control. Growth curves were prepared in GraphPad Prism (version 9).

To assess stage specific growth phenotypes, ring stage parasites were synchronised to 10-18 hpi by incubation with 5% sorbitol (Lambros and Vanderberg, 1979) two times 10 hours apart. The parasites were brought back into culture and KS or diCre excision was induced. Giemsa smears were prepared at time points 10-18 hpi, 24-32 hpi, 34-42 hpi, 48-56 hpi (6-14 hpi in 2^nd^ cycle) and 58-66 hpi (16-24 hpi in 2^nd^ cycle) and fresh culture medium (containing 250 nM rapalog where applicable) was added at each time point. For each time point and condition 800 to 27000 red blood cells were counted manually and the parasitemia and stages were recorded.

### Invasion and egress assay

The invasion/egress assay was adapted from previously published assays (Geiger et al., 2020; Wichers et al., 2019). Briefly, Chr3C13 KS late schizonts were purified using Percoll (Rivadeneira et al., 1983), followed by an incubation period of 30 minutes at 37°C in the shaking incubator (800 rpm) allowing the parasites to reinvade new erythrocytes. Subsequently, the cells were transferred to static standard culture conditions, incubated for 3.5 hours and sorbitol synchronization (Lambros and Vanderberg, 1979) was performed resulting in tightly synchronized parasite culture with a time window of 4 h. These parasites were grown for another 20 hours at 37°C and KS was induced. Both cultures were incubated at 37°C until Giemsa-stained smears were prepared at the time points 38-42 hpi (‘pre-egress’) and 48-52 hpi (‘post-egress’). The number of merozoites per schizont was assessed by live fluorescence microscopy counting the number of Hoechst-stained nuclei per parasite in the ‘pre-egress’ timepoint in 16-23 (mean 20.5) schizonts per condition. The numbers of ring and schizont stage parasites were determined by counting parasites in 2023-4360 (mean 2570) erythrocytes per condition in randomly selected sight field and used to calculate the percentage of ruptured schizonts and number of new rings per ruptured schizont in the ‘post-egress’ samples.

### Live cell imaging

Live cell imaging of parasites endogenously expressing GFP and/or episomally expressing mCherry was carried out as previously described (Grüring and Spielmann, 2012) by placing the parasites on a glass slide and covering them with a cover slip. To visualise parasite nuclei, parasites were incubated with 1 μg/μl 4’,6’-diamidine-2’-phenylindole dihydrochloride DAPI (Roche) or 50 ng/ml Hoechst 33342 (as indicated in the figure legends) in RPMI medium for 10 min at 37°C. For tubulin staining the parasites were harvested and resuspended in medium containing 1:1000 Tubulin Tracker™ Deep Red (Thermo Fisher Scientific; resolved in DMSO according to manufacturer’s instructions). The cells were incubated for 20 minutes at 37°C in the shaking incubator (800 rpm) and 50 ng/ml Hoechst 33342 was added for additional staining of nuclei. The parasites were imaged with a Zeiss AxioImager M1 or M2 equipped with a Hamamatsu Orca C4742-95 camera and using either a 100× /1.4–numerical or a 63×/1.4– numerical aperture lens. Images were taken using the AxioVision software (version 4.7). The brightness and intensity of images was adjusted, and the channels were merged in Corel Photo-Paint (version X6). Corel Draw (version X6) was used for preparing the figures.

### Mitosis assay

To visualise parasites during their first nuclear division round, tightly synchronised parasites (2 hours window) were generated. To achieve this, late-schizont stage parasites were purified using 60% Percoll density centrifugation (Rivadeneira et al., 1983) and incubated in an appropriate volume of blood (50 μl per timepoint to be imaged) and medium (1:1 ratio) for 30 minutes at 37°C in the shaking incubator (800 rpm) allowing merozoites to invade into new erythrocytes. Subsequently, the cells were transferred to static standard culture conditions and incubated for 90 minutes. The remaining non-ruptured schizonts were lysed with 5% sorbitol to obtain 0-2 hpi rings stage parasites. The cells were brought back into culture by splitting into two 2 ml dishes per timepoint to be imaged to permit removing one culture for imaging without removing the other cultures from 37°C and KS or diCre excision was induced. At each timepoint during the experiment, starting at 27-29 hpi until the first nuclear division was completed (for 6 hours if not indicated otherwise), the parasites were harvested, incubated with medium containing 50 ng/ml Hoechst 33342 and 1:1000 Tubulin Tracker™ Deep Red for 20 minutes and placed on a glass slide covered with a cover slip for imaging.

For quantification, the parasites were scored by microscopy based on number and position of GFP foci, tubulin signal and number of nuclei into five pre-defined stages of mitosis at each timepoint and condition (n numbers and numbers of biological replicas indicated in the figure legends). All parasites were scored except of those that did not express the mislocaliser. The following parameters were measured using ImageJ version 2 (Schneider et al., 2012): parasite area from DIC pictures, Hoechst signal area representing the nuclei and Hoechst mean integrated density from the same selected area in UV-light channel pictures. Subsequently, the area was moved away from nuclei to measure Hoechst background fluorescence. To calculate total Hoechst signal per cell representing the DNA content, the mean integrated intensity values were corrected for background and summed up for all nuclei to obtain the total for each individual cell. The nucleus size of Chr3C30 schizonts cultured without rapalog as controls (Figure 3E) was determined from one representative nucleus per cell. The mean parasite size, nucleus size and Hoechst signal from each timepoint and condition was applied for statistical analysis as indicated in the figure legend using GraphPad Prism (version 9).

### DiQ-BioID

DiQ-BioID experiments were carried out as previously described (Birnbaum et al., 2020). Knock-in parasite culture episomally expressing the biotinyliser constructs were split into two identical cultures (150 ml each) and dimerisation was induced in one culture by adding rapalog (final concentration 250 nM). RPMI supplemented with 50 μM biotin (Sigma-Aldrich) was added to both cultures. After 12 h culturing, medium and supplements were replaced. Cultures were harvested after a total of 24 h culturing with biotin and washed twice in 1 × DPBS. Parasites were purified from the host cells using 0.03% saponin in 1x DPBS on ice for 10 minutes, washed five times in 1 × DPBS, lysed in 2 ml lysis buffer (50mM Tris-HCL pH 7.5, 500 mM NaCl, 1% Triton-X-100) containing 1 mM DTT, 80 μL of 25× protein inhibitor cocktail (Roche) and 1 mM PMSF and the lysate was frozen at −80 °C. For purification of biotinylated proteins, lysates were frozen and thawed two times and centrifuged at 16,000 g for 10 minutes. 50 μL Streptavidin Sepharose (GE Healthcare) was added to the lysate and incubated by rotating overnight at 4 °C. The beads were washed twice in lysis buffer, once in dH2O, twice in Tris-HCl (pH 7.5) and three times in 100 mM Triethylammonium bicarbonate buffer pH 8.5 (TEAB, Sigma-Aldrich) before treatment with 50 μL elution buffer (2 M Urea in 100 mM Tris pH 7.5 containing 10 mM DTT). Elution was performed at room temperature, shaking for 20 minutes. Subsequently, iodoacetamide (IAA) was added to a final concentration of 50 mM and the samples were further incubated in the dark, shaking for 10 minutes. The proteins were then treated with Trypsin/LysC 0,1 μg/μl, while shaking at room temperature. After two hours, the beads were pelleted, the supernatants containing eluted proteins were collected and the beads were rinsed with an extra 50 μL of elution buffer. After for 5 minutes at room temperature, the supernatant was pooled with the previous elution. The final 100 μL of eluted proteins were added 1 μL of Trypsin/LysC and treated overnight while shaking at room temperature.

The samples were dimethyl-labelled on C18 stage-tips (Rappsilber et al., 2007) and stored at 4 °C until analysed via mass spectrometry. Preparation of the C18 stage tips involved washing the membrane once with 50 μL methanol, once with 50 μL buffer B (80% acetonitrile, 0.1% formic acid) and twice with 50 μL buffer A (0.1% formic acid) before loading the overnight-digested peptides and washing once again with buffer A. Biological replicates were processed under label-swap conditions, where in the forward experiments (Chr3C7 experiments 1, 3 and 5), control samples were labelled with the “light” label and the rapalog-treated samples carried the “heavy” label whereas in the reverse experiments (Chr3C7 DiQ-BioID experiments 2, 4, 6 and the Chr3C16 DiQ-BioID experiment), control samples carried the “heavy” label and the rapalog-treated samples were labelled with the “light” label. The “light” labelling buffer was prepared dissolving 2 mg of sodium cyanoborohydride powder (NaBH_3_CN, Sigma-Aldrich) per millilitre of dimethyl labelling buffer (10 mM monosodium phosphate, 35 mM disodium phosphate) containing 0.2 % of formaldehyde solution (CH_2_O, Sigma-Aldrich). The “heavy” labelling buffer was prepared as the “light” but using 2 mg/ml sodium cyanoborodeuteride (NaBD_3_CN, Sigma-Aldrich) in dimethyl labelling buffer, and replacing the CH_2_O for 0.2 % of formaldehyde-^13^C, d2 solution (^13^CD_2_O, Sigma-Aldrich). To each stage tip sample loaded with peptides, 300 μL of the appropriate labelling buffer was eluted through, followed by a final wash with 100 μL of buffer A.

### Mass spectrometry and data analysis

On the day of measuring the samples by mass spectrometry, each C18 stage tip was rehydrated with 30 μL of buffer A and peptides were eluted in 30 μL of buffer B. For each experiment, “light” and “heavy”-labelled samples were eluted into a single tube. The acetonitrile was evaporated in a SpeedVac and the concentrated sample was then reconstituted to a final volume of 12 μL with buffer A. After resuspension, 5 μL of sample was loaded onto a 30 cm C18-reverse phase column (1.8 μm Reprosil-Pur C18-AQ) and eluted using an Easy-nLC 1000 (Thermo Fisher Scientific) over a 114 min gradient (7.2% Acetonitrile/0.1% formic acid-25.6% acetonitrile/0.1% formic acid) and directly injected into an Orbitrap Fusion mass spectrometer (Thermo Fisher Scientific). Data was acquired in data-dependent top speed mode in a 3 sec cycle with dynamic exclusion set at 60 sec. Resolution for MS was set at 120.000.

Raw mass spectrometry data were processed using MaxQuant (version 1.5.3.30) and as described in Birnbaum et al. (2020) and summarised here. Parameters were set to default except for the following: Multiplicity was set at 2 and the labels were added to lysine residues and peptide N-termini (light: dimethLys0 and dimethNter0; heavy: dimethLys8 and dimethNter8). Deamidation (NQ) was added as a variable modification together with oxidation (M) and acetyl (N-term). Match-between-runs and re-quantify options were enabled with default parameters and iBAQ values were calculated. Mass spectra were compared to peptide masses from the *Plasmodium falciparum* 3D7 annotated proteome (PlasmoDB v33). The “proteinGroups” file generated by MaxQuant was analysed using the Perseus software package (version 1.4.0.20). The data were filtered against peptides assigned as “only identified by site”, “reverse” and/or “potential contaminant” hits in the datasets. H/L normalized ratios were transformed to log2 values. Significant outliers were identified using the two-sided Benjamini-Hochberg test with an FDR cut-off of 0.01 (Chr3C7 DiQ-BioID experiments 1 and 2) or using boxplot statistics with the cut-off at 1% percentile (Chr3C7 DiQ-BioID experiments 3-6). The H/L normalized ratios of reverse experiments were transformed to -x values to obtain rapalog/control ratios. Finally, scatterplots were produced in R (R i386 3.3.3) by plotting experiment 1 (Chr3C7+BirA*-N^L^) against experiment 2 (Chr3C7+BirA*-C^L^) or forward against reverse experiments (experiment 3 against 4 and 5 against 6, respectively). For the Chr3C16 DiQ-BioID experiment, the significant outliers (cut-off 1% percentile) from H/L normalized ratio (rapalog/control) were plotted using boxplot statistics in GraphPad Prism (version 9).

### Streptavidin-fluorescence assay

To visualise biotinylation in DiQ-BioID experiments, 300 μl of culture from DiQ-BioID experiments were washed twice with 1 x PBS and dried on a 10 well slide followed by fixation in 100% acetone for 30 minutes at room temperature (Spielmann et al., 2003). Cells were re-hydrated in 1 × PBS, blocked in 3% BSA/ 1 × PBS and the primary antibody monoclonal mouse anti-GFP (Roche) at a dilution of 1:1000 in 3% BSA/ 1 × PBS was added for 45 minutes at room temperature. After three wash steps with 1 × PBS, the secondary antibody Alexa Fluor® 488 conjugated anti-mouse (Invitrogen) at a dilution of 1:2000 together with streptavidin coupled to Alexa Fluor® 594 (Invitrogen) at 1:2000 in 3% BSA/ 1 × PBS were applied for 1 hour at room temperature. Unbound antibody and streptavidin were removed by washing three times with 1 × PBS and nuclei were stained with 50 ng/ml Hoechst in 1 × PBS for 10 minutes. For imaging, the slides were mounted with 75% glycerol in 1 × PBS and covered with a cover slip.

### Western Blot analysis

For immunoblots, parasites were harvested, released from infected erythrocytes with 0,03% saponin in 1 x PBS, washed several times in 1 x PBS and lysed in 4% SDS, 0.5% Triton X-100 and 0.5 × PBS with protease inhibitors (Roche). The samples were centrifuged at maximum speed in a tabletop centrifuge and the supernatants were resuspended in Laemmli sample buffer that had been pre-warmed to 80 °C. After separation of proteins using 12% polyacrylamide gels, proteins were transferred to Amersham Protan membranes (GE Healthcare) using a tank blot device (BioRad). Blocking and antibody incubations were performed in 5% skim milk in 1 × TBS supplemented with 0.1% Tween. As primary antibodies, 1:3000 rabbit anti-FKBP (Santa Cruz) and 1:4000 rabbit anti-aldolase (Mesén-Ramírez et al., 2016) were used. 1:3000 diluted HRP-conjugated anti-mouse (Dianova) and 1:4000 diluted HRP-conjugated antirabbit (Dianova) antibodies were used as secondary antibodies. Biotinylated proteins from DiQ-BioID were probed using HRP-conjugated streptavidin (Thermo Fisher) diluted 1:1000 in 5% BSA in 1 × TBS/ 0.1% Tween. Detection was done with ECL (Bio-Rad) and signals were recorded with a ChemiDoc XRS imaging system (Bio-Rad). For quantification of protein levels after diCre-based gene excision in Chr3C30 cKO and Chr3C32 cKO parasites, protein extracts after 24 hours and 44 hours culturing in the presence of 250 nM rapalog and controls were compared on the same immunoblots that were re-probed with aldolase.

Densitometric analyses were performed using the Image Lab software version 5.2.1 (BioRad). The FKBP signal was normalized to the aldolase signal and the ratio of the rapalog-treated extract compared to the control extract that was set to 100%.

### Quantification and statistical analysis

Images of parasites showing localisation, KS and cKO are representative of at least three independent experiments with at least 10 images series per session and condition. All flow cytometry growth curves were performed three times as independent biological replicates (all experiments provided in Figure S2A, D). The second replicates of the two independent biological replicates of detailed stage growth analyses using Giemsa smears are provided in Figure S3F, G; S4A; S6A. Efficiency of di-Cre gene excision of Chr3C30 and Chr3C32 was quantified from three biological replicates shown in Figure S3B. Details about quantification of experiments are given in the respective methods sections. Details about n values, number of replicates and statistical tests used are given in each figure legend if deviating from number indicated here. P values are displayed in the figures. P values >0.05 were considered as not significant. All error bars shown are standard deviations. The assessment of invasion or egress phenotype in Chr3C13 KS parasites was performed in three biological replicates and the examiner was blinded for the identity of samples for counting. Statistical significance was determined using ratio-paired t-test. A twotailed, unpaired t-test was used to compare the parasite area, Hoechst mean fluorescence intensity (MFI) per cell, Hoechst focus area and Hoechst MFI per focus of Chr3C30 cKO parasites after diCre-based excision compared to controls (Figure 3E, G), as well as to compare parasite area, Hoechst MFI per cell, mean nucleus area and number of nuclei in controls to Chr3C32 cKO parasites after gene excision (Figure 3J) as well as for parasite size and number of nuclei in Chr3C16 parasites after KS induction and compared to control (Figure 6C). Parasite size, nuclei size, number of nuclei and DNA content of Chr3C7 and Chr3C16 parasites after KS induction at time points during mitosis assay (Figure 4C, 6G) was compared to controls by using matched repeated-measures one-way ANOVA with assumed sphericity and Bonferroni correction for multiple testing. Statistical analysis was done in GraphPad Prism (version 9).

