## Supplemental information for "Gene-by-gene screen of the unknown proteins encoded on *P. falciparum* chromosome 3"

### Supplementary figures

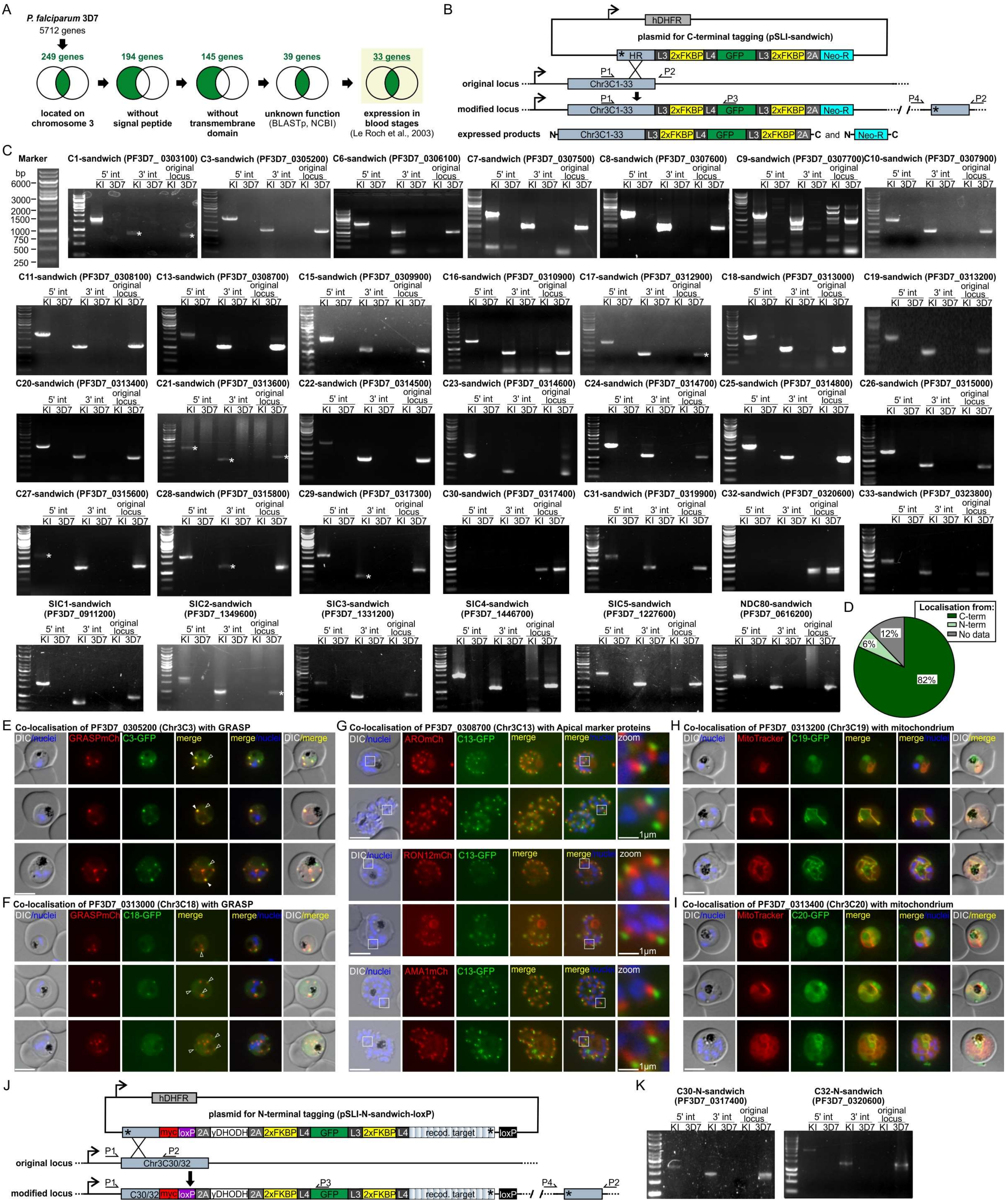


Figure S1: Generation of integrant cell lines and co-localisation analysis of candidates

(A) Schematic of selection strategy. Venn diagrams (black circles) show the selection of genes (green filling, number of genes indicated) that were retained for the next selection step using 'my search strategy' with the indicated parameter in PlasmoDB (v31), resulting in a final set of chosen genes (underlined green number). The step 'unknown function' included all proteins with this annotation in PlasmoDBv31 where in addition all genes encoding proteins with homology outside the Apicomplexa (identified using manual BLAST searches, e value < 1e-50 and percent identity ≥ 55 %) were subtracted. In the last step, all 33 genes that are expressed during erythrocytic stages were chosen (Le Roch et al., 2003, expression level ≥ 13.7)

(B) Integration strategy using SLI to obtain endogenously 2xFKBP-GFP-2xFKBP (C-terminal) tagged Chr3C1-33, SICs and NDC80. Small arrows show position of primers used for assessing correct integration of the plasmid into the genome shown in (C). L3 and L4 are linkers, asterisks indicate stop codons and 2A denotes the skip peptide, corresponding to the features of pSLI-sandwich (Birnbaum et al., 2017).

(C) Agarose gels showing PCR products amplified from genomic DNA of the indicated knock-in (KI) parasite lines compared to 3D7 wildtype parasites (3D7) to confirm SLI integration into the correct gene locus. Primer combinations (indicated in B) produced products across the 5' (P1 and P3) and 3' (P2 and P4) integration junctions ('5'int' and '3'int' lanes, respectively) and absence of original locus (P1 and P2). 'Sandwich' indicates C-terminal tagging with 2xFKBP-GFP-2xFKBP (Birnbaum et al., 2017). Note that for C30-sandwich and C32-sandwich no C-terminal integration was achieved. White asterisks indicate poorly visible bands. Marker fragment length is indicated in base pairs at the top left.

(D) Pie chart displaying proportion of candidates that were C-terminally ('C-term') or N-terminally ('N-term') tagged. Chr3C2, Chr3C4, Chr3C5, Chr3C14 were refractory to C-terminal tagging and were not N-terminally tagged ('no data').

(E and F) Representative live cell fluorescence images showing parasites with endogenously GFP-tagged Chr3C3 (E) or Chr3C18 (F) co-expressing GRASP-mCherry as a marker protein for Golgi. Filled white arrows show co-localisation of GFP-foci with GRASP-mCherry foci. Non-filled arrows indicate no co-localisation.

(G) Representative live cell fluorescence images showing parasites with endogenously GFP-tagged Chr3C13 co-expressing either ARO-mCherry, RON12-mCherry or AMA1-mCherry. White boxes show enlarged regions ('zoom').

(H and I) Representative live cell fluorescence images showing parasites with endogenously GFP-tagged Chr3C19 (H) or Chr3C20 (I) and stained with MitoTracker Red for co-localisation with mitochondrion.

(J) Integration strategy using SLI to obtain endogenously 2xFKBP-GFP-2xFKBP (N-terminal) tagged Chr3C30 and Chr3C32. The functional copy of the gene was recodonised ('recod. target'). Small arrows show position of primers used for testing correct integration into the genome (C). L3 and L4 are linkers, asterisks indicate stop codons and 2A denotes the skip peptide, corresponding to the features of pSLI-N-sandwich (Birnbaum et al., 2017).

(K) Agarose gels showing PCR products amplified from genomic DNA of Chr3C30 and Chr3C32 N-terminal knock-in (KI) parasite line compared to 3D7 wildtype parasites (3D7) to confirm SLI integration into the correct gene locus. Primer combinations (indicated in J) produced products across the 5' (P1 and P3) and 3' (P2 and P4) integration junctions ('5'int' and '3'int' lanes, respectively) and absence of original locus (P1 and P2). 'N-Sandwich' indicates N-terminally tagging with 2xFKBP-GFP-2xFKBP (Birnbaum et al., 2017). Marker fragment length is indicated in base pairs at the top left of C.

*Scale bars, 5 µm (if not indicated otherwise); DIC, differential interference contrast; merge, merged green and red channel; mCh, mCherry; nuclei stained with Hoechst 33342.*

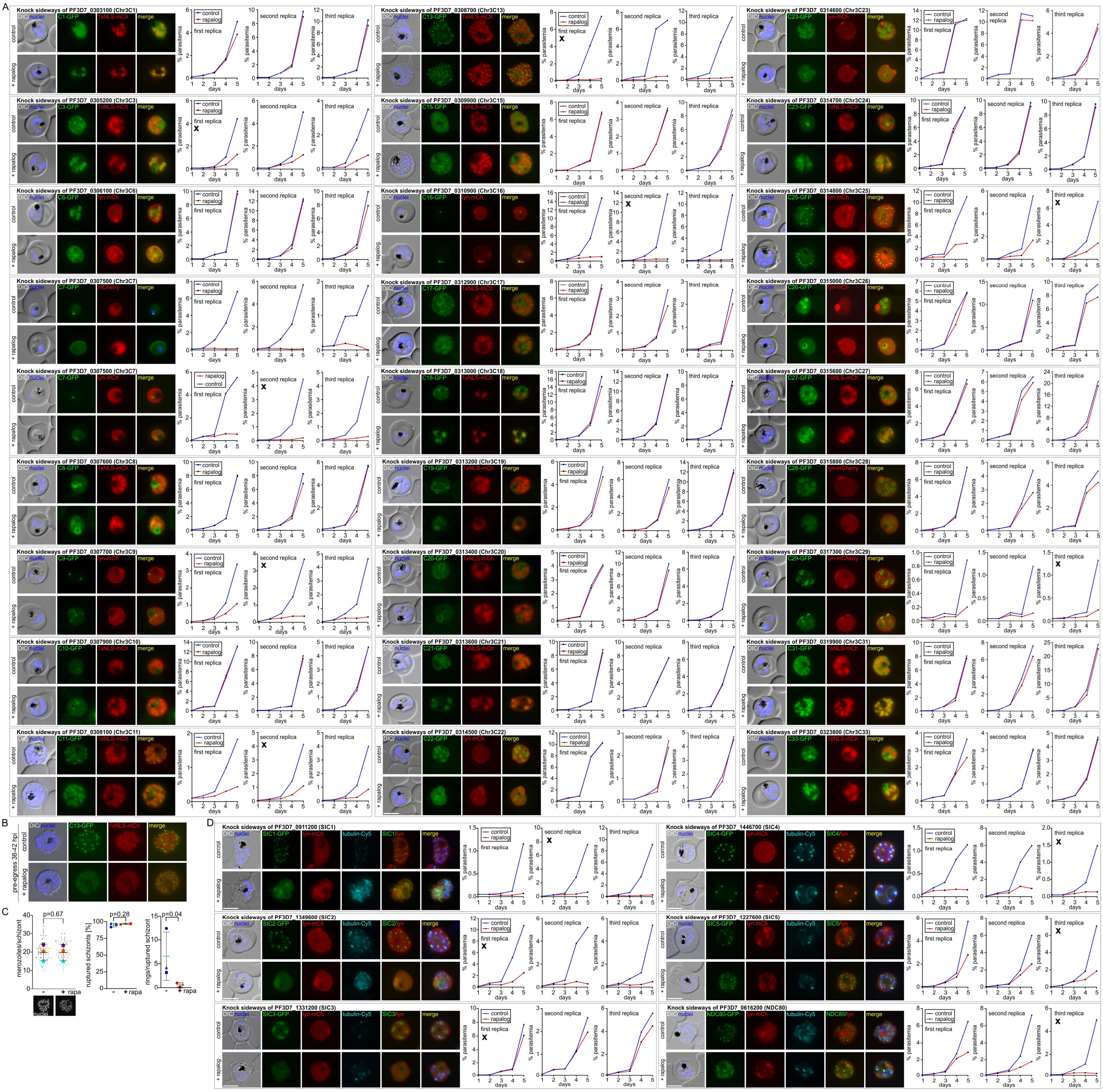


<u>Figure S2: Analysis of essentiality of all candidates in this study</u>

(A and D) Full panels of KS with knock-in parasite line of the indicated chromosome 3 candidates (A) or SICs and NDC80 (D) episomally expressing either the lyn-FRB-mCherry (lyn-mCh) for mislocalisation to the parasite plasma membrane or 1xNLS-FRB-mCherry mislocaliser (1xNLS-mCh) for mislocalisation into the nucleus grown in absence (control) or presence of rapalog (+rapalog, induced KS) shown next to all replicates of flow cytometry growth curves. Images (17-24h after addition of rapalog) are representatives of at least 10 image series per experiment and condition). Where applicable, the experiment shown in Figure 2 (A) or Figure 5 (D) is marked with an ‘x’ symbol.

(B) Representative live cell fluorescence images showing synchronised (0-4 h synchronisation window) Chr3C13 KS late schizonts at the ‘pre-egress’ time point (38-42 hpi) grown in absence (control) and presence (+ rapalog, induced Chr3C13 KS) of rapalog.

(C) Quantification of merozoites per schizont at ‘pre-egress’ time point (left; example pictures of Hoechst-stained nuclei below the graph), ruptured schizonts (middle) and rings per ruptured schizont (right) at ‘post-egress’ time point in Chr3C13 KS schizonts after KS (+rapa) compared to control (-rapa) from n=3 independent experiments. Unpaired ratio-paired t-test, P values are indicated, error bars are SD. *Scale bars, 5 µm; DIC, differential interference contrast; merge, merged green and red channel; mCh, mCherry; nuclei stained with DAPI or Hoechst 33342; tubulin-Cy5; stained with Tubulin Tracker Deep Red; hpi, hours post invasion.*

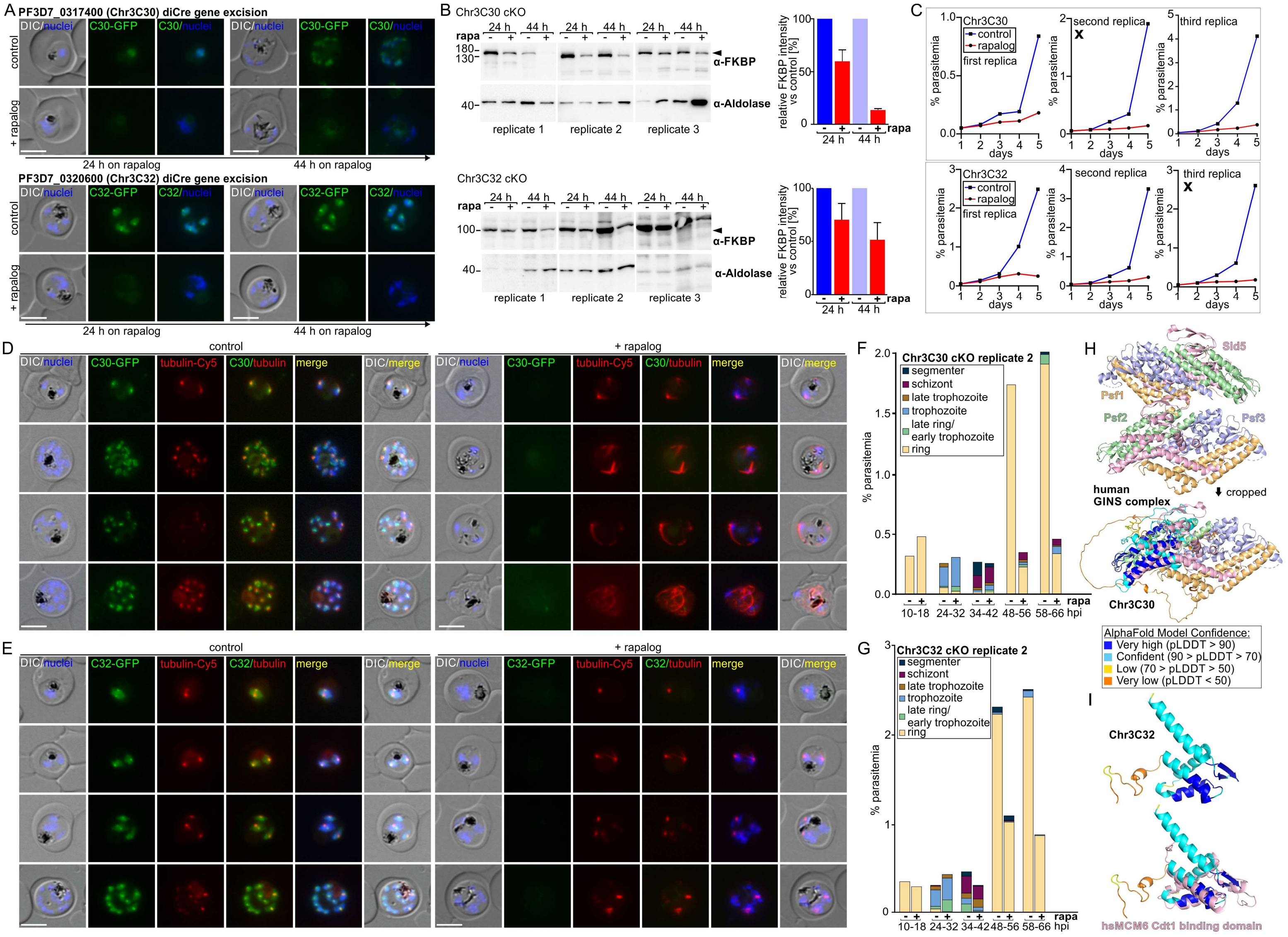


Figure S3: Phenotype analysis of Chr3C30 and Chr3C32 cKO

(A) Live cell fluorescence microscopy images (representatives of 3 independent microscopy sessions with each at least 10 image series per session and condition) showing synchronised parasites (0-8 h synchronisation window) with N-terminally GFP-tagged Chr3C30 (top) or Chr3C32 (bottom) episomally expressing diCre (from pSkipFlox) grown in absence (control) and presence (+rapalog, induced diCre based gene excision) of rapalog for 24 h (left panel) or 44 h (right panel).

(B) Western blots (left) probed with anti-FKBP antibodies to detect N-terminally 2xFKBP-GFP-2xFKBP tagged C30 (top) or C32 (bottom) from cKO parasites grown in presence (+rapa, induced diCre based gene excision) and absence (- rapa) of rapalog for 24 h or 44 h. Anti-aldolase antibodies were used as loading control. Levels of 2xFKBP-GFP-2xFKBP-C30 (top) or 2xFKBP-GFP-2xFKBP-C32 (bottom) 24 h and 44 h after addition of rapalog were quantified (right) based on densitometric analysis of anti-FKPB western blots (mean of n=3 independent experiments) shown left. FKBP signal was normalised to aldolase. Error bars indicate SD.

(C) Flow cytometry growth curves over 5 days of 3 independent experiments for Chr3C30 cKO (top) and Chr3C32 cKO (bottom). The experiments shown in Figure 2E are marked with an 'x' symbol.

(D and E) Live cell fluorescence microscopy images (representatives of 3 independent microscopy sessions with each at least 10 image series per session) of Chr3C30 cKO (D) or Chr3C32 cKO (E) parasites grown in absence (control) and presence of rapalog (+rapalog, induced diCre based gene excision) and co-stained with Tubulin Tracker Deep Red.

(F and G) Second biological replicate (first replicates in Figure 3C, D) of stages and growth based on Giemsa smears in synchronised (0-8 h synchronisation window) in Chr3C30 cKO (F) or Chr3C32 cKO (G) parasites grown in presence (+rapa, induced diCre gene excision) and absence (-rapa) of rapalog at the indicated time points.

(H) Top, full protein structure of human GINS complex (2Q9Q) that was cropped into monomer for Figure 3I. Bottom, alignment of Chr3C30 structure predicted by alphafold (color coded according to alphafold prediction confidence as indicated in legend) with human GINS monomer structure (RMSD=2.8 over 168 residues).

(I) Chr3C32 protein structure prediction by alphafold shown in Figure 3L (colour coded according to alphafold prediction confidence as indicated in legend in H) and alignment with human MCM6 Cdt1 binding domain (2KLQ; RMSD=5.3 over 88 residues).

*Scale bars, 5 µm; DIC, differential interference contrast; merge, merged green, red, and blue channel; mCh, mCherry; nuclei stained with Hoechst 33342; tubulin-Cy5; stained with Tubulin Tracker Deep Red; hpi, hours post invasion.*

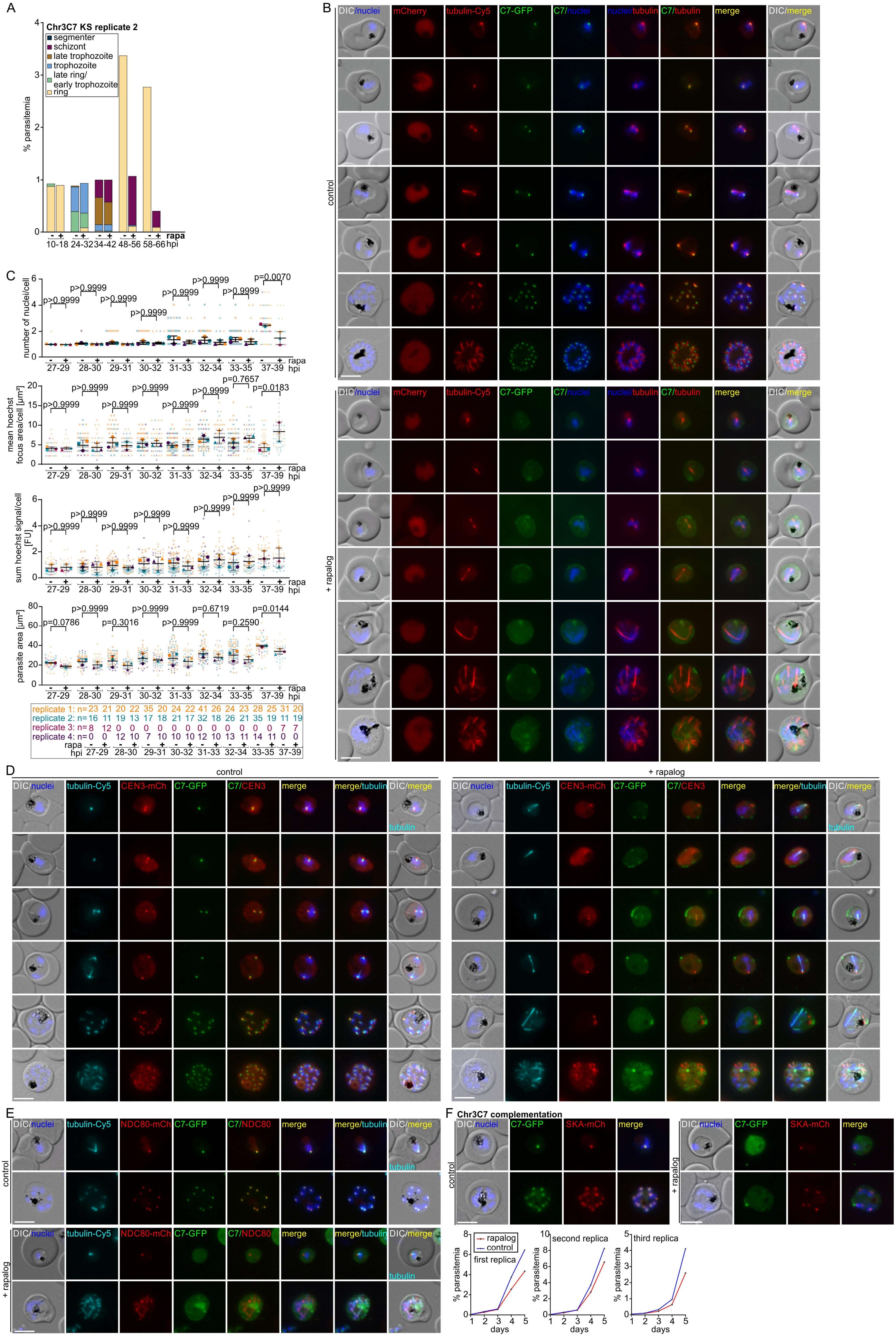
A
Chr3C7 KS replicate 2
segmenter
schizont
late trophozoite
trophozoite
late ring/ early trophozoite
ring
% parasitemia
rapa
10-18 24-32 34-42 48-56 58-66 hpi
B
DIC/nuclei
mCherry
tubulin-Cy5
C7-GFP
C7/nuclei
nuclei/tubulin
C7/tubulin
merge
DIC/merge
control
+ rapalog
C
number of nuclei/cell
mean hoechst focus area/cell [µm²]
sum hoechst signal/cell [FU]
parasite area [µm²]
p>0.9999
p=0.0070
p=0.7657
p=0.0183
p=0.0786
p=0.3016
p=0.6719
p=0.2590
p=0.0144
27-29 28-30 29-31 30-32 31-33 32-34 33-35 37-39 hpi
replicate 1: n=23 21 20 22 35 20 24 22 41 26 24 23 28 25 31 20
replicate 2: n=16 11 19 13 17 18 21 17 32 18 26 21 35 19 11 19
replicate 3: n=8 12 0 0 0 0 0 0 0 0 0 0 0 0 7 7
replicate 4: n=0 0 12 10 7 10 10 10 12 10 13 11 14 11 0 0
rapa
hpi 27-29 28-30 29-31 30-32 31-33 32-34 33-35 37-39
D
control
+ rapalog
DIC/nuclei
tubulin-Cy5
CEN3-mCh
C7-GFP
C7/CEN3
merge
merge/tubulin
DIC/merge
tubulin
E
DIC/nuclei
tubulin-Cy5
NDC80-mCh
C7-GFP
C7/NDC80
merge
merge/tubulin
DIC/merge
tubulin
control
+ rapalog
F
Chr3C7 complementation
DIC/nuclei
C7-GFP
SKA-mCh
merge
control
+ rapalog
rapalog
control
first replica
second replica
third replica
% parasitemia
days

Figure S4: Functional analysis of Chr3C7

(A) Second biological replicate (first replicates in Figure 4A) of stages and growth based on Giemsa smears in synchronised (0-8 h synchronisation window) in Chr3C7 KS parasites grown in presence (+rapa, to induce KS) and absence (- rapa) of rapalog at the indicated time points.

(B) Superplots showing quantification of nuclei number, nucleus size, DNA content and parasite size in synchronised (0-2 h synchronisation window) Chr3C7 parasites after KS induction (+ rapa) compared to control (- rapa) at all time points of the experiment shown in Figure 4C (n=4 independent experiments). Multiple paired t-tests (matched repeated-measures one-way ANOVA with assumed sphericity and Bonferroni correction for multiple testing); P values indicated; error bars are SD; n numbers of scored parasites at the indicated time points in the individual experiments are shown on the right.

(C) Full panels of live cell fluorescence microscopy images shown in Figure 4B, D, F (representative images of 3 independent microscopy sessions with at least 10 image series per session and condition) of Chr3C7 KS parasites co-stained with Tubulin Tracker Deep Red and grown in absence (control) and presence (+ rapalog, induced Chr3C7 KS) of rapalog. Presence of modified lyn-mislocaliser (lyn-FRB-T2A-mCherry (Hoeijmakers et al., 2019)) shown in 'mCherry' image column.

(D and E) Full panels of live cell fluorescence microscopy images shown in Figure 4E, G, H (representative images of 3 independent microscopy sessions with at least 10 image series per session and condition) of Chr3C7 KS parasites co-expressing the marker proteins CEN3-mCherry (D) and NDC80-mCherry (E) for co-localisation during schizogony and co-stained with Tubulin Tracker Deep Red. Parasites were grown in absence (control) and presence (+ rapalog, induced Chr3C7 KS) of rapalog.

(F) Complementation of Chr3C7 KS. Representative live cell fluorescent images (top; 3 independent microscopy sessions with at least 10 image series per session and condition) and flow cytometry growth curves showing parasitemia over 5 days from 3 independent experiments (bottom) of endogenously GFP-tagged Chr3C7 parasites co-expressing an episomal copy of Chr3C7 (SKA-mCherry) and the lyn-mislocaliser at the same time for complementation of Chr3C7 KS. Parasites grown in absence (control) and presence (+rapalog, induced KS of endogenously tagged Chr3C7) of rapalog.

*Scale bars, 5 µm; DIC, differential interference contrast; merge, merged green, red, and blue channel; mCh, mCherry; nuclei stained with Hoechst 33342; tubulin-Cy5; stained with Tubulin Tracker Deep Red; hpi, hours post invasion; FU, arbitrary fluorescence units.*

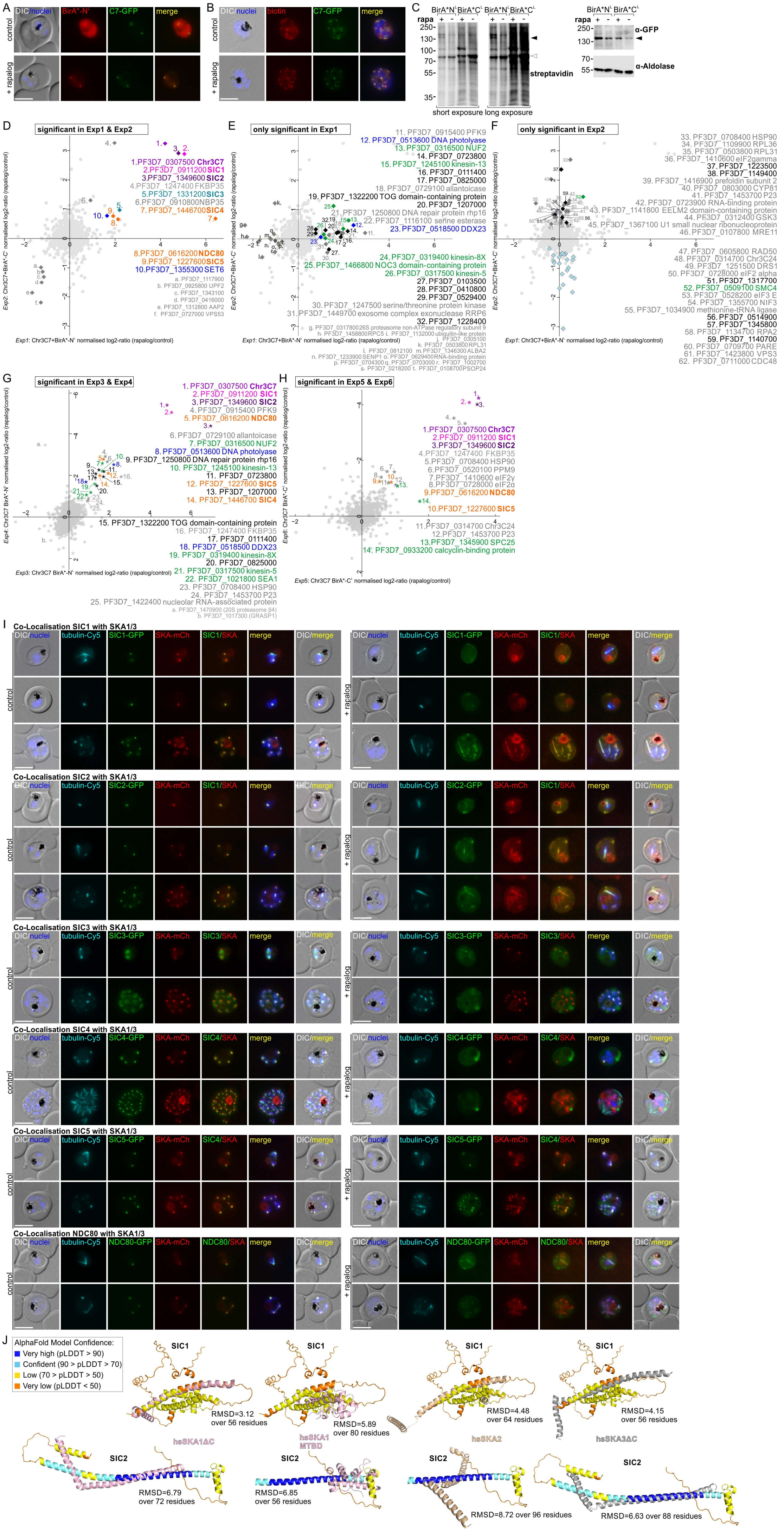
A
DIC/nuclei
BirA*-N^L
C7-GFP
merge
control
+ rapalog
B
DIC/nuclei
biotin
C7-GFP
merge
C
BirA*N^L BirA*C^L
rapa
short exposure long exposure
streptavidin
α-GFP
α-Aldolase
D
significant in Exp1 & Exp2
1.PF3D7_0307500 Chr3C7
2.PF3D7_0911200 SIC1
3.PF3D7_1349600 SIC2
4.PF3D7_1247400 FKBP35
5.PF3D7_1331200 SIC3
6.PF3D7_0910800 NBP35
7.PF3D7_1446700 SIC4
8.PF3D7_0616200 NDC80
9.PF3D7_1227600 SIC5
10.PF3D7_1355300 SET6
a. PF3D7_1117900
b. PF3D7_0925800 UPF2
c. PF3D7_1343100
d. PF3D7_0416000
e. PF3D7_1312800 AAP2
f. PF3D7_0727000 VPS53
Exp1: Chr3C7+BirA*-N^L normalised log2-ratio (rapalog/control)
Exp2: Chr3C7+BirA*-C^L normalised log2-ratio (rapalog/control)
E
only significant in Exp1
11. PF3D7_0915400 PFK9
12. PF3D7_0513600 DNA photolyase
13. PF3D7_0316500 NUF2
14. PF3D7_0723800
15. PF3D7_1245100 kinesin-13
16. PF3D7_0111400
17. PF3D7_0825000
18. PF3D7_0729100 allantoicase
19. PF3D7_1322200 TOG domain-containing protein
20. PF3D7_1207000
21. PF3D7_1250800 DNA repair protein rhp16
22. PF3D7_1116100 serine esterase
23. PF3D7_0518500 DDX23
24. PF3D7_0319400 kinesin-8X
25. PF3D7_1466800 NOC3 domain-containing protein
26. PF3D7_0317500 kinesin-5
27. PF3D7_0103500
28. PF3D7_0410800
29. PF3D7_0529400
30. PF3D7_1247500 serine/threonine protein kinase
31. PF3D7_1449700 exosome complex exonuclease RRP6
32. PF3D7_1228400
F
only significant in Exp2
33. PF3D7_0708400 HSP90
34. PF3D7_1109900 RPL36
35. PF3D7_0503800 RPL31
36. PF3D7_1410600 eIF2gamma
37. PF3D7_1223500
38. PF3D7_1149400
39. PF3D7_1416900 prefoldin subunit 2
40. PF3D7_0803000 CYP81
41. PF3D7_1453700 P23
42. PF3D7_0723900 RNA-binding protein
43. PF3D7_1141800 EELM2 domain-containing protein
44. PF3D7_0312400 GSK3
45. PF3D7_1367100 U1 small nuclear ribonucleoprotein
46. PF3D7_0107800 MRE11
47. PF3D7_0605800 RAD50
48. PF3D7_0314700 Chr3C24
49. PF3D7_1251500 DRS1
50. PF3D7_0728000 eIF2 alpha
51. PF3D7_1317700
52. PF3D7_0509100 SMC4
53. PF3D7_0528200 eIF3 E
54. PF3D7_1355300 NIF3
55. PF3D7_1034900 methionine-tRNA ligase
56. PF3D7_0514900
57. PF3D7_1345800
58. PF3D7_1134700 RPA2
59. PF3D7_1140700
60. PF3D7_0709700 PARE
61. PF3D7_1423800 VPS3
62. PF3D7_0711000 CDC48
G
significant in Exp3 & Exp4
1. PF3D7_0307500 Chr3C7
2. PF3D7_0911200 SIC1
3. PF3D7_1349600 SIC2
4. PF3D7_0915400 PFK9
5. PF3D7_0616200 NDC80
6. PF3D7_0729100 allantoicase
7. PF3D7_0316500 NUF2
8. PF3D7_0513600 DNA photolyase
9. PF3D7_1250800 DNA repair protein rhp16
10. PF3D7_1245100 kinesin-13
11. PF3D7_0723800
12. PF3D7_1227600 SIC5
13. PF3D7_1207000
14. PF3D7_1446700 SIC4
15. PF3D7_1322200 TOG domain-containing protein
16. PF3D7_1247400 FKBP35
17. PF3D7_0111400
18. PF3D7_0518500 DDX23
19. PF3D7_0319400 kinesin-8X
20. PF3D7_0825000
21. PF3D7_0317500 kinesin-5
22. PF3D7_1021800 SEA1
23. PF3D7_0708400 HSP90
24. PF3D7_1453700 P23
25. PF3D7_1422400 nucleolar RNA-associated protein
a. PF3D7_1470900 (20S proteasome β4)
b. PF3D7_1017300 (GRASP1)
Exp3: Chr3C7 BirA*-N^L normalised log2-ratio (rapalog/control)
Exp4: Chr3C7 BirA*-N^L normalised log2-ratio (rapalog/control)
H
significant in Exp5 & Exp6
1.PF3D7_0307500 Chr3C7
2.PF3D7_0911200 SIC1
3.PF3D7_1349600 SIC2
4.PF3D7_1247400 FKBP35
5.PF3D7_0708400 HSP90
6.PF3D7_0520100 PPM9
7.PF3D7_1410600 eIF2γ
8.PF3D7_0728000 eIF2α
9.PF3D7_0616200 NDC80
10.PF3D7_1227600 SIC5
11.PF3D7_0314700 Chr3C24
12.PF3D7_1453700 P23
13.PF3D7_1345900 SPC25
14. PF3D7_0933200 calcyclin-binding protein
Exp5: Chr3C7 BirA*-C^L normalised log2-ratio (rapalog/control)
Exp6: Chr3C7 BirA*-C^L normalised log2-ratio (rapalog/control)
I
Co-Localisation SIC1 with SKA1/3
Co-Localisation SIC2 with SKA1/3
Co-Localisation SIC3 with SKA1/3
Co-Localisation SIC4 with SKA1/3
Co-Localisation SIC5 with SKA1/3
Co-Localisation NDC80 with SKA1/3
DIC/nuclei
tubulin-Cy5
SKA-mCh
merge
DIC/merge
control
+ rapalog
J
AlphaFold Model Confidence:
Very high (pLDDT > 90)
Confident (90 > pLDDT > 70)
Low (70 > pLDDT > 50)
Very low (pLDDT < 50)
SIC1
SIC2
RMSD=3.12 over 56 residues
RMSD=5.89 over 80 residues
RMSD=4.48 over 64 residues
RMSD=4.15 over 56 residues
hsSKA1ΔC
hsSKA1 MTBD
hsSKA2
hsSKA3ΔC
RMSD=6.79 over 72 residues
RMSD=6.85 over 56 residues
RMSD=8.72 over 96 residues
RMSD=6.63 over 88 residues

Figure S5: DiQ-BioID with Chr3C7 and validation of SKA interacting candidates

(A) Representative live cell images showing recruitment of the episomally expressed biotinyliser (BirA*-$N^L$) to endogenously 2xFKBP-GFP-2xFKBP tagged Chr3C7 after addition of rapalog (+rapalog) compared to control.

(B) Streptavidin-fluorescence assay images of acetone fixed parasites showing biotinylation of proteins (streptavidin coupled to Alexa Fluor® 594, red) at location of endogenously 2xFKBP-GFP-2xFKBP tagged Chr3C7 (mouse anti-GFP, green) after adding rapalog (+rapalog, induced recruitment of biotinyliser) compared to control.

(C) Streptavidin blot (left) showing biotinylation in Chr3C7 BirA*-$N^L$ and BirA*-$C^L$ parasites cultured in the presence (+ rapa, induced recruitment of biotinyliser) and absence (- rapa) of rapalog with biotin supplementation. Long exposure shows multiple background bands, indicating biotinylation of other protein. White arrow indicates self-biotinylation of biotinyliser. Black arrow indicates biotinylation at the target (Chr3C7), as evident from the band at similar height shown on anti-GFP blot (right). Anti-Aldolase antibodies were used as loading control.

(D-H) Full scatters plots of DiQ-BioID experiments showing proteins significantly enriched (FDR<0.01) in both experiment 1 and 2 using BirA*-$N^L$ and BirA*-$C^L$, respectively (D) or in only one of the experiments (E and F). For experiment 3 and 4 (G) the BirA*-$N^L$ biotinyliser and for experiment 5 and 6 (H) BirA*-$C^L$ was used, respectively and proteins significantly enriched in both the indicated experiments are shown in the scatter plots. Enriched proteins are numbered according to their enrichment score and colour coded (for colour code, see Figure 5): for SICs and NDC80 (orange) known mitosis proteins (green), nuclear proteins (blue), unknowns (black) and common contaminants (grey), depleted proteins are lettered. PlasmoDB identifiers and annotations are given.

(I) Full panels of live cell fluorescence microscopy images shown in Figure 5D, F (representatives of 3 independent microscopy sessions with at least 10 image series per session and condition), showing co-localisation of SKA1/3 in SIC1-5 and NDC80 KS late schizonts grown in absence (control) and presence (+rapalog, induced KS) of rapalog and co-stained with Tubulin Tracker Deep Red.

(J) Alignment of SIC1 (top) and SIC2 (bottom) structure predicted by alphafold (shown in alphafold confidence colours) with structure of SKA domain of hsSKA1ΔC (4AJ5), microtubule-binding domain (MTBD) of hsSKA1 (4C9Y), hsSKA2 (4AJ5) and hsSKA3ΔC (4AJ5). Alignment scores (RMSD) indicated below the structures.

*Size bars, 5 µm; DIC, differential interference contrast; tubulin, stained with Tubulin Tracker Deep Red; nuclei stained with Hoechst 33342; merge, merged GFP (green), mCherry (red), Cy5 (cyan) and Hoechst (blue) channels.*

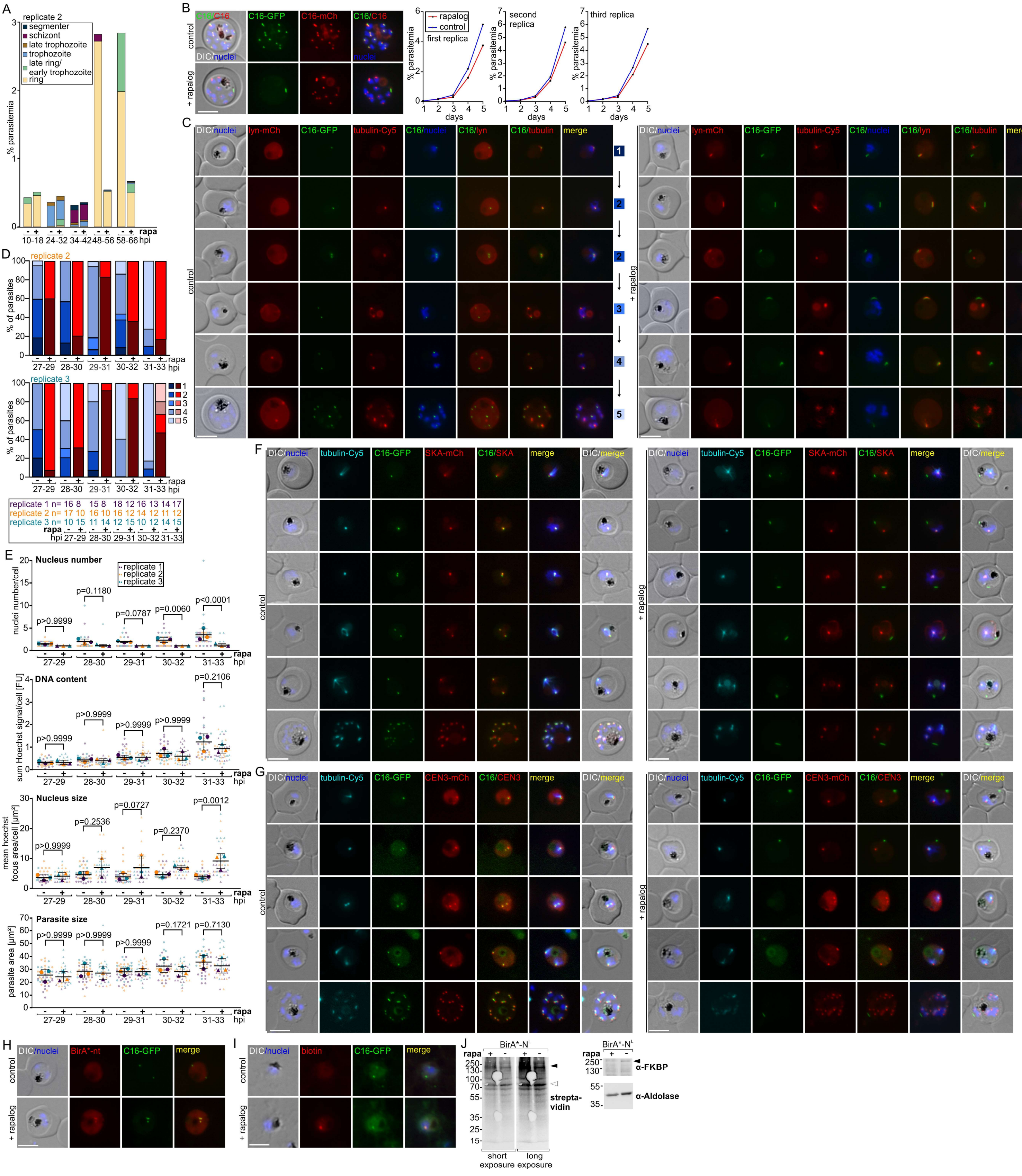


Figure S6: Functional analysis and DiQ-BioID of Chr3C16

(A) Second biological replicate (first replicate in Figure 6A) of stages and growth based on Giemsa smears in synchronised (0-8 h synchronisation window) in Chr3C16 KS parasites grown in presence (+ rapa, induced Chr3C16 KS) and absence of rapalog (- rapa) at the indicated time points.

(B) Complementation of Chr3C16 KS. Representative live cell fluorescent images (top; 3 independent microscopy sessions with at least 10 image series per session and condition) and flow cytometry growth curves showing parasitemia over 5 days from 3 independent experiments (bottom) of endogenously GFP-tagged Chr3C16 parasites co-expressing an episomal copy of Chr3C16 and the lyn-mislocaliser at the same time for complementation of Chr3C16 KS. Parasites were grown in absence (control) and presence (+ rapalog, induced Chr3C16 KS) of rapalog.

(C) Full panels of live cell fluorescence microscopy images from Figure 6B, D, E (representative images of 3 independent microscopy sessions with at least 10 image series per session and condition) of Chr3C16 KS parasites co-stained with Tubulin Tracker Deep Red and grown in absence (control) and presence (+ rapalog, induced Chr3C16 KS) of rapalog. Presence of lyn-mislocaliser is shown in 'lyn-mCh' image column.

(D) Second and third replicate of quantification of the experiment shown in Figure 6F of synchronised Chr3C16 KS parasites (0-2 h synchronisation window) grown with (+ rapalog, induced Chr3C16 KS) or without rapalog (- rapalog) showing the 5 phases defined in (C) at the indicated time points during the first nuclear division. Stage 5 parasites were defined as all parasites with more than two nuclei or with two nuclei where a new round of nuclear division had been started. n numbers of scored parasites at the indicated timepoints from the individual experiments are shown below the graph.

(E) Superplots showing quantification of nuclei number, nucleus size, DNA content and parasite size in synchronised (0-2 h synchronisation window) Chr3C16 parasites after KS (+ rapa) compared to control (- rapa) at all time points of the experiment shown in Figure 6G (n=3 independent experiments). Multiple paired t-tests (matched repeated-measures one-way ANOVA with assumed sphericity and Bonferroni correction for multiple testing); P values indicated; error bars are SD; n numbers of scored parasites at the indicated timepoints from the individual experiments are shown in D.

(F and G) Full panels of live cell fluorescence microscopy images from Figure 6H-M (representative images of 3 independent microscopy sessions with at least 10 image series per session and condition) of Chr3C16 KS parasites co-expressing the marker proteins SKA1/3-mCherry (F) and CEN3-mCherry (G) for co-localisation during schizogony and co-stained with Tubulin Tracker Deep Red. Parasites were grown in absence (control) and presence (+ rapalog, induced Chr3C16 KS) of rapalog.

(H) Representative live cell images showing recruitment of the episomally expressed biotinyliser (BirA*-$N^L$) to endogenously 2xFKBP-GFP-2xFKBP tagged Chr3C16 after addition of rapalog (+ rapalog) compared to control.

(I) Streptavidin-fluorescence assay images of acetone fixed parasites showing biotinylation of proteins (streptavidin coupled to Alexa Fluor® 594, red) at location of endogenously 2xFKBP-GFP-2xFKBP tagged Chr3C16 (mouse anti-GFP, green) after adding rapalog (+ rapalog, induced recruitment of the biotinyliser) compared to control from the experiment shown in Figure 6N.

(J) Streptavidin blot (left) showing biotinylation in Chr3C16 BirA* $N^L$ parasites cultured in the presence (- rapa) and absence (+ rapa, induced recruitment of biotinyliser) of rapalog with biotin supplementation from the experiment shown in Figure 6N. Long exposure shows multiple background bands, indicating biotinylation of other proteins. White arrow indicates self-biotinylation of biotinyliser. Black arrow indicates biotinylation at the target (Chr3C16), as evident from the band at similar height shown on anti-FKBP blot (right). Anti-aldolase antibodies were used as loading control.

*Size bars, 5 µm; DIC, differential interference contrast; tubulin, stained with Tubulin Tracker Deep Red; nuclei stained with Hoechst 33342; merge, merged green, red and blue channels; hpi, hours post invasion; FU, arbitrary fluorescence units.*

### Tables

Table S1: Detailed information about candidates

Table S2: Cell lines, recombinant DNA, and oligonucleotides

Table S3: Mass spectrometry results of Chr3C7 DiQ-BioID and Chr3C16 DiQ-BioID

### Datafiles

Datafile 1: Subcellular localisation of all tagged candidates in this study

Full panels of live cell fluorescence microscopy images showing the location of GFP-tagged candidates, SICs and NDC80 expressed from the endogenous locus (representative images of 3 independent microscopy sessions with at least 10 image series each) during asexual blood stages (ring, early trophozoite, late trophozoite, early schizont, late schizont, segmenter stage parasites). Free merozoites of GFP-tagged Chr3C13 are also shown. SICs and NDC80 parasites were stained with Tubulin Tracker Deep Red for co-localisation with microtubules.
