## Supplementary material for "Gene-by-gene screen of the unknown proteins encoded on *P. falciparum* chromosome 3": Datafile 1

**Localisation of PF3D7_0303100 (Chr3C1)**

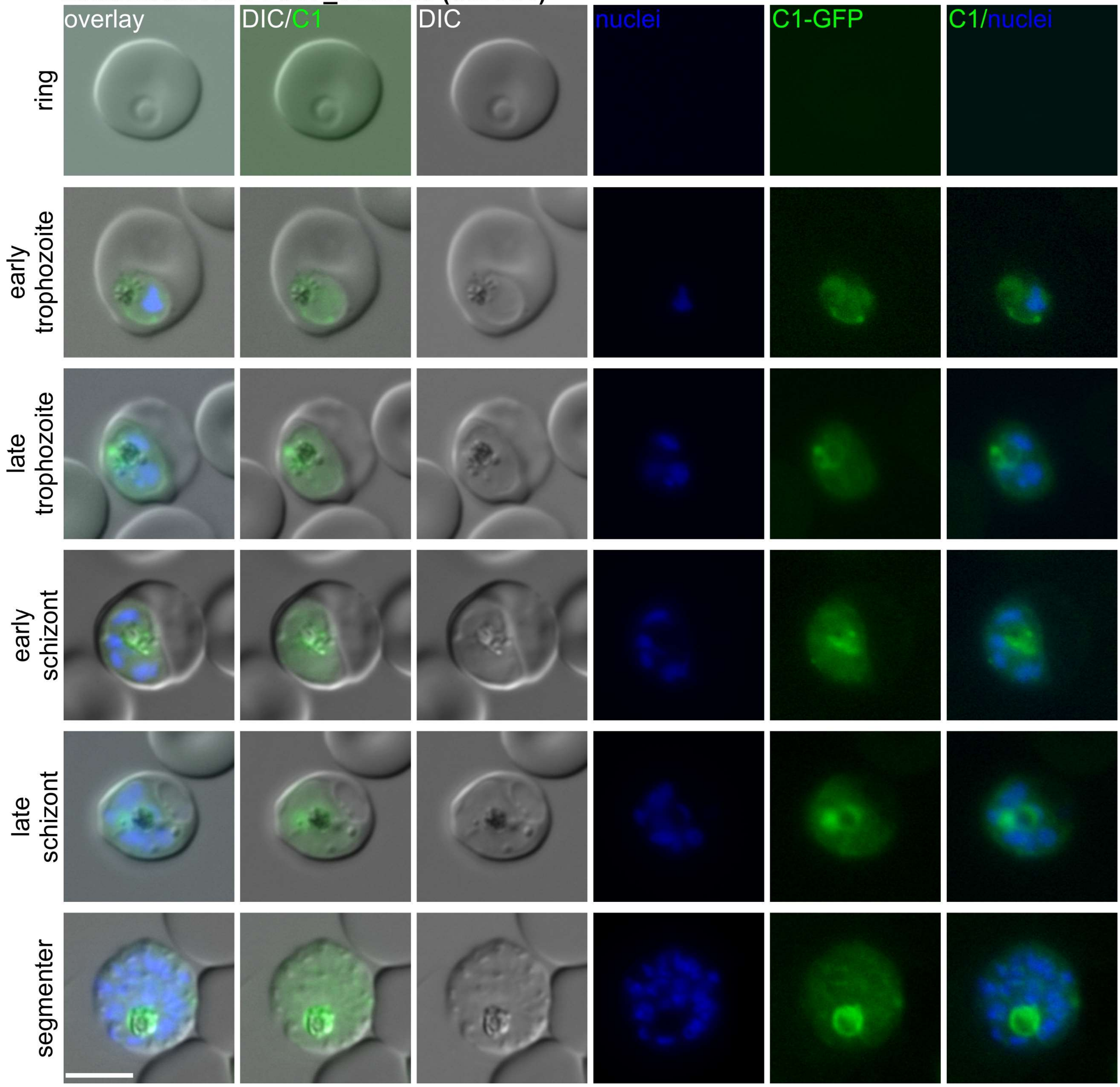

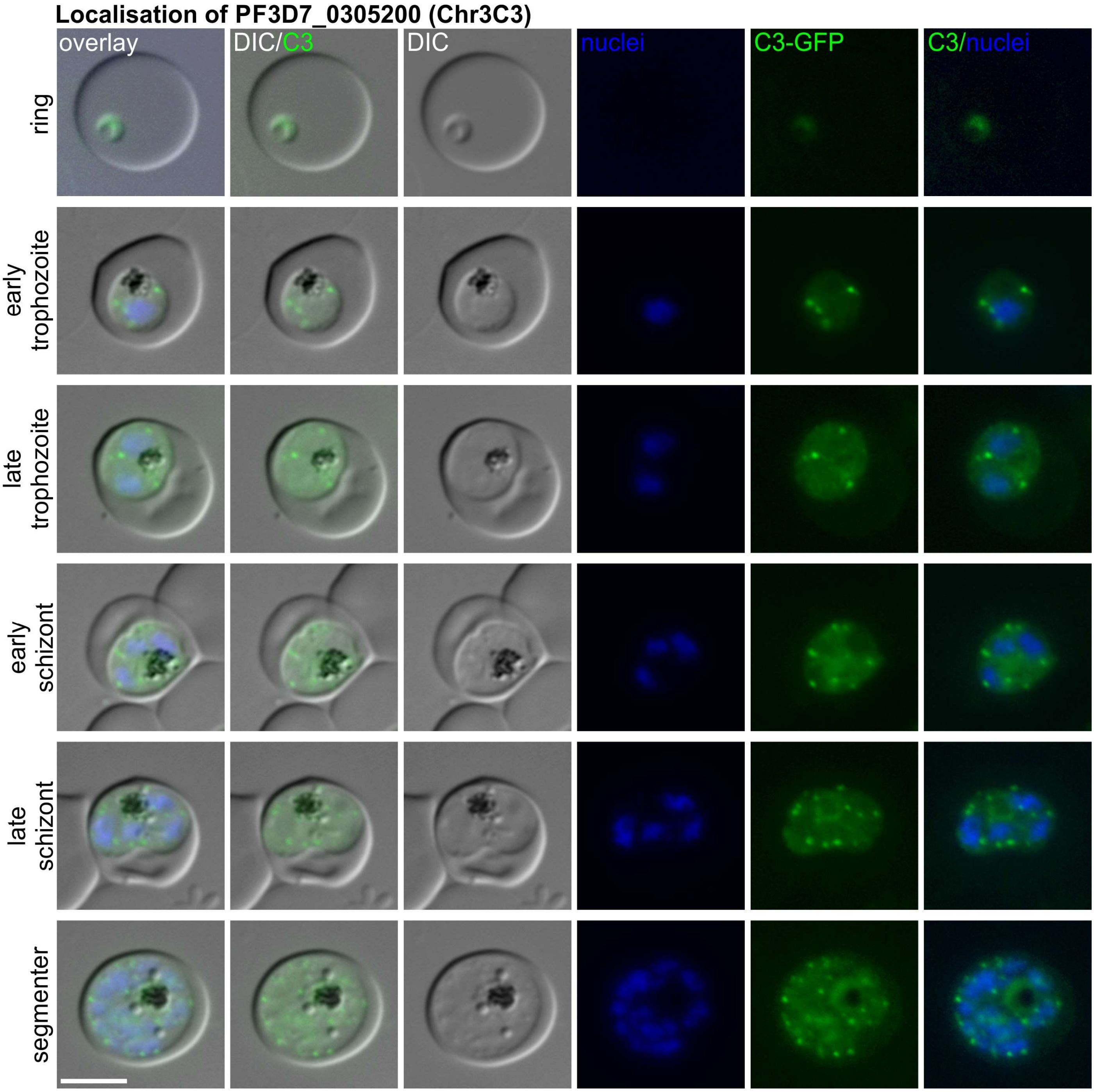
Localisation of PF3D7_0305200 (Chr3C3)
overlay
DIC/C3
DIC
nuclei
C3-GFP
C3/nuclei
ring
early trophozoite
late trophozoite
early schizont
late schizont
segmenter

**Localisation of PF3D7_0306100 (Chr3C6)**

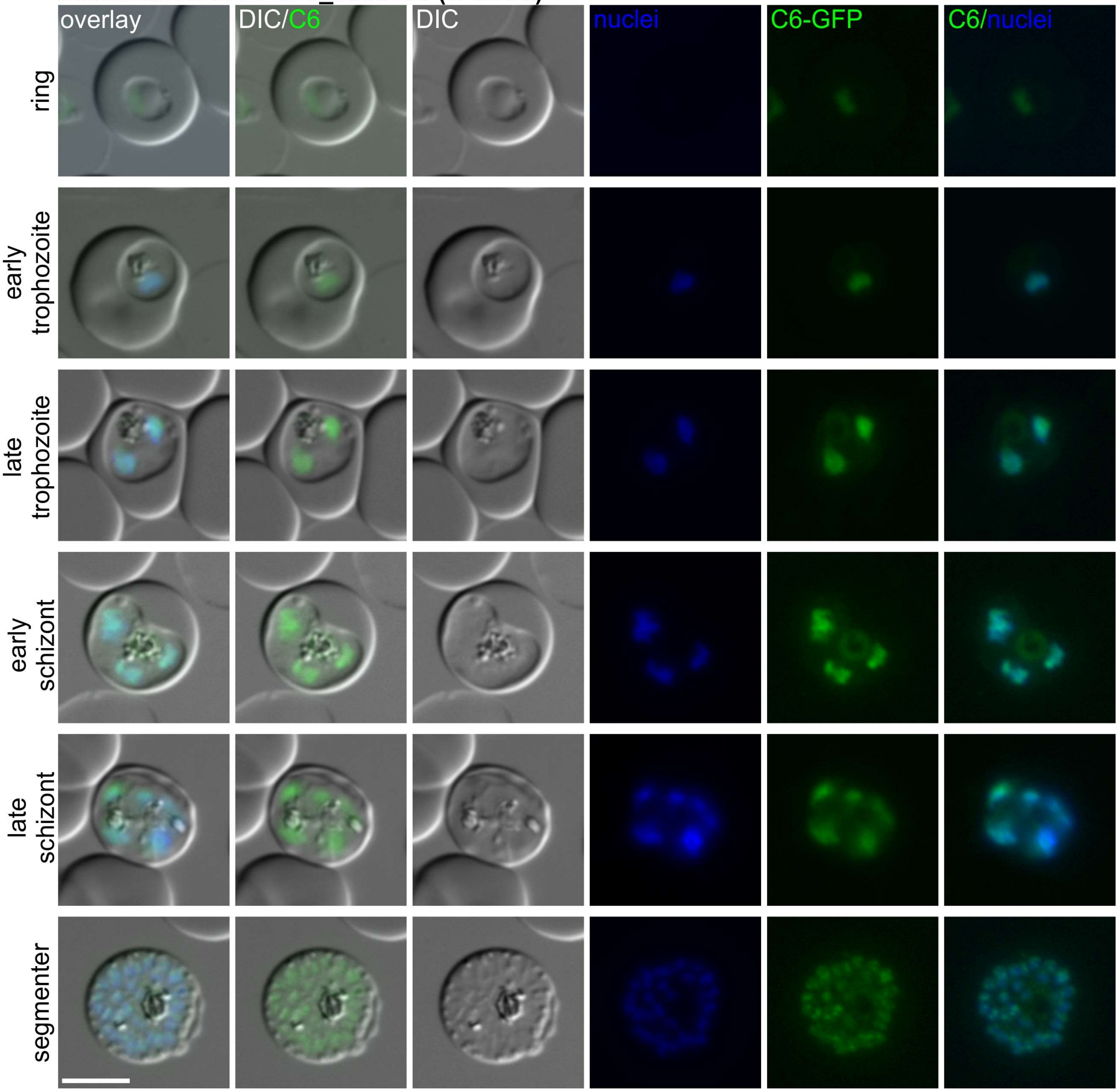

**Localisation of PF3D7_0307500 (SKA2)**

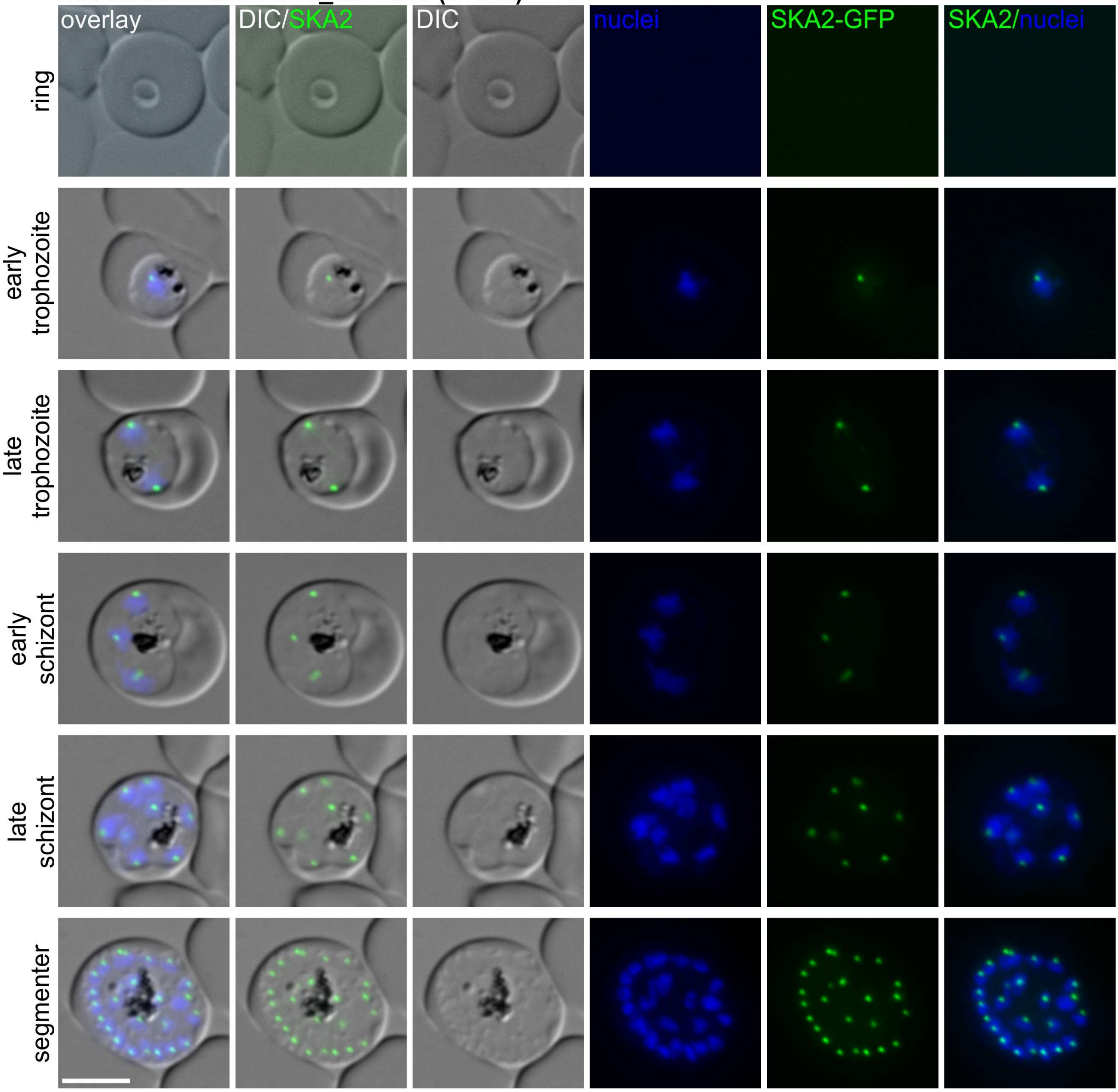

**Localisation of PF3D7_0307600 (Chr3C8)**

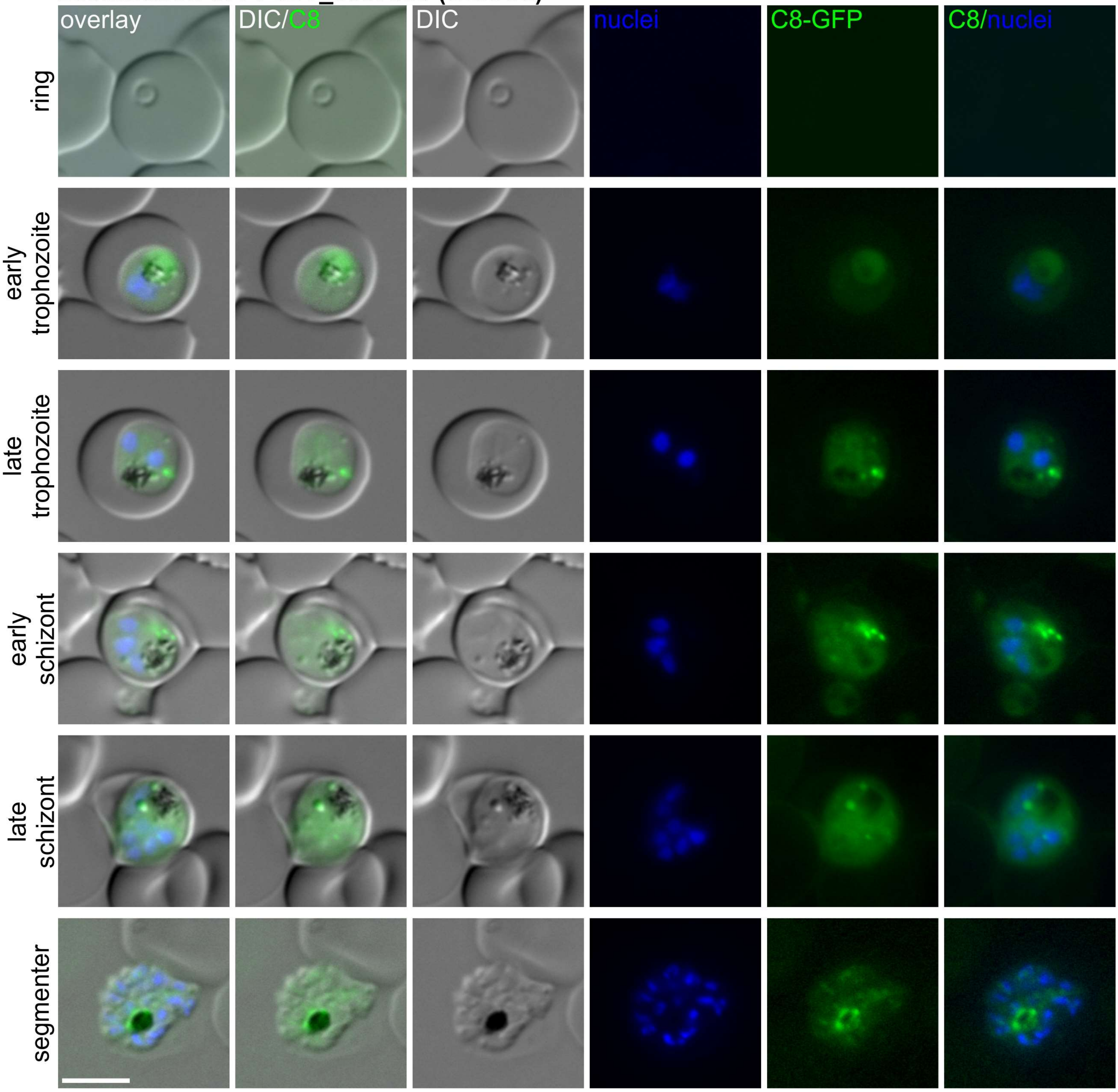

**Localisation of PF3D7_0307700 (Chr3C9)**

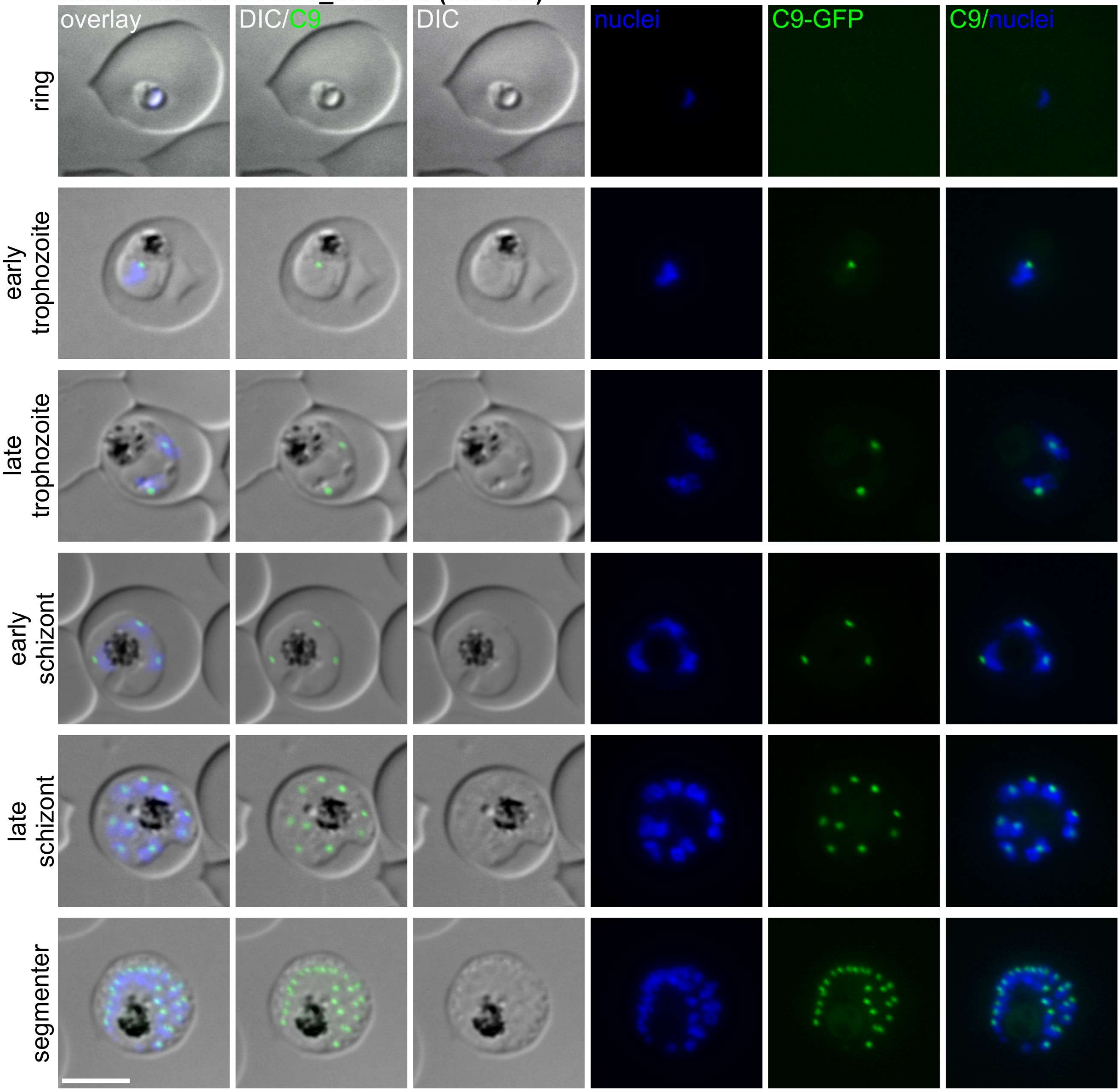

**Localisation of PF3D7_0307900 (Chr3C10)**

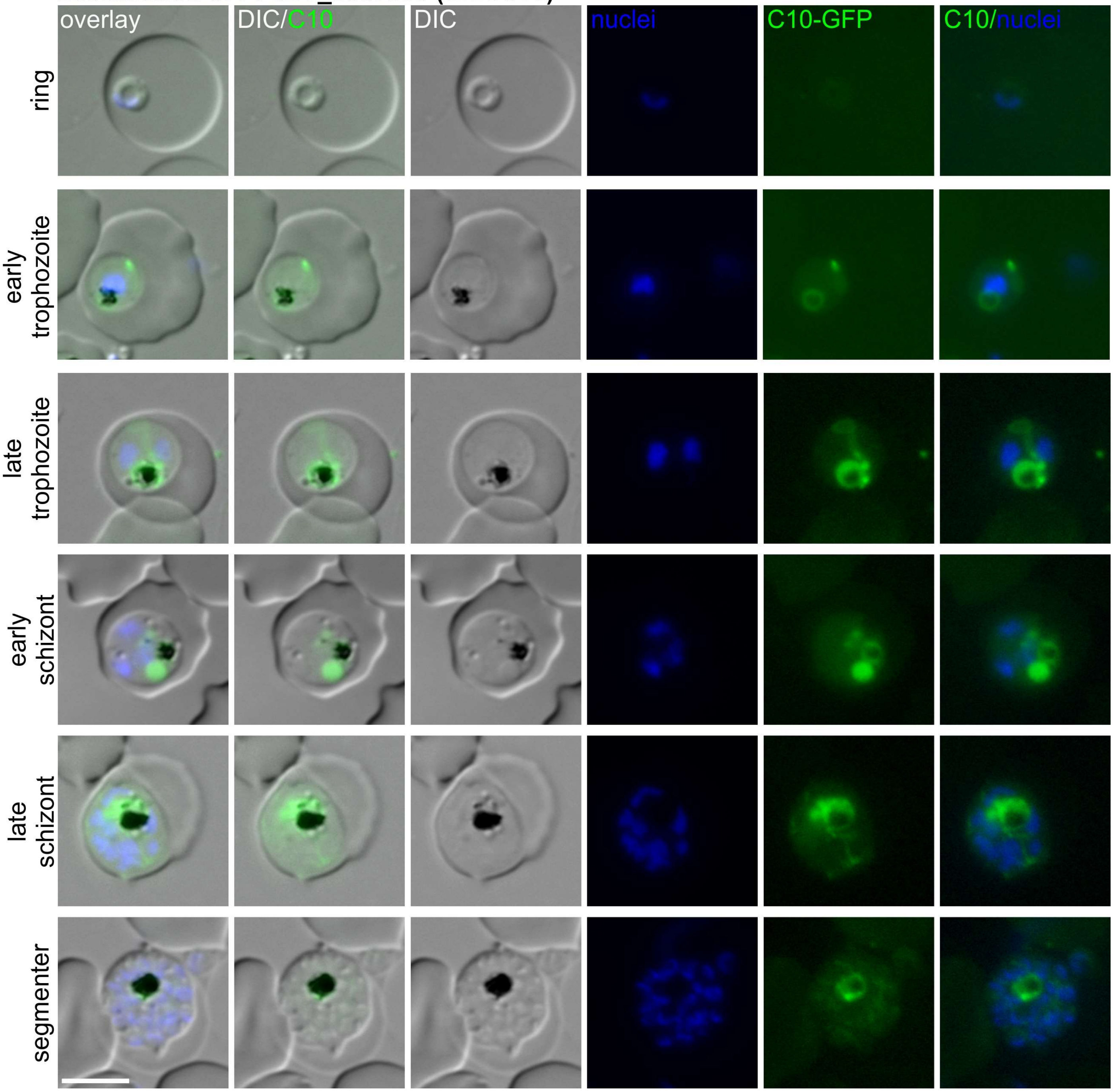

**Localisation of PF3D7_0308100 (Chr3C11)**

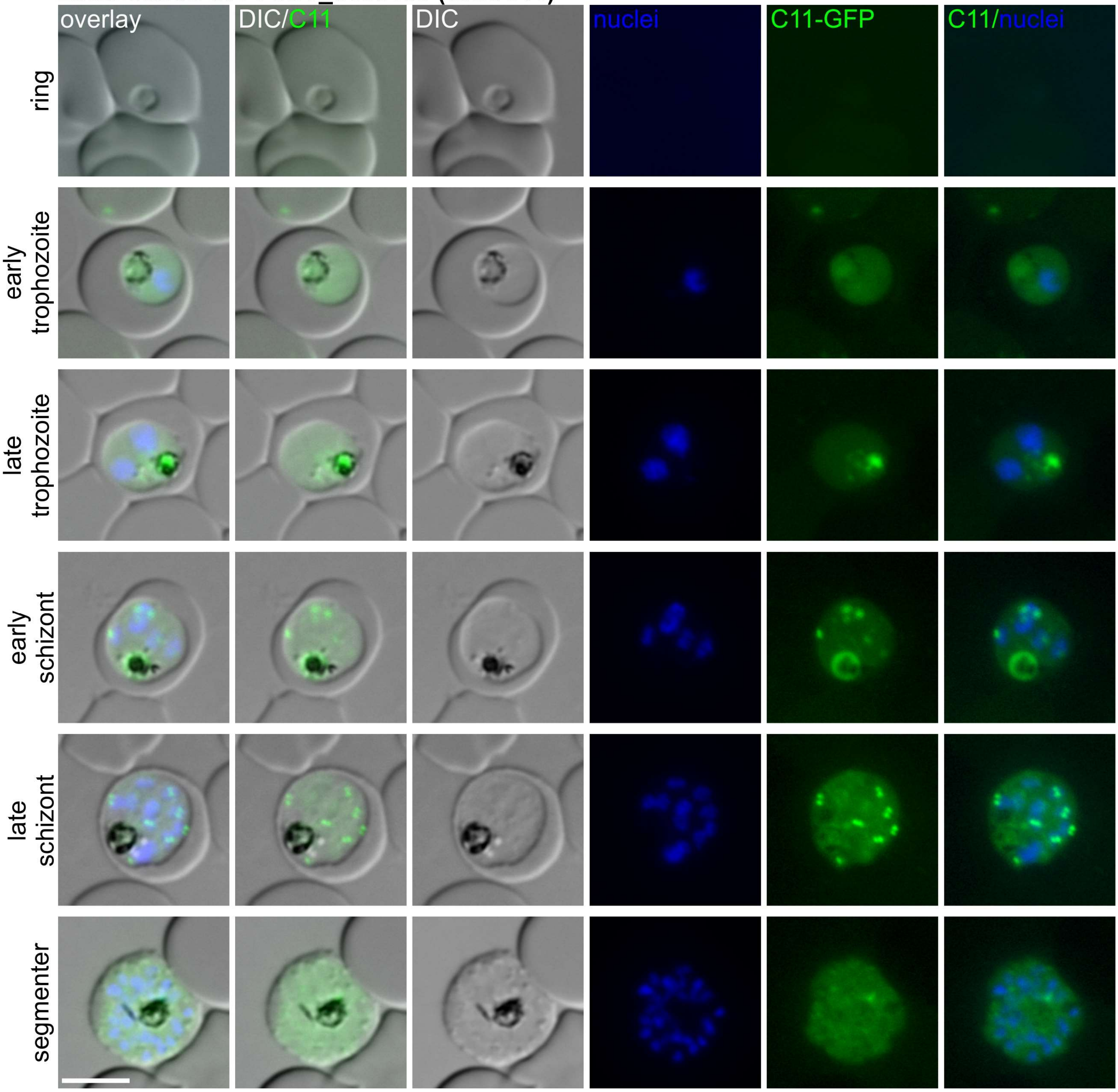

**Localisation of PF3D7_0308700 (Chr3C13)**

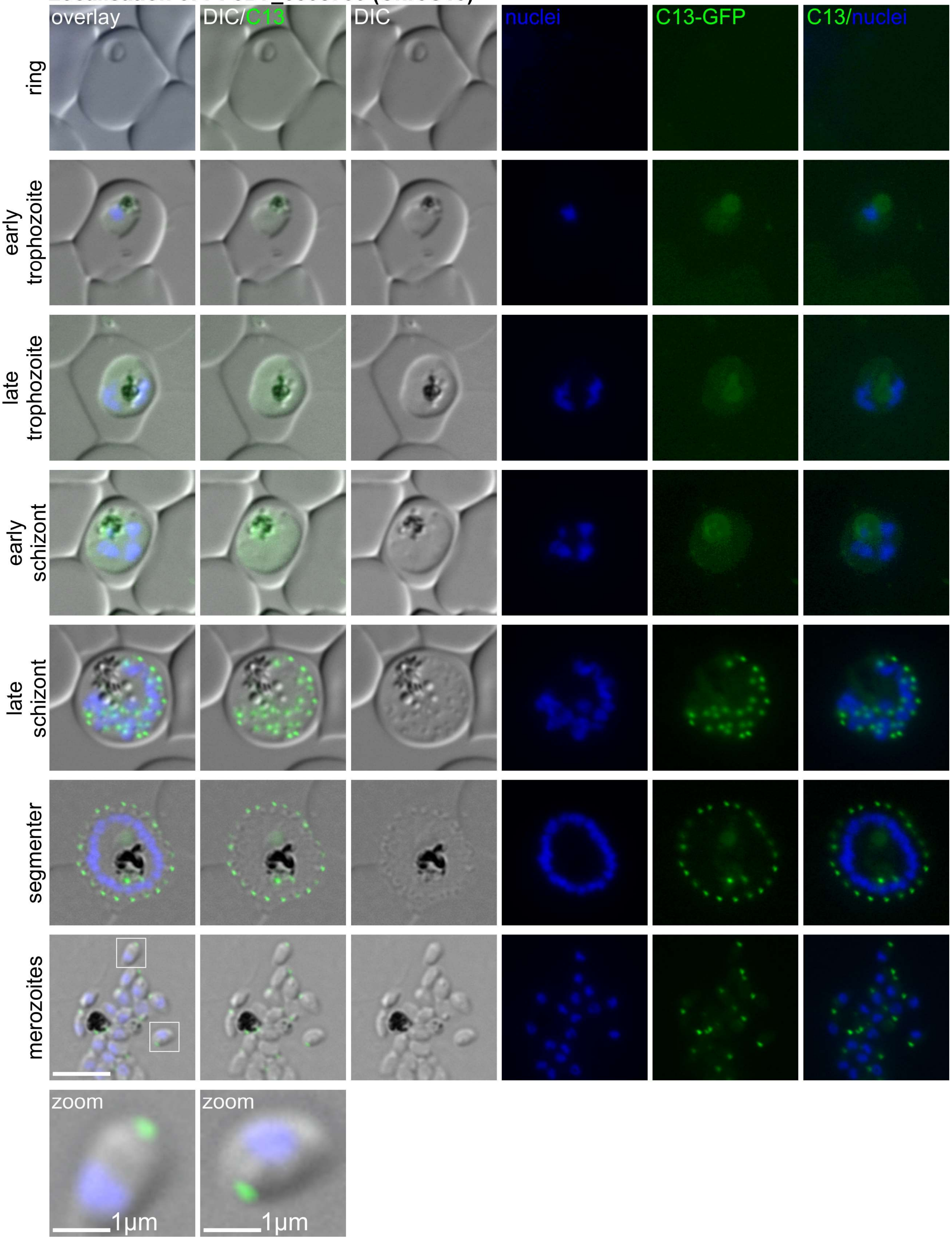

Localisation of PF3D7_030990 (Chr3C15)

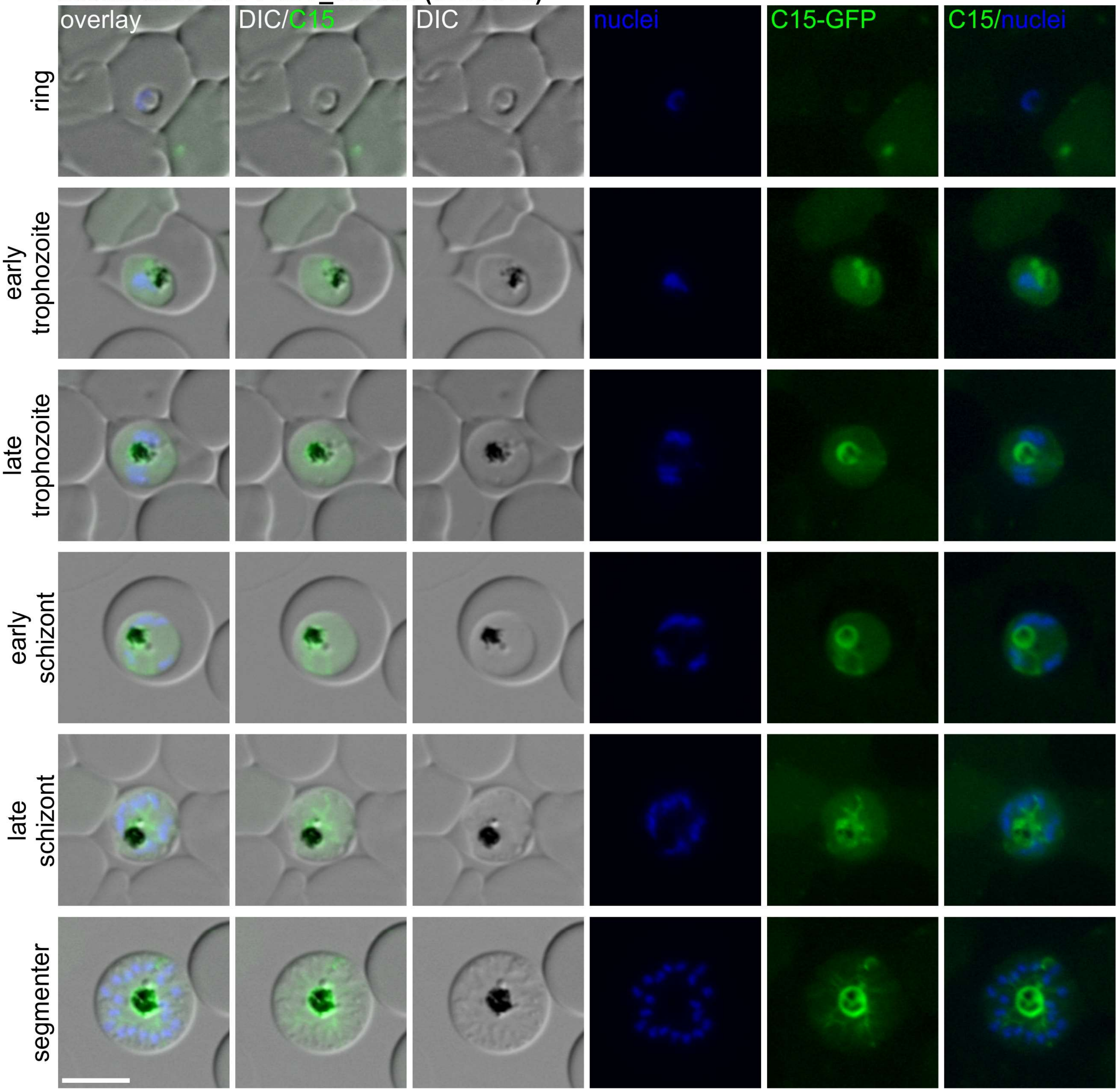

**Localisation of PF3D7_0310900 (Chr3C16)**

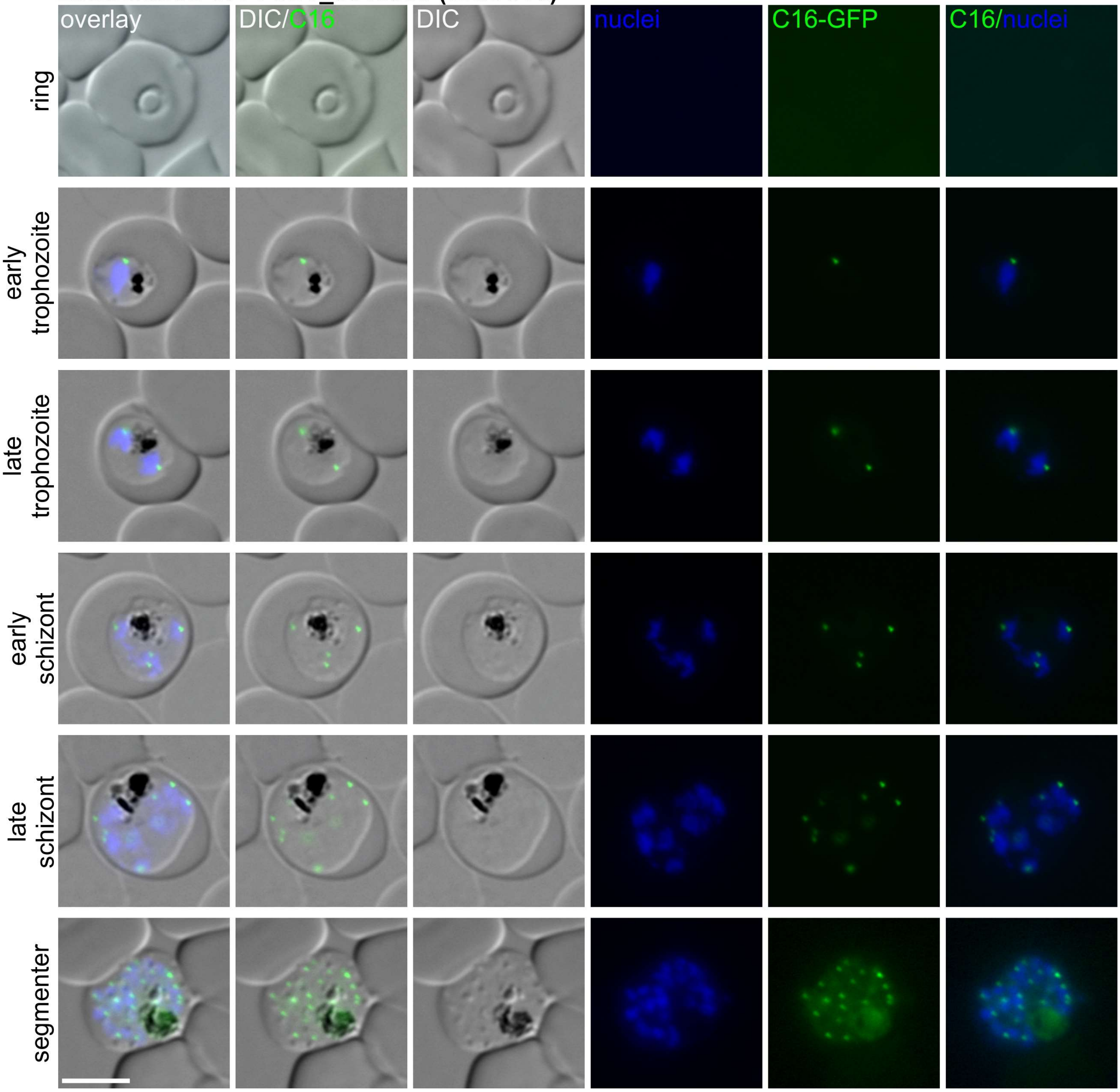

### Localisation of PF3D7_0312900 (Chr3C17)

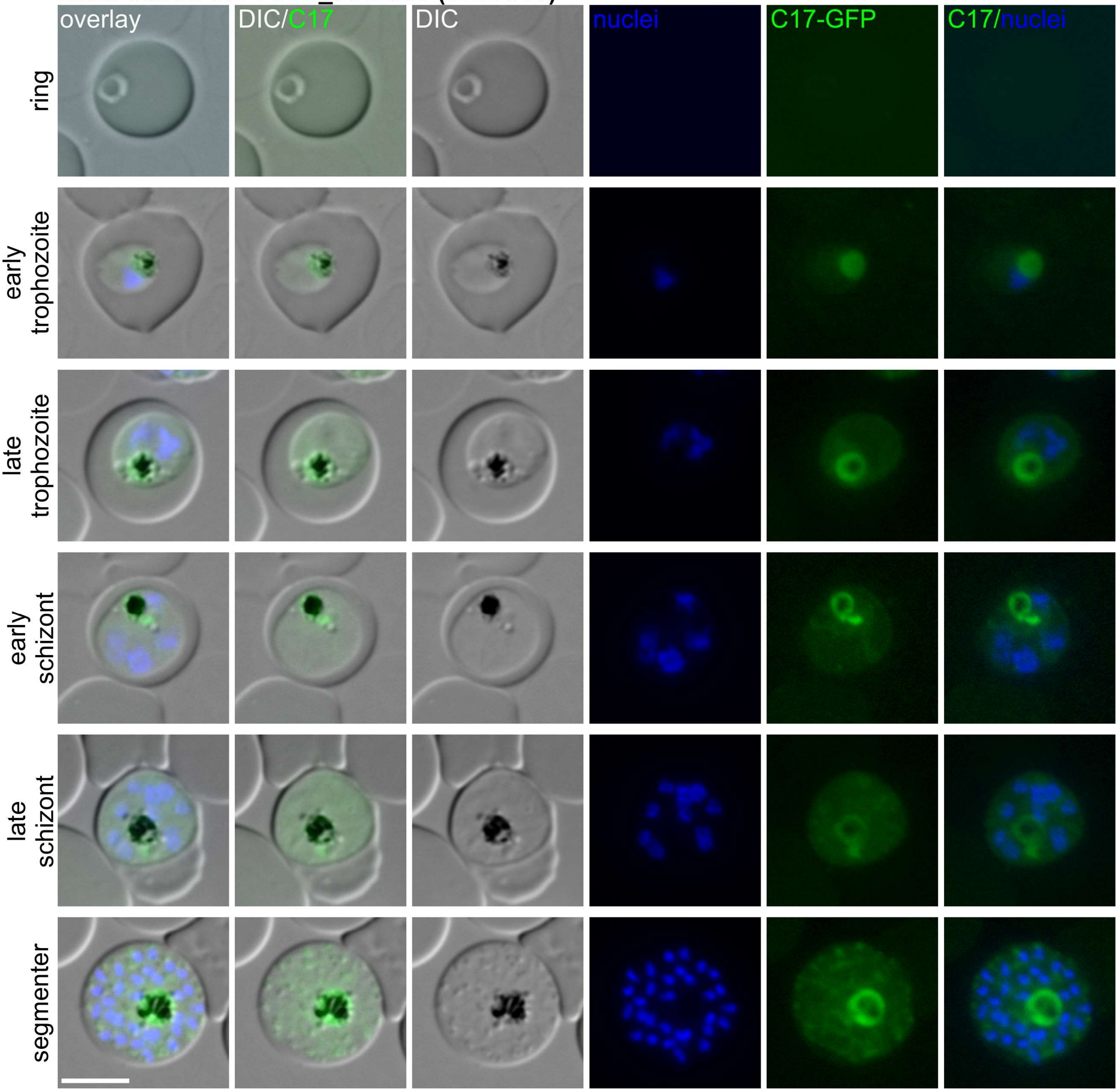

**Localisation of PF3D7_0313000 (Chr3C18)**

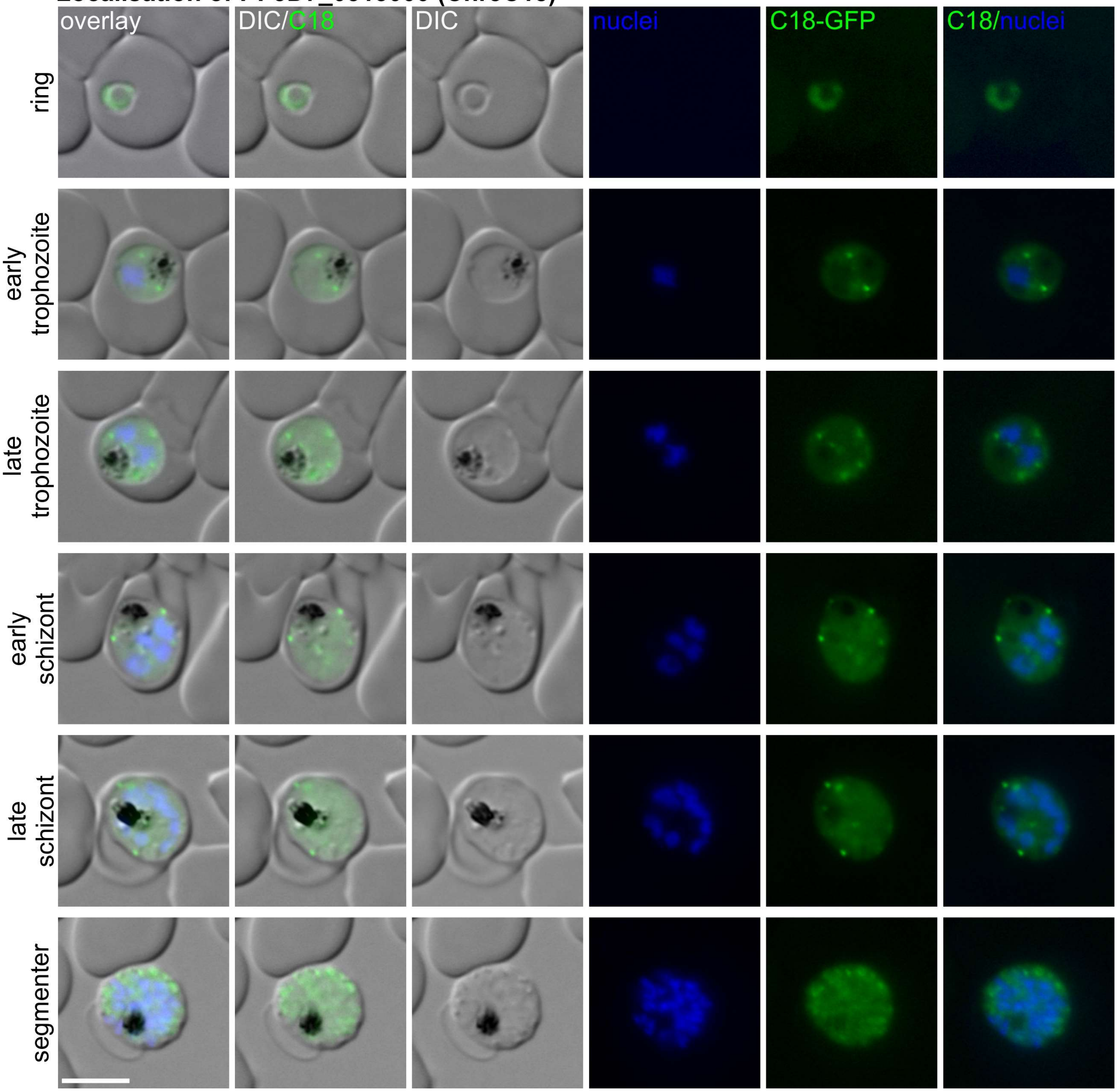

**Localisation of PF3D7_0313200 (Chr3C19)**

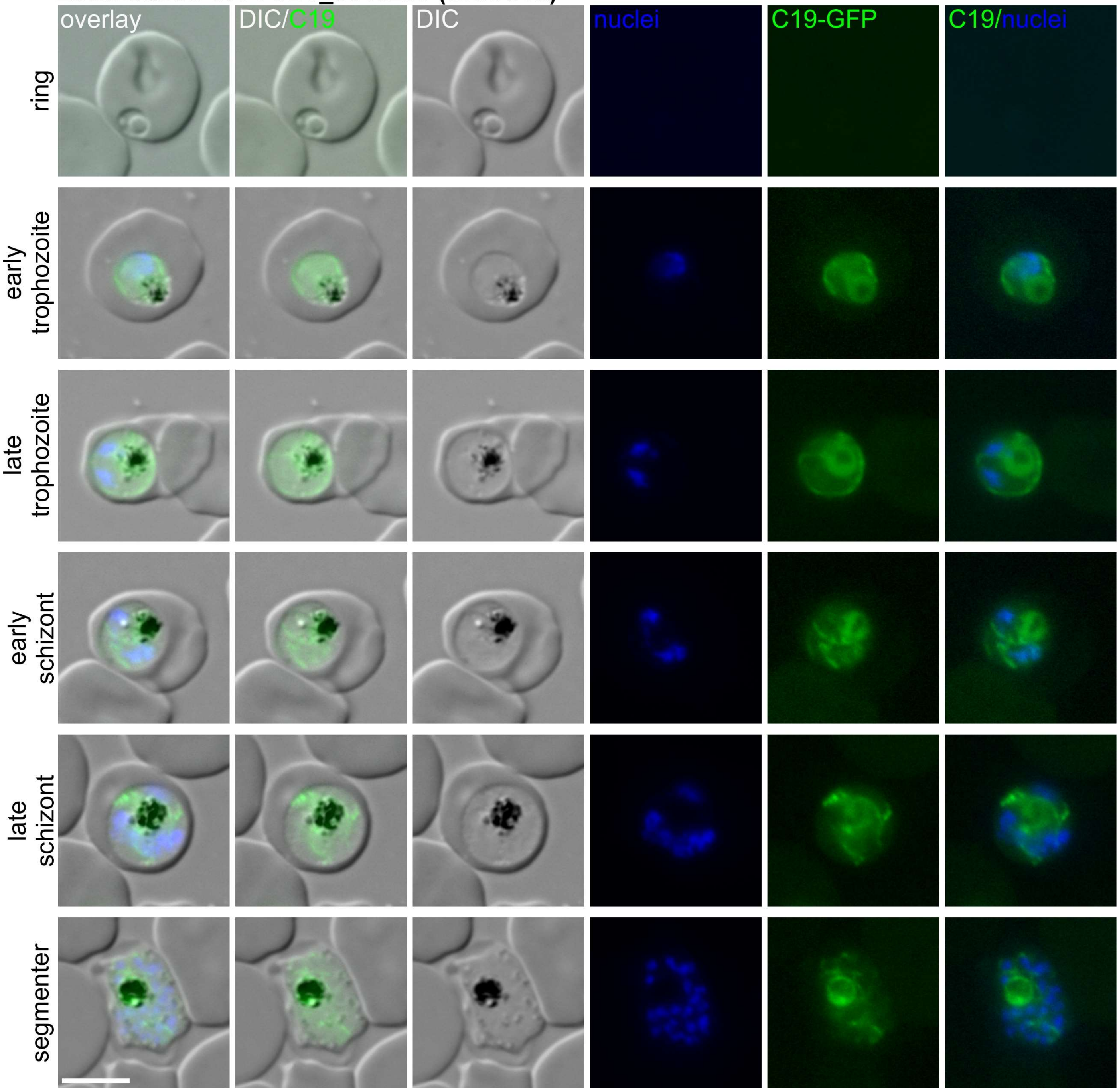

**Localisation of PF3D7_0313400 (Chr3C20)**

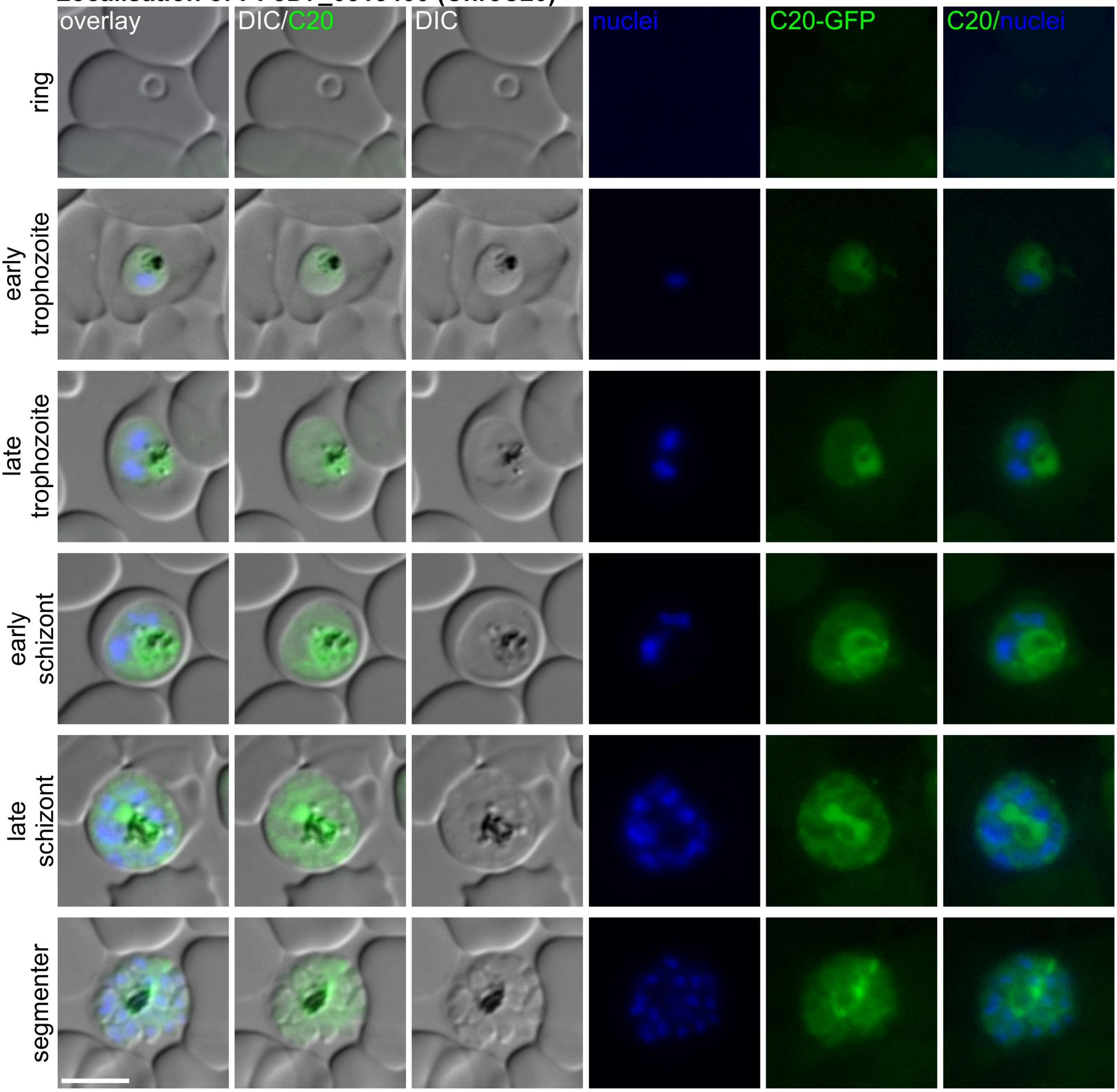

**Localisation of PF3D7_0313600 (Chr3C21)**

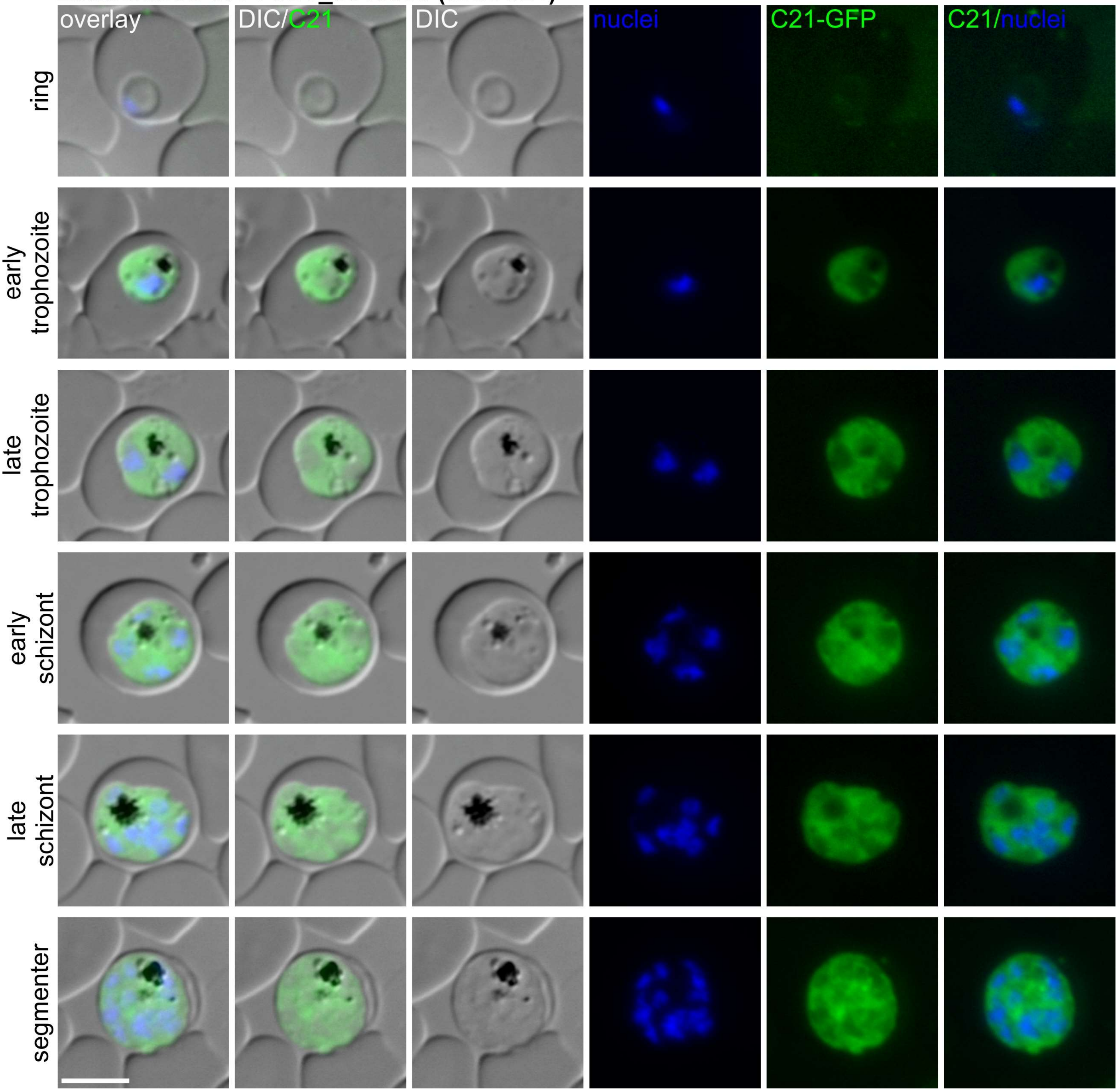

**Localisation of PF3D7_0314500 (Chr3C22)**

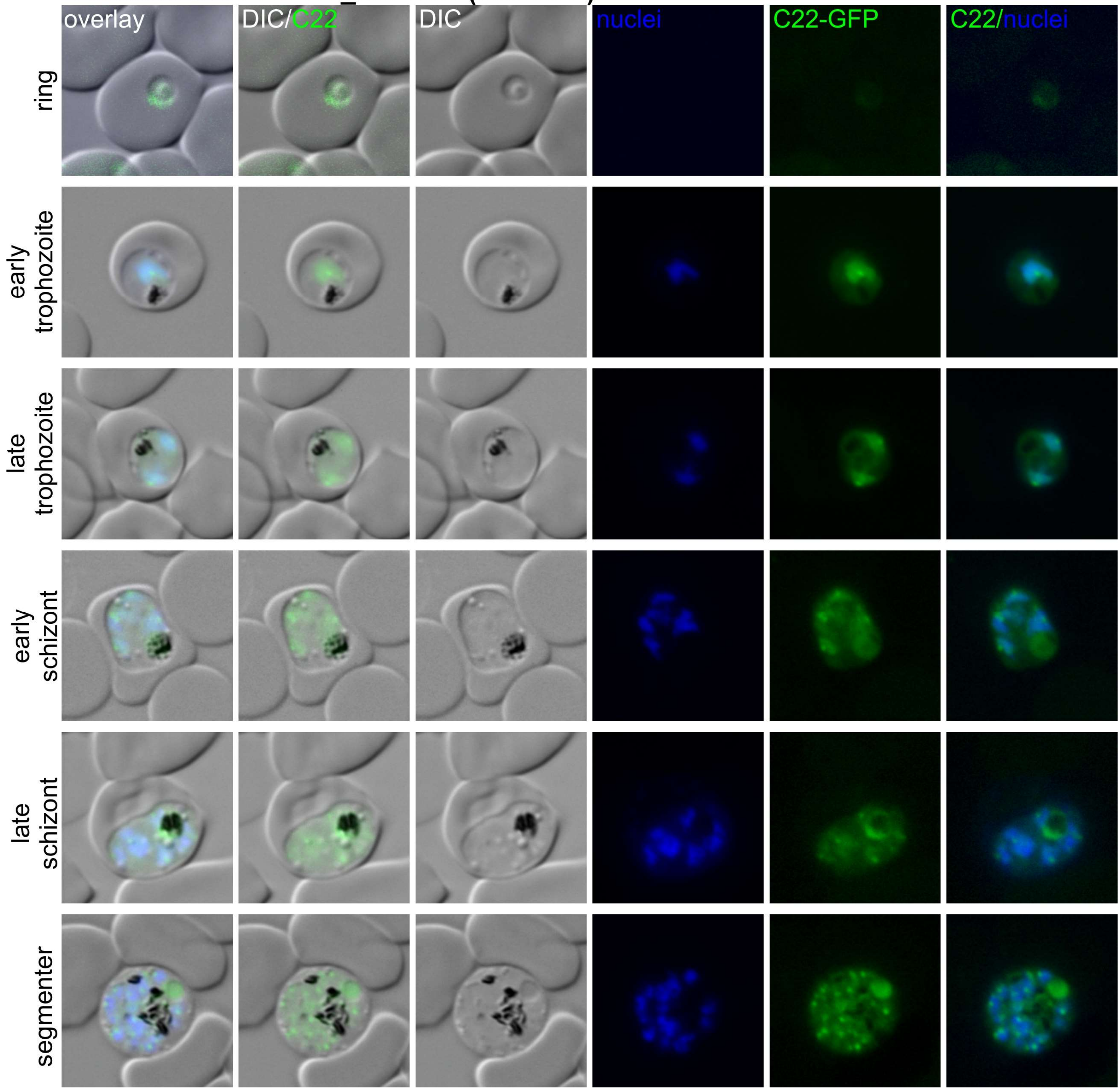

**Localisation of PF3D7_0314600 (Chr3C23)**

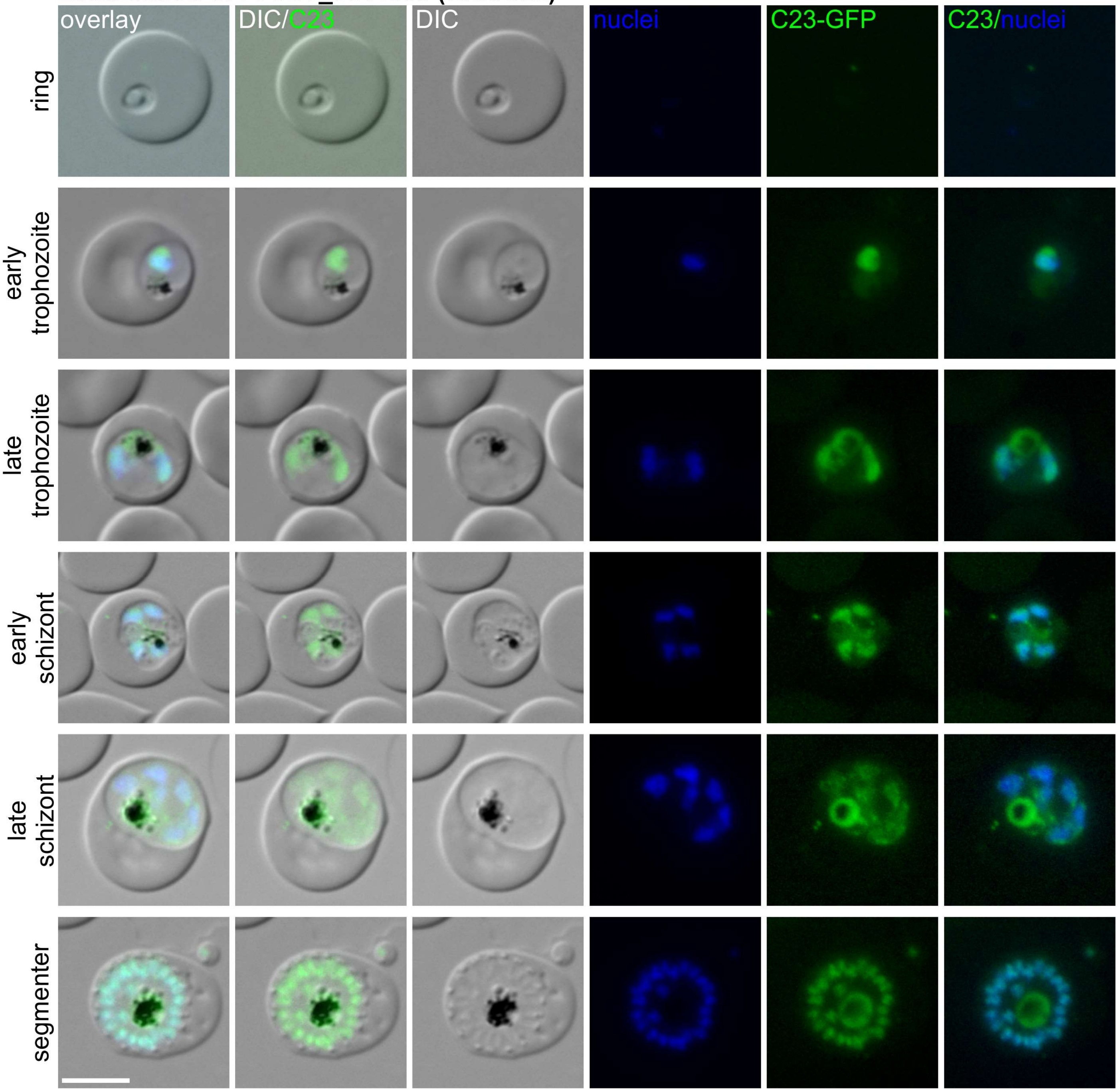

**Localisation of PF3D7_0314700 (Chr3C24)**

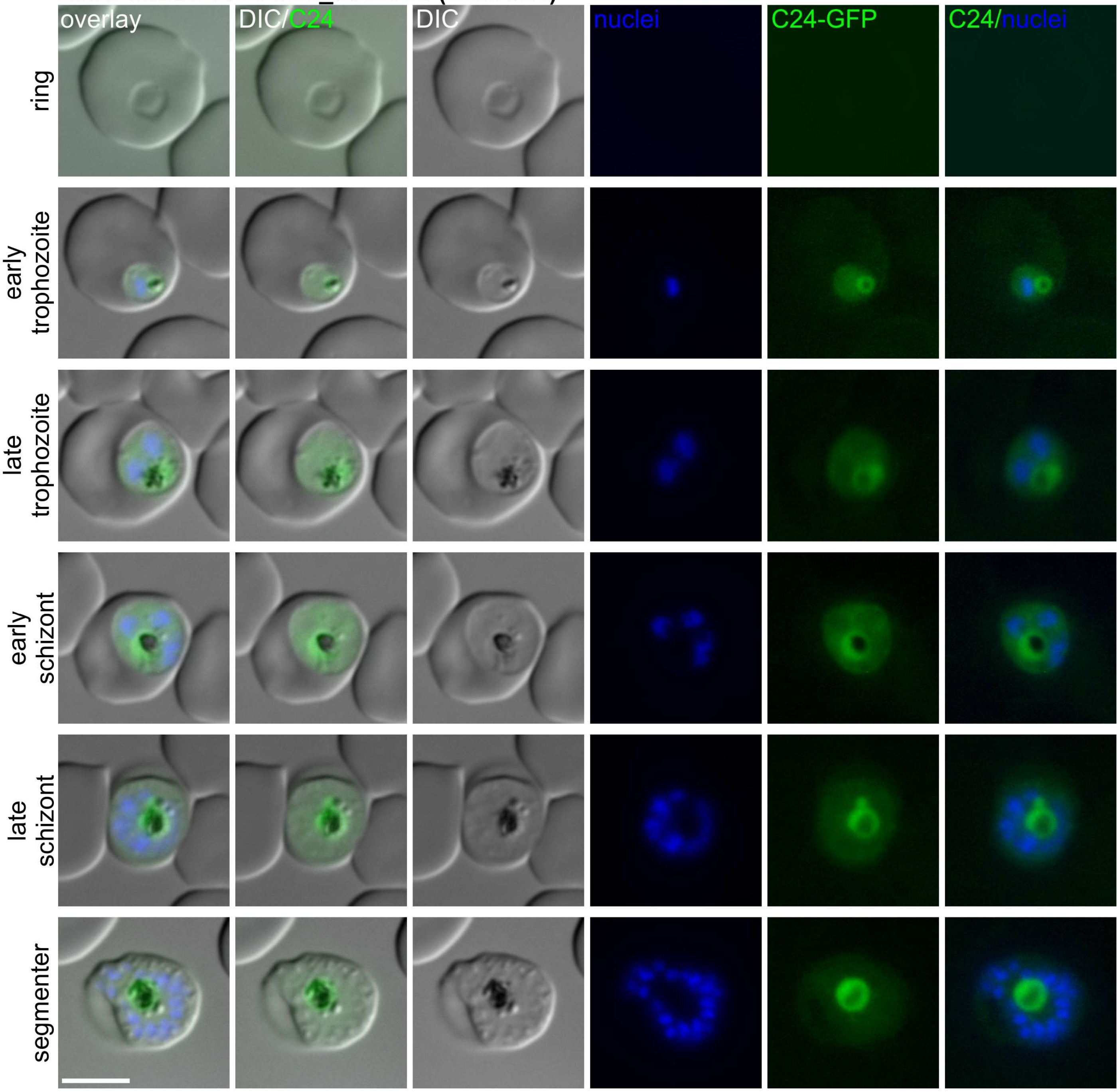

**Localisation of PF3D7_0314800 (Chr3C25)**

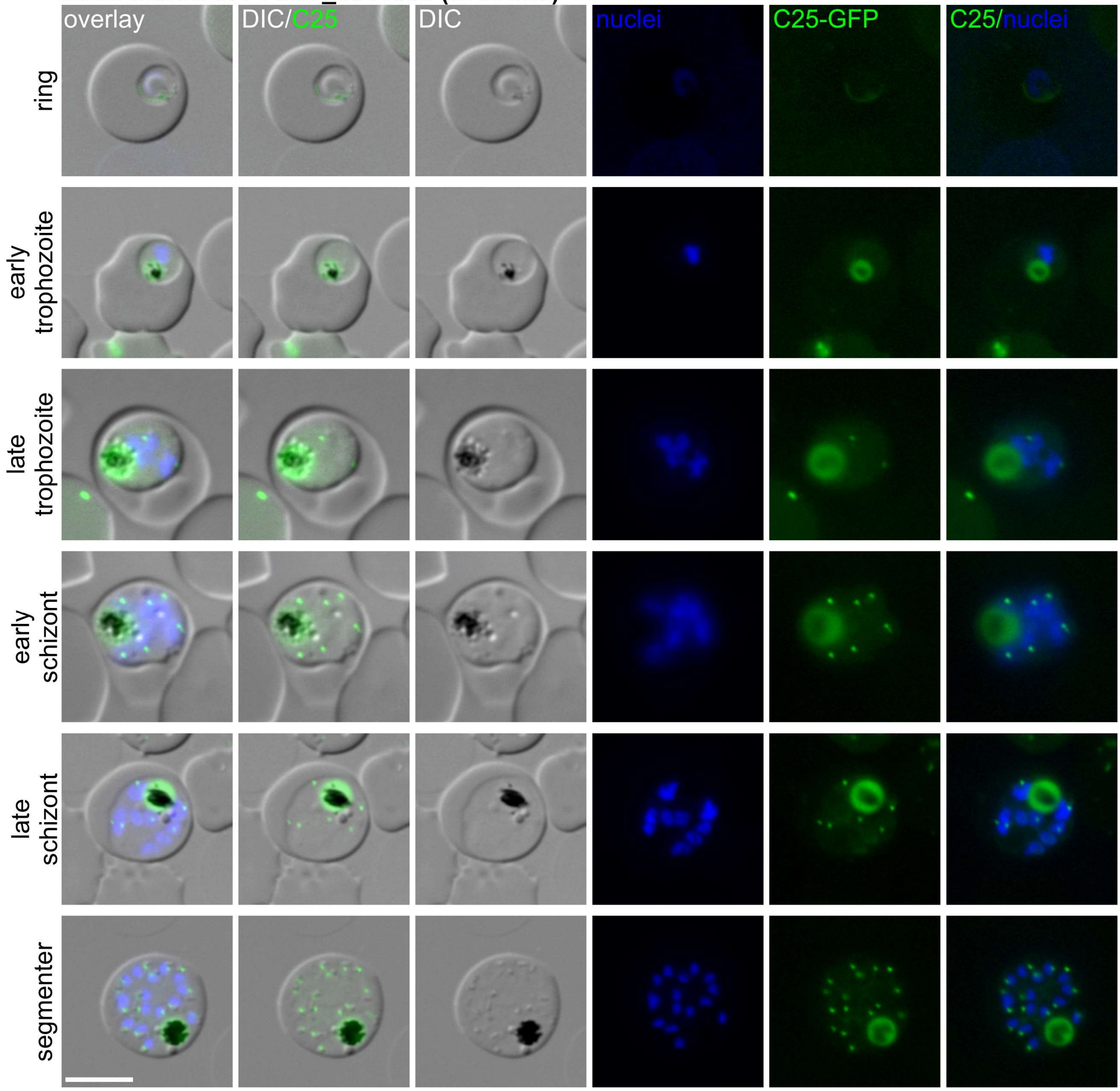

**Localisation of PF3D7_0315000 (Chr3C26)**

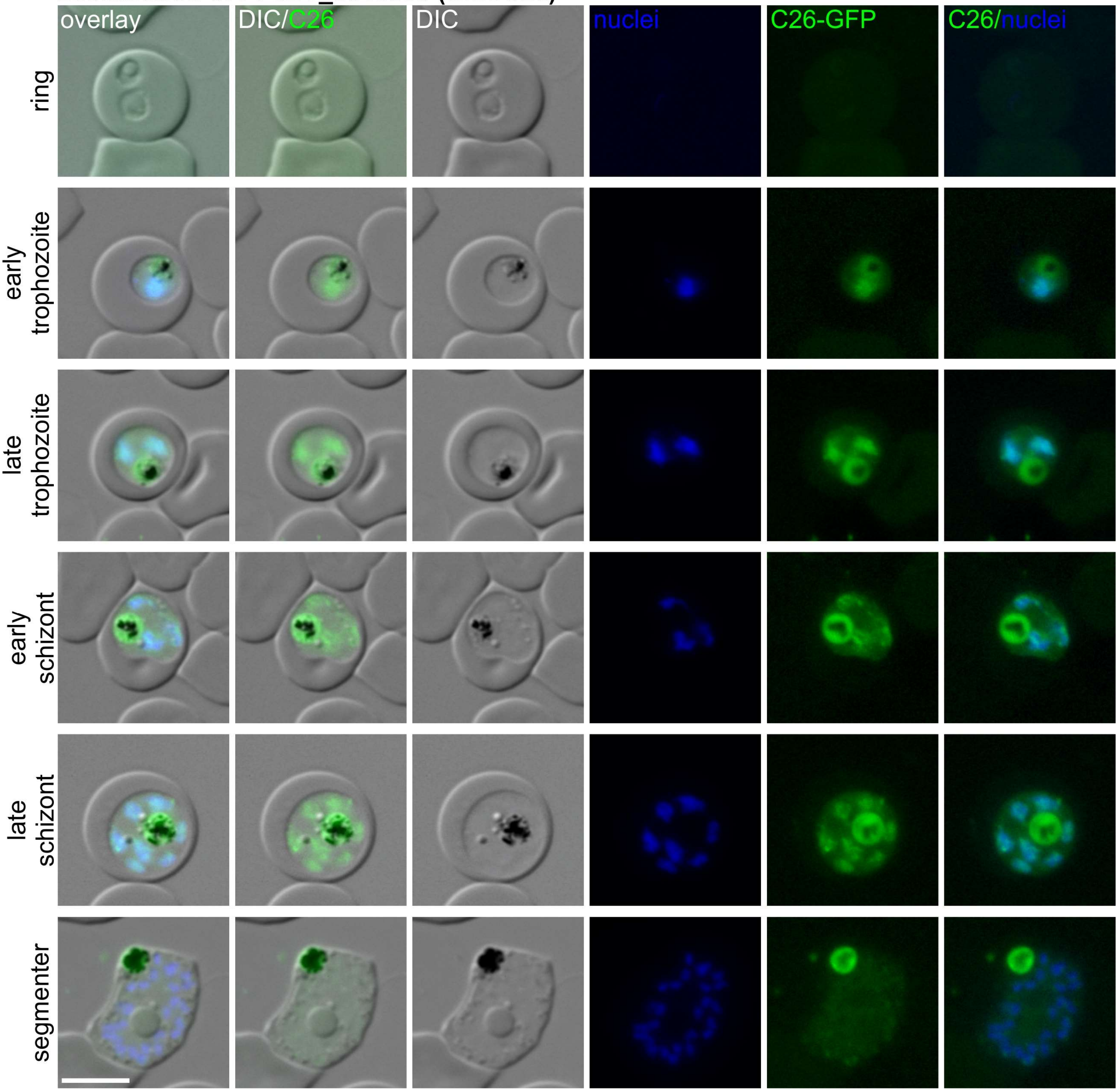

**Localisation of PF3D7_0315600 (Chr3C27)**

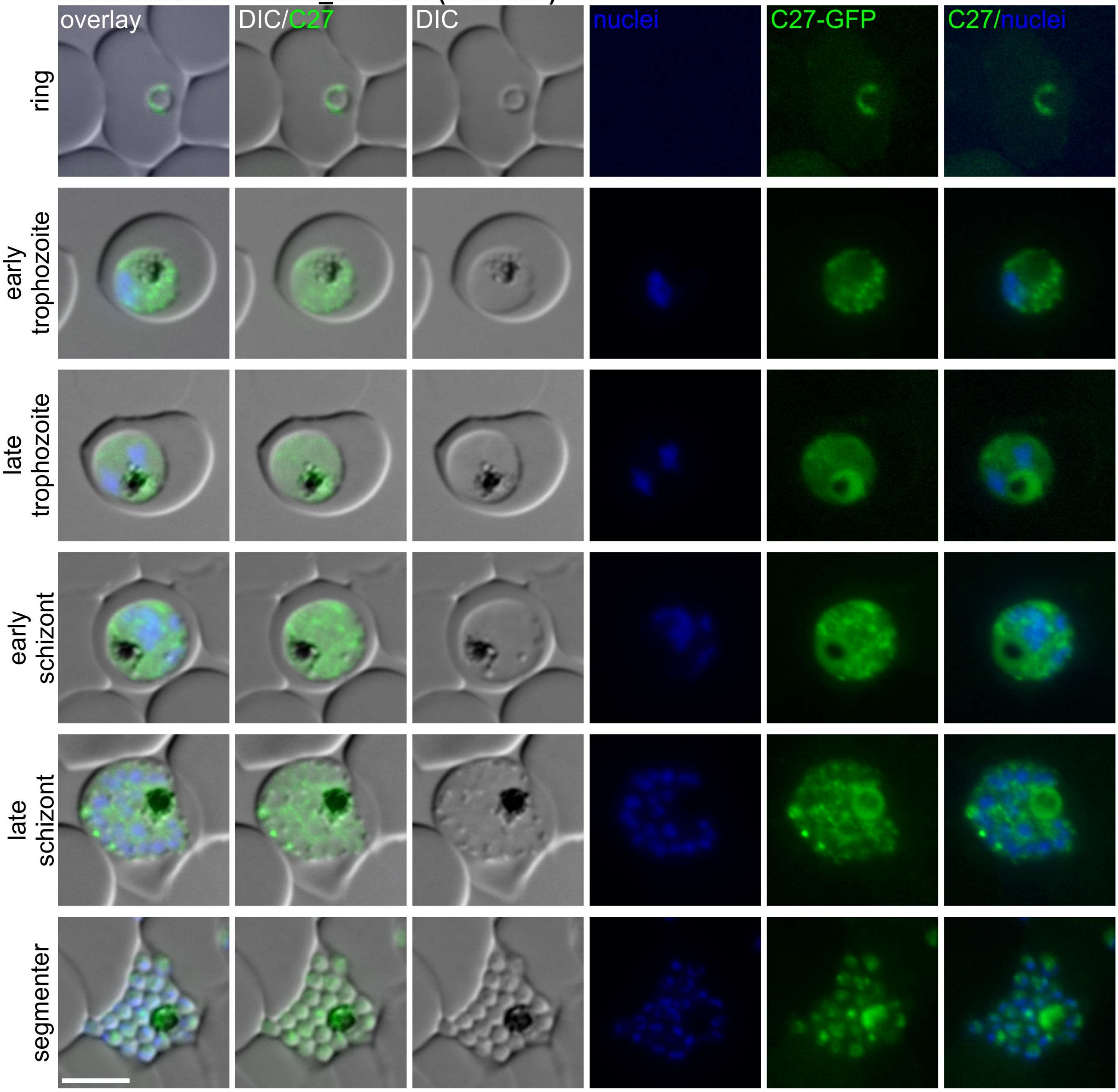

**Localisation of PF3D7_0315800 (Chr3C28)**

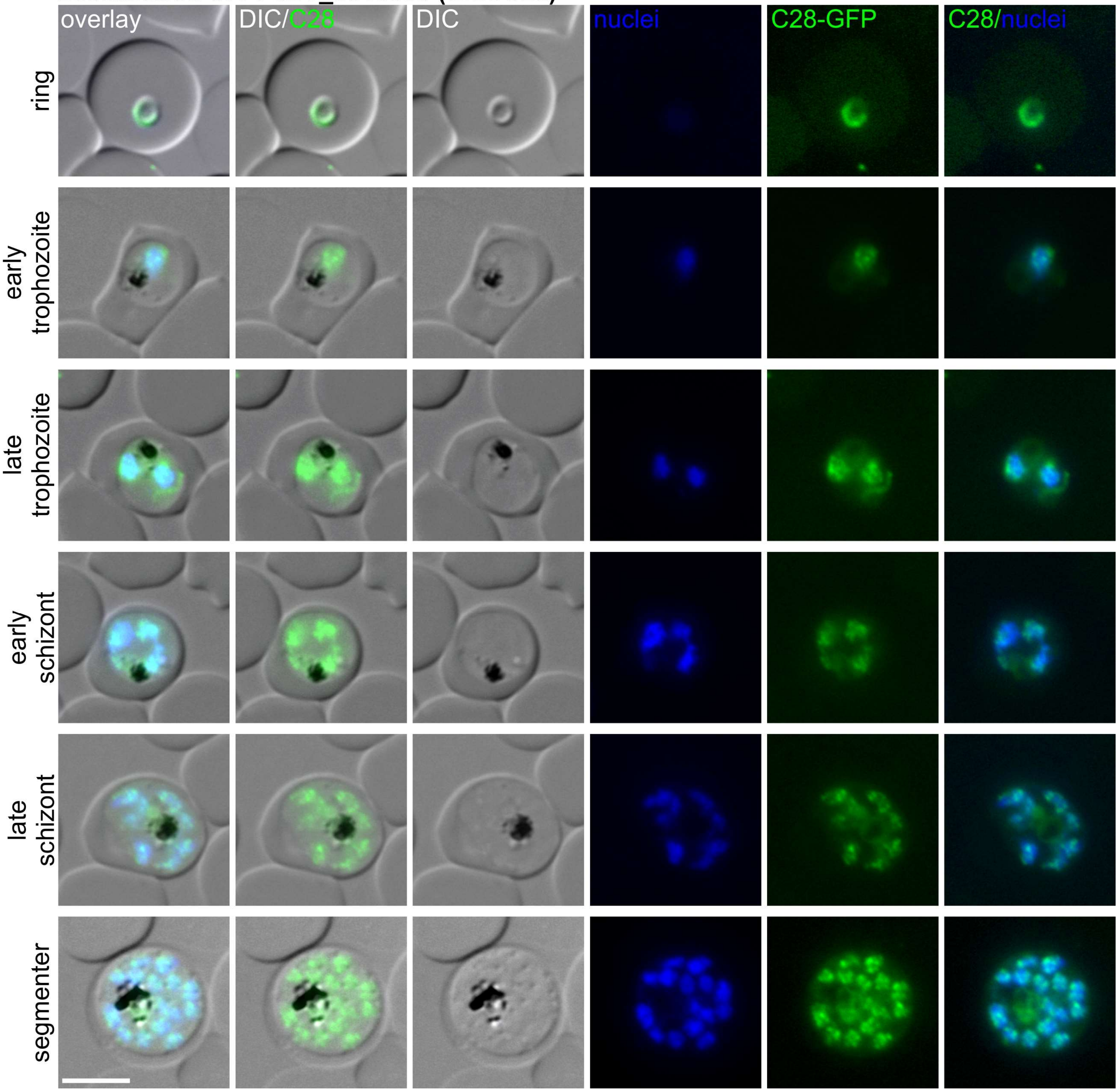

**Localisation of PF3D7_0317300 (Chr3C29)**

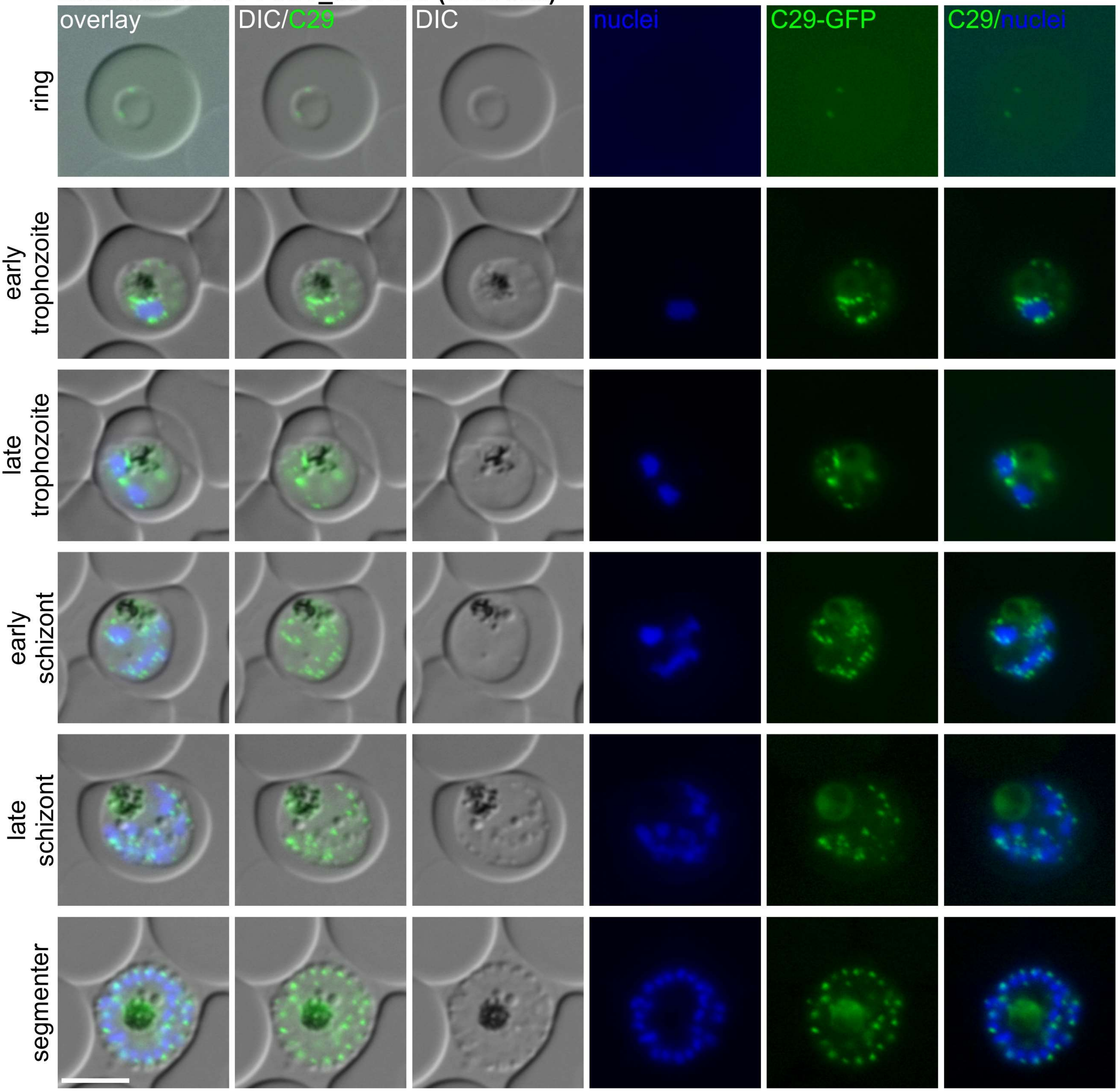

**Localisation of PF3D7_0317400 (Chr3C30)**

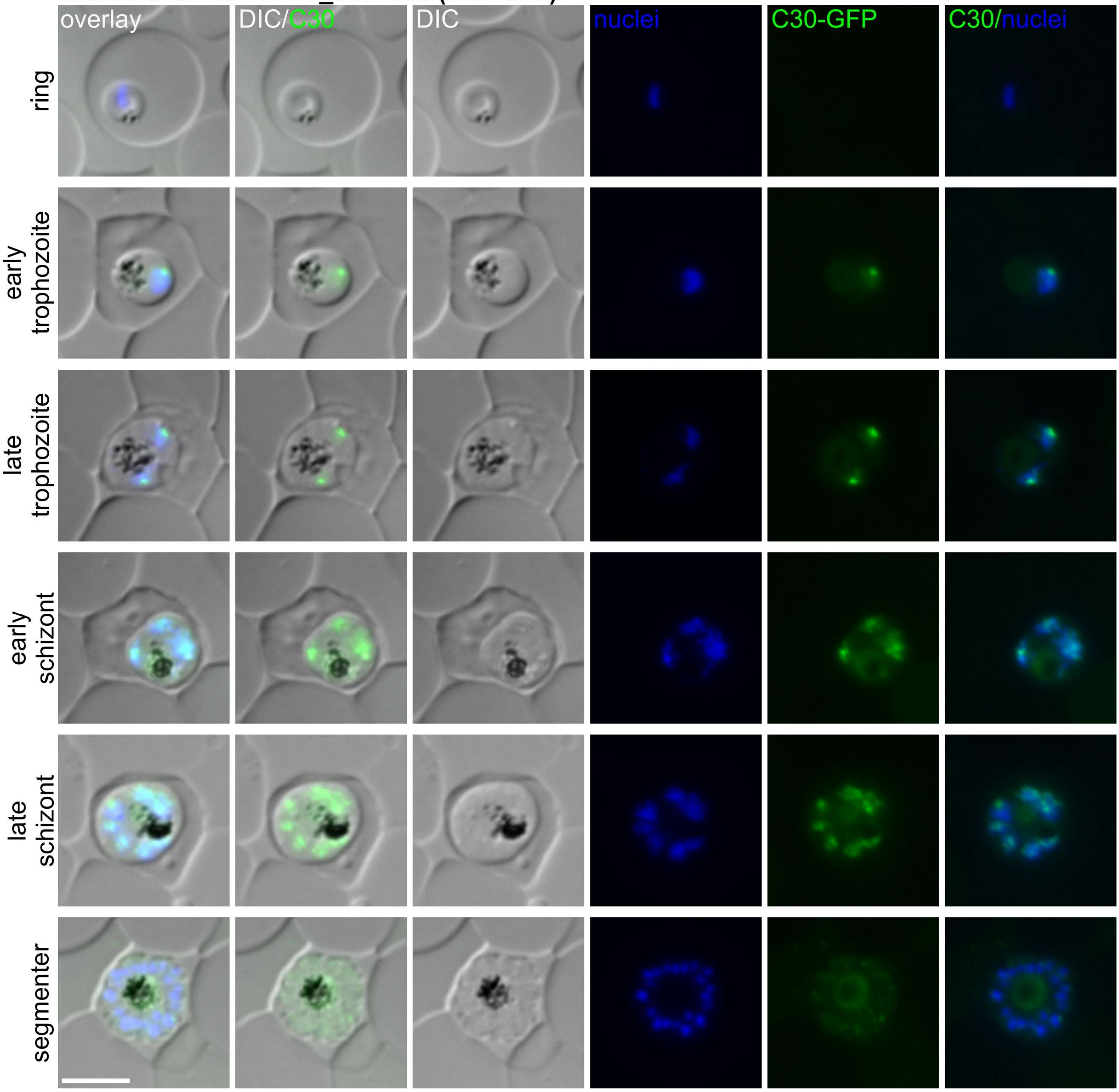

**Localisation of PF3D7_0319900 (Chr3C31)**

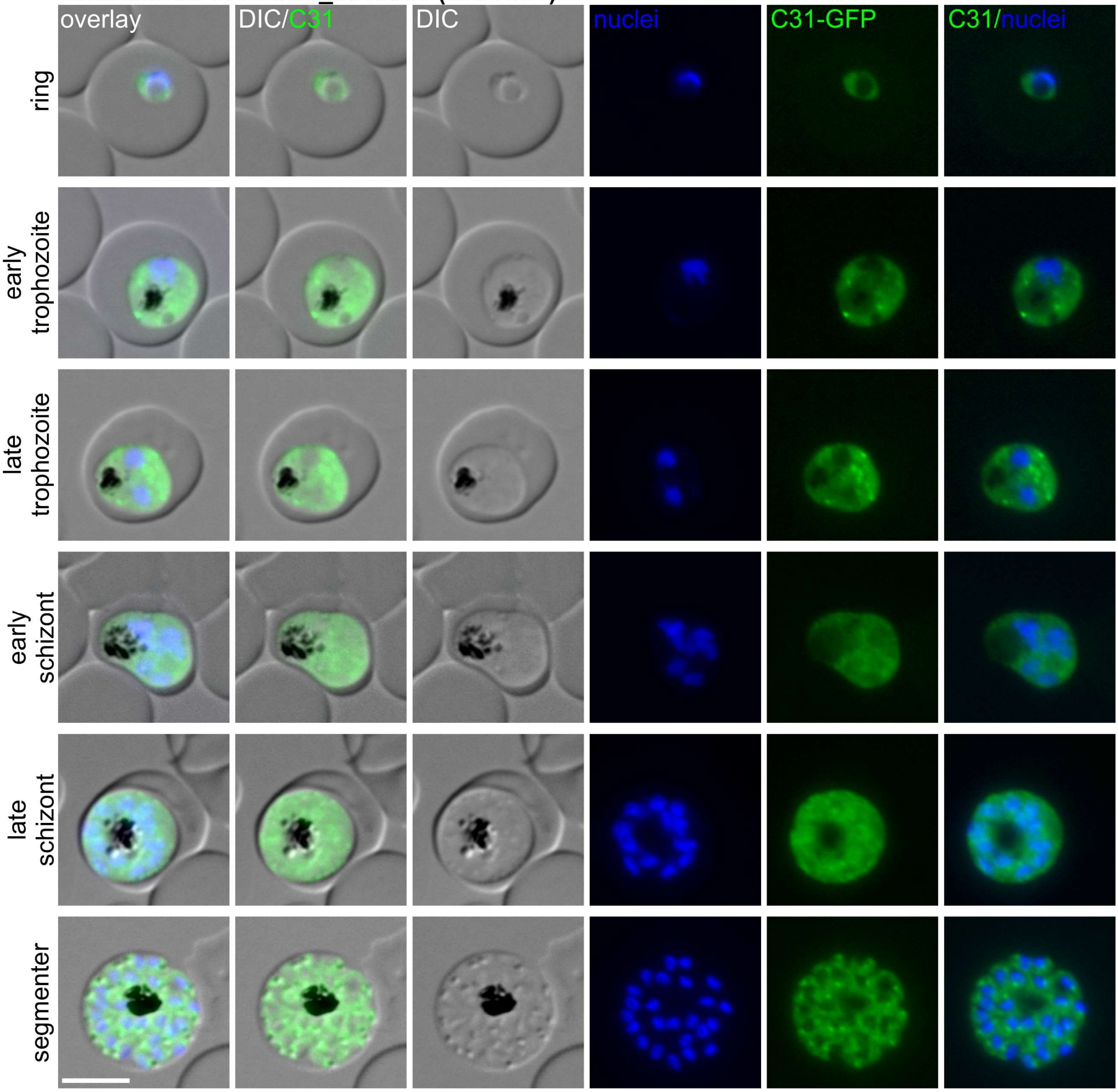

**Localisation of PF3D7_0320600 (Chr3C32)**

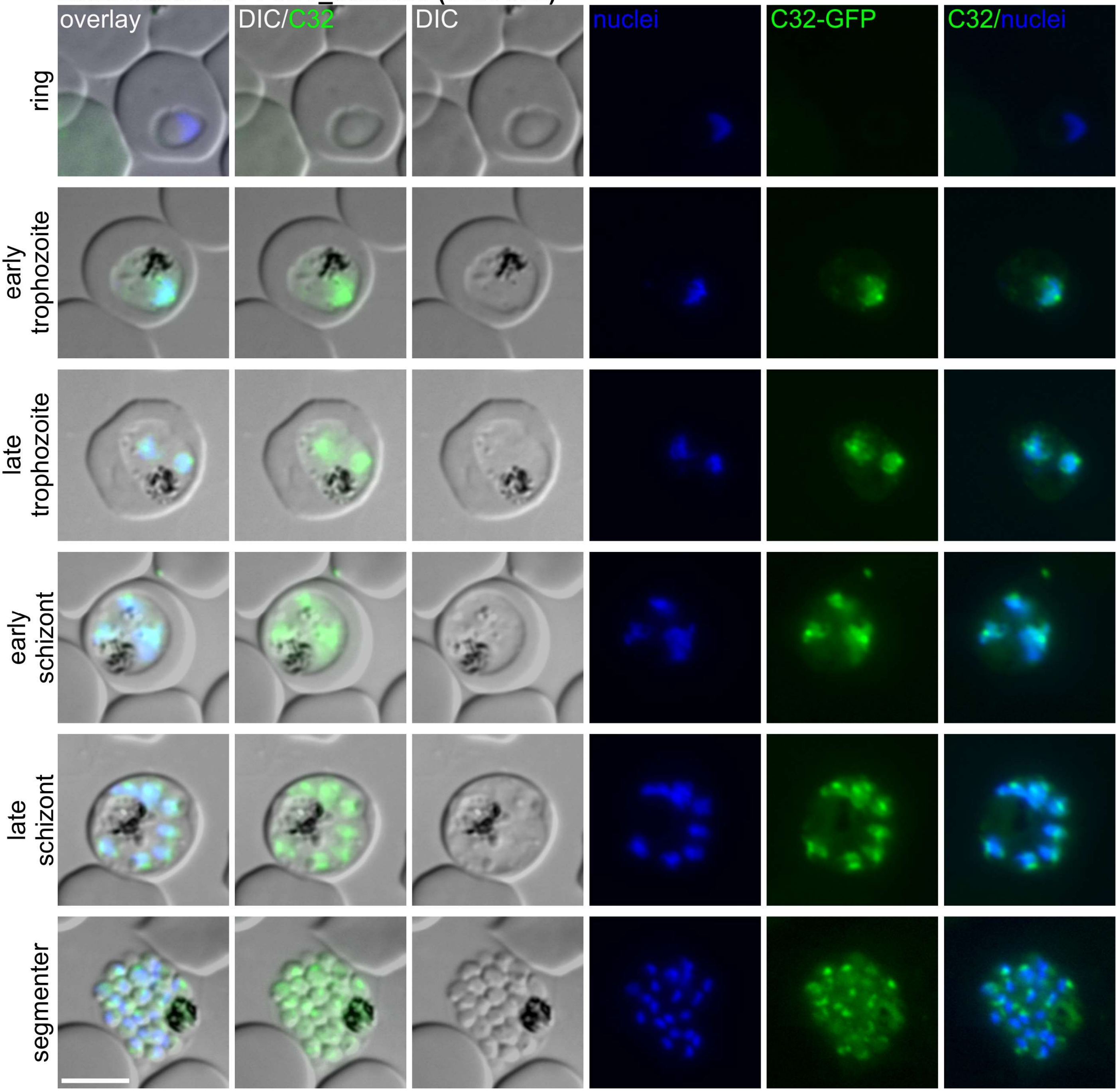

**Localisation of PF3D7_0323800 (Chr3C33)**

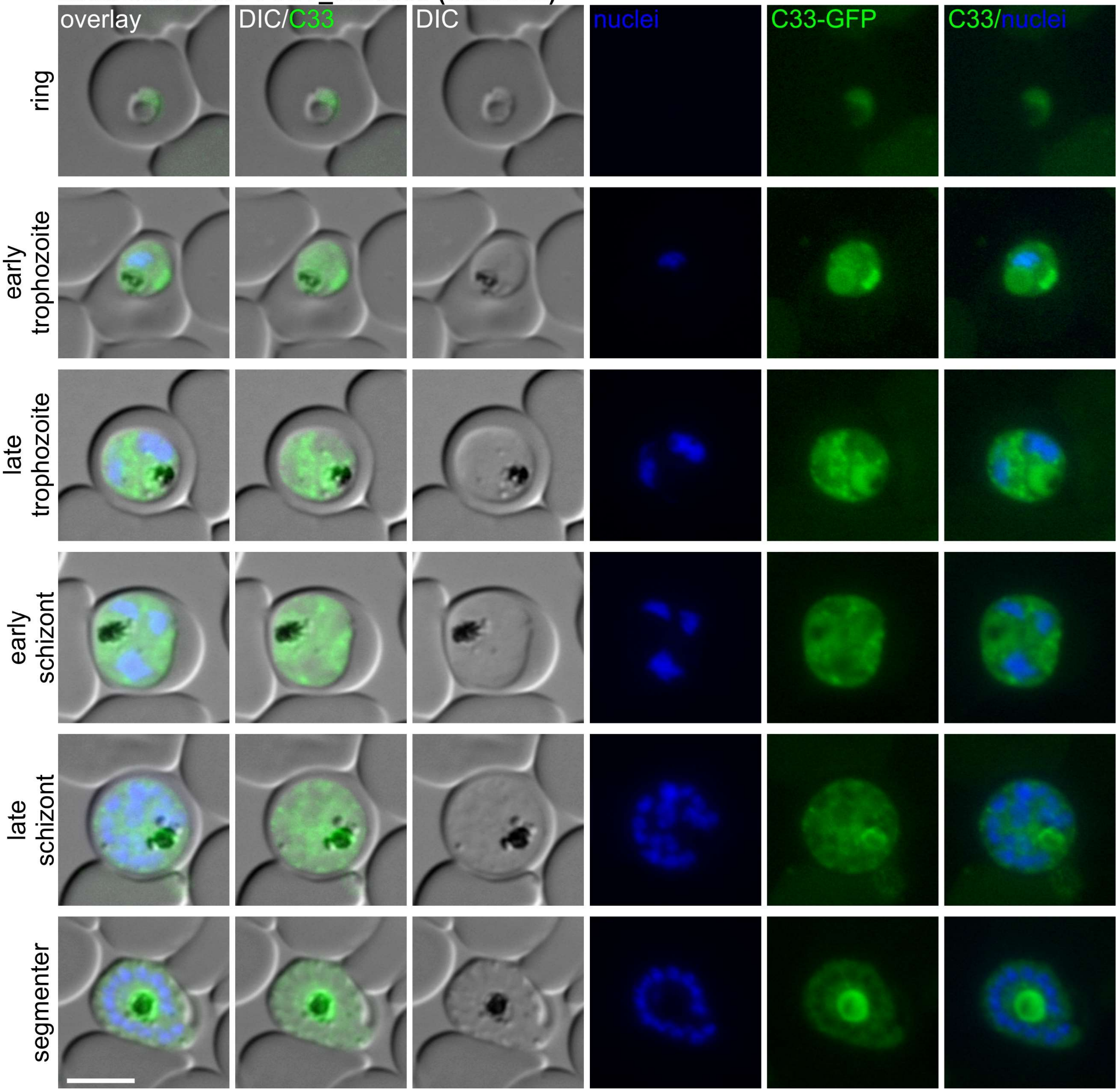

**Localisation of PF3D7_0911200 (SIC1)**

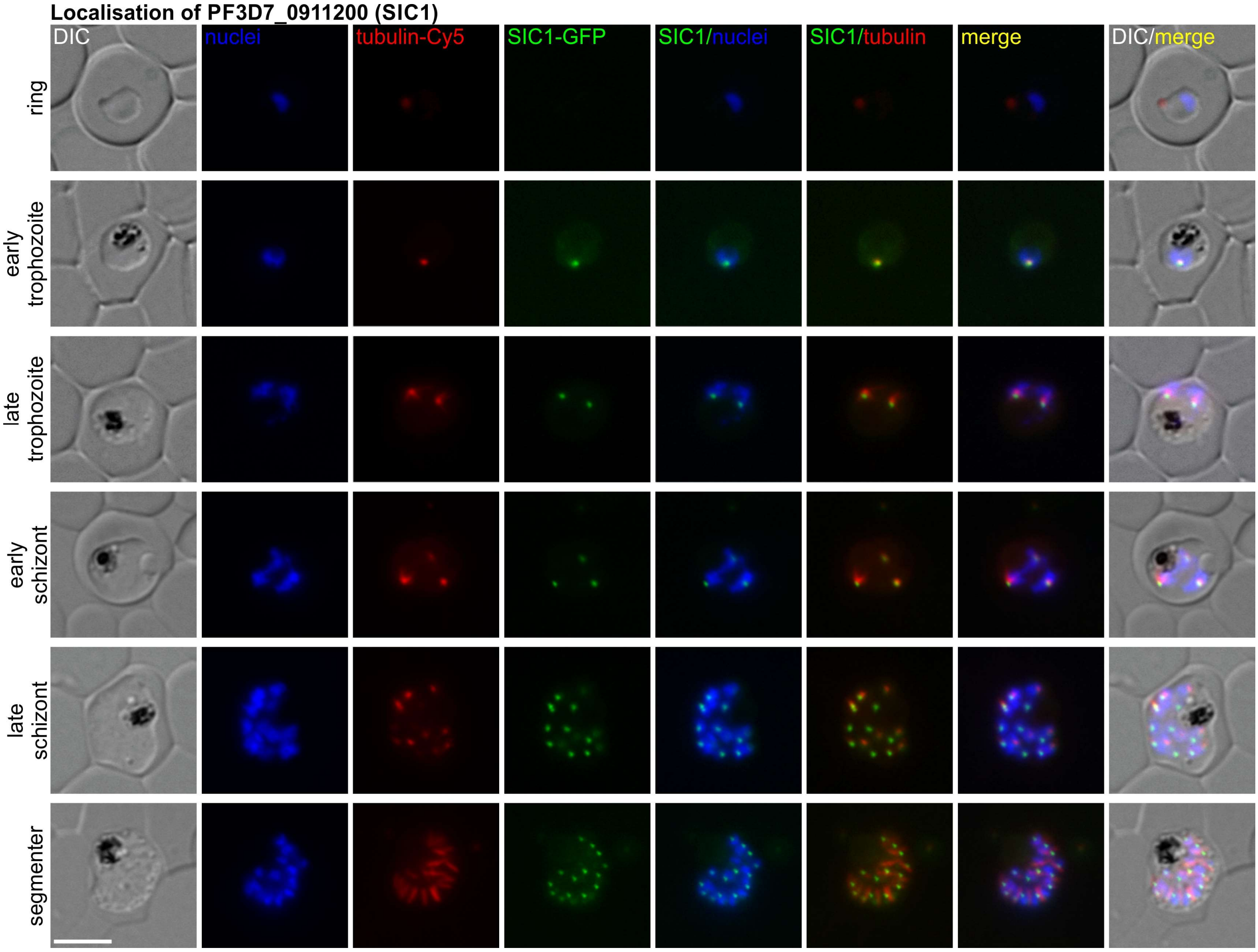

**Localisation of PF3D7_1349600 (SIC2)**

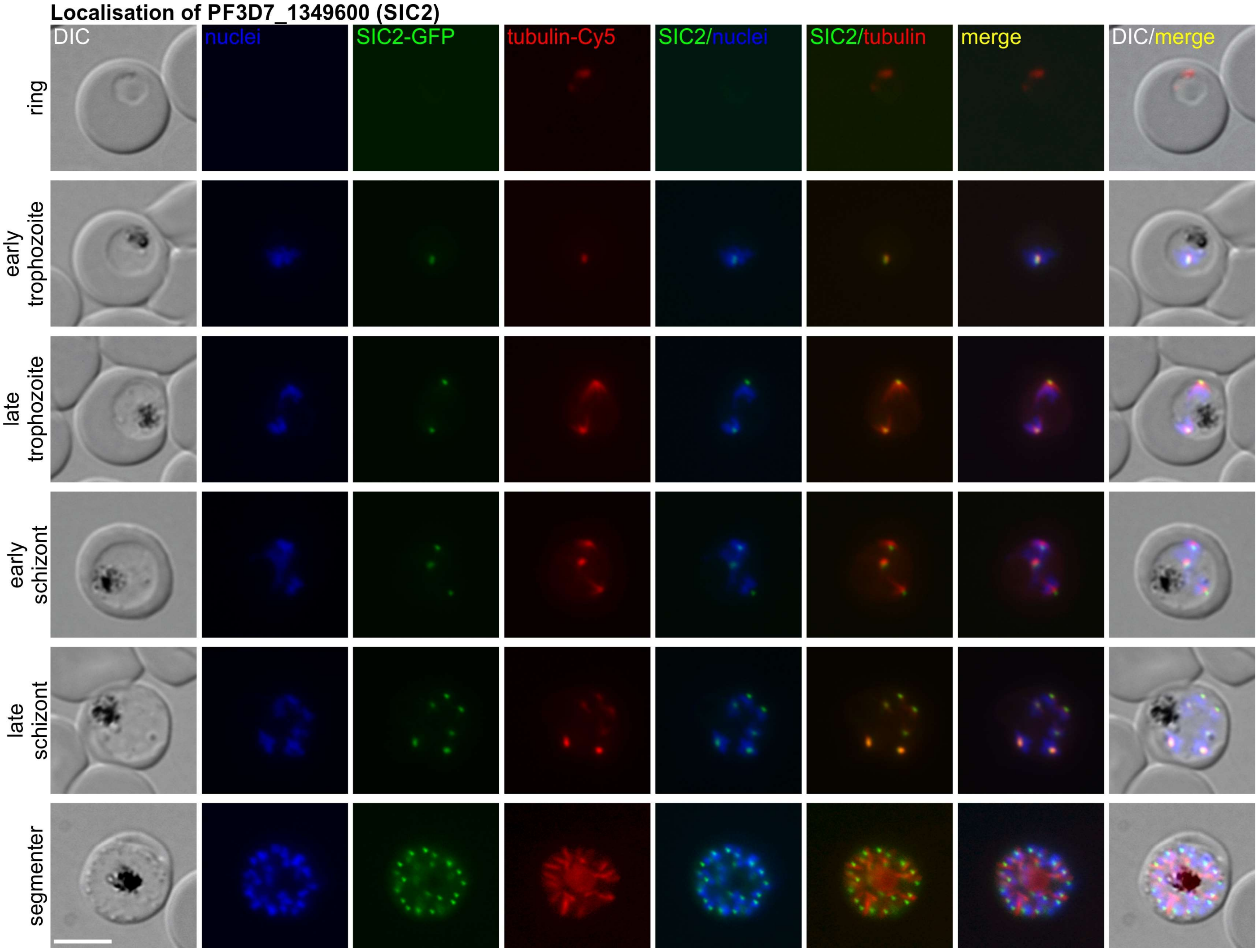

**Localisation of PF3D7_1331200 (SIC3)**

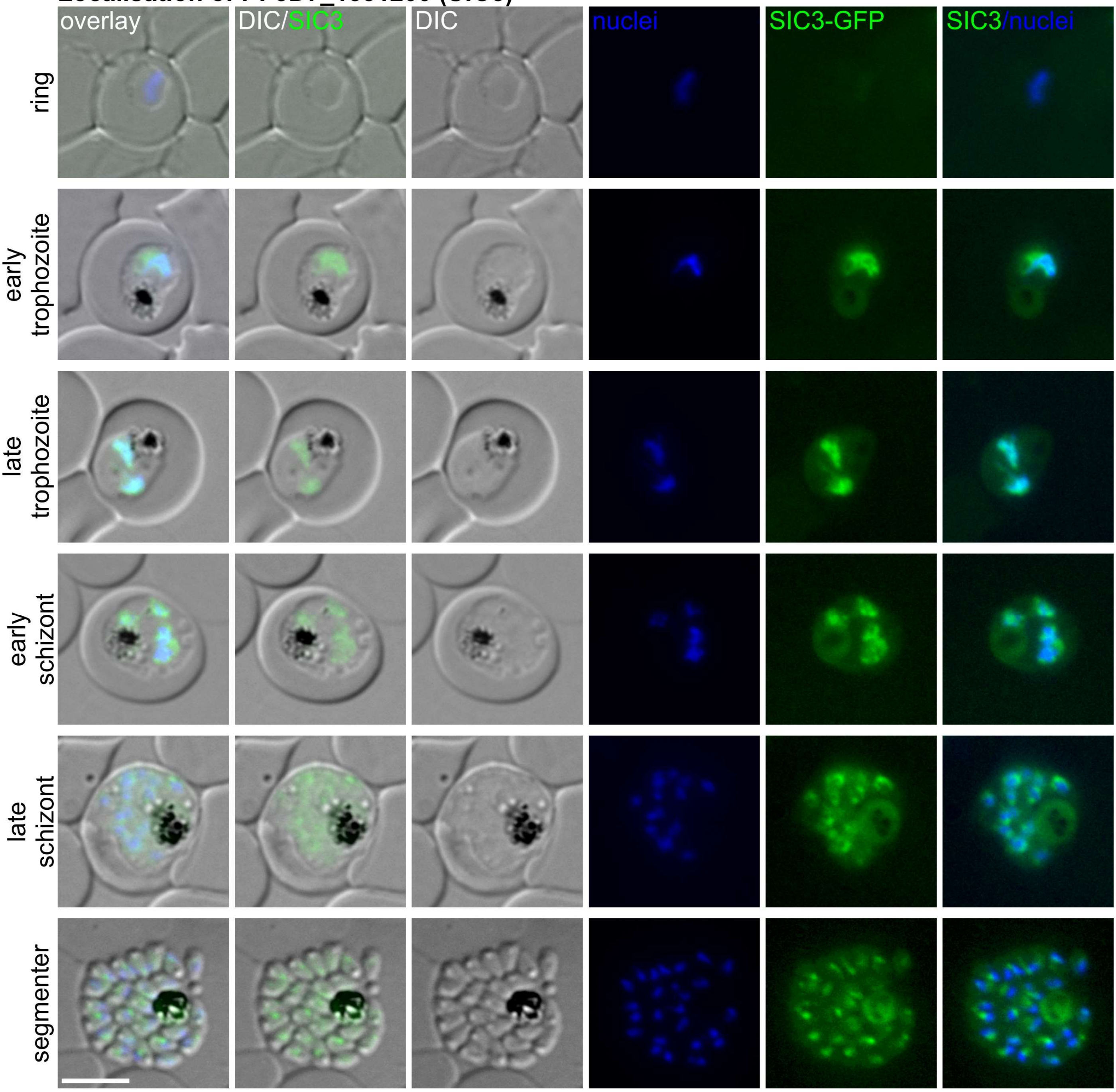

**Localisation of PF3D7_1446700 (SIC4)**

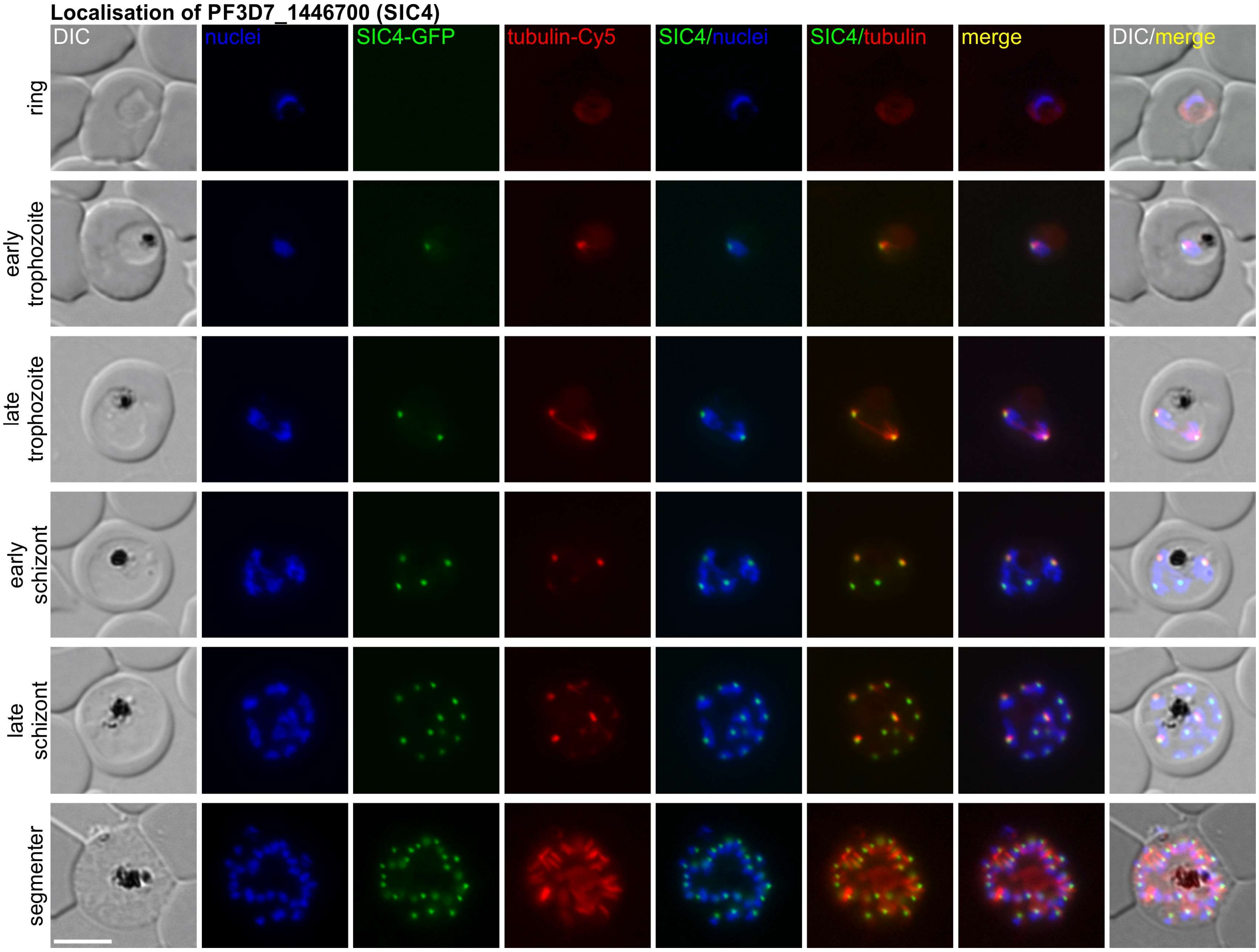

**Localisation of PF3D7_1227600 (SIC5)**

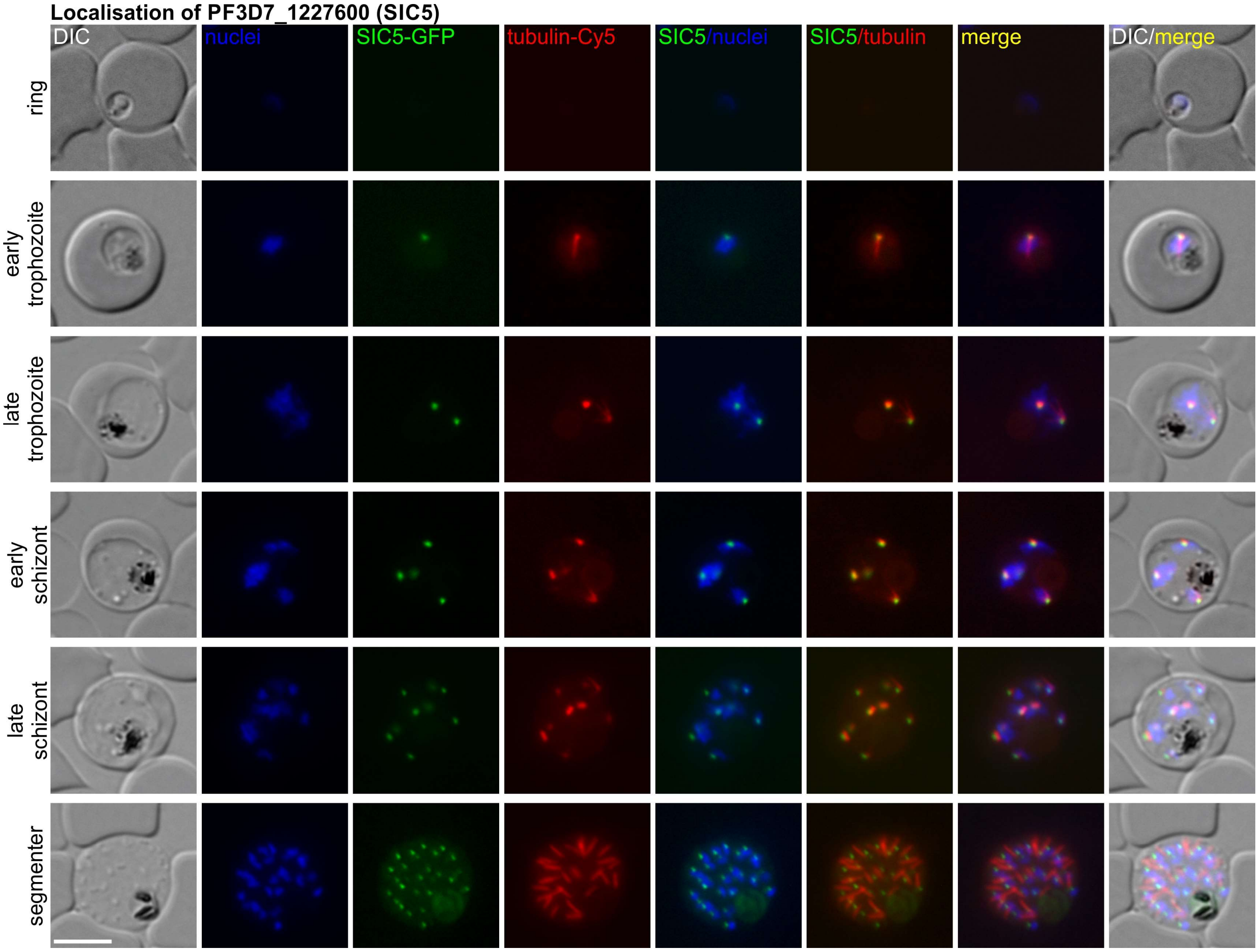

**Localisation of PF3D7_0616200 (NDC80)**

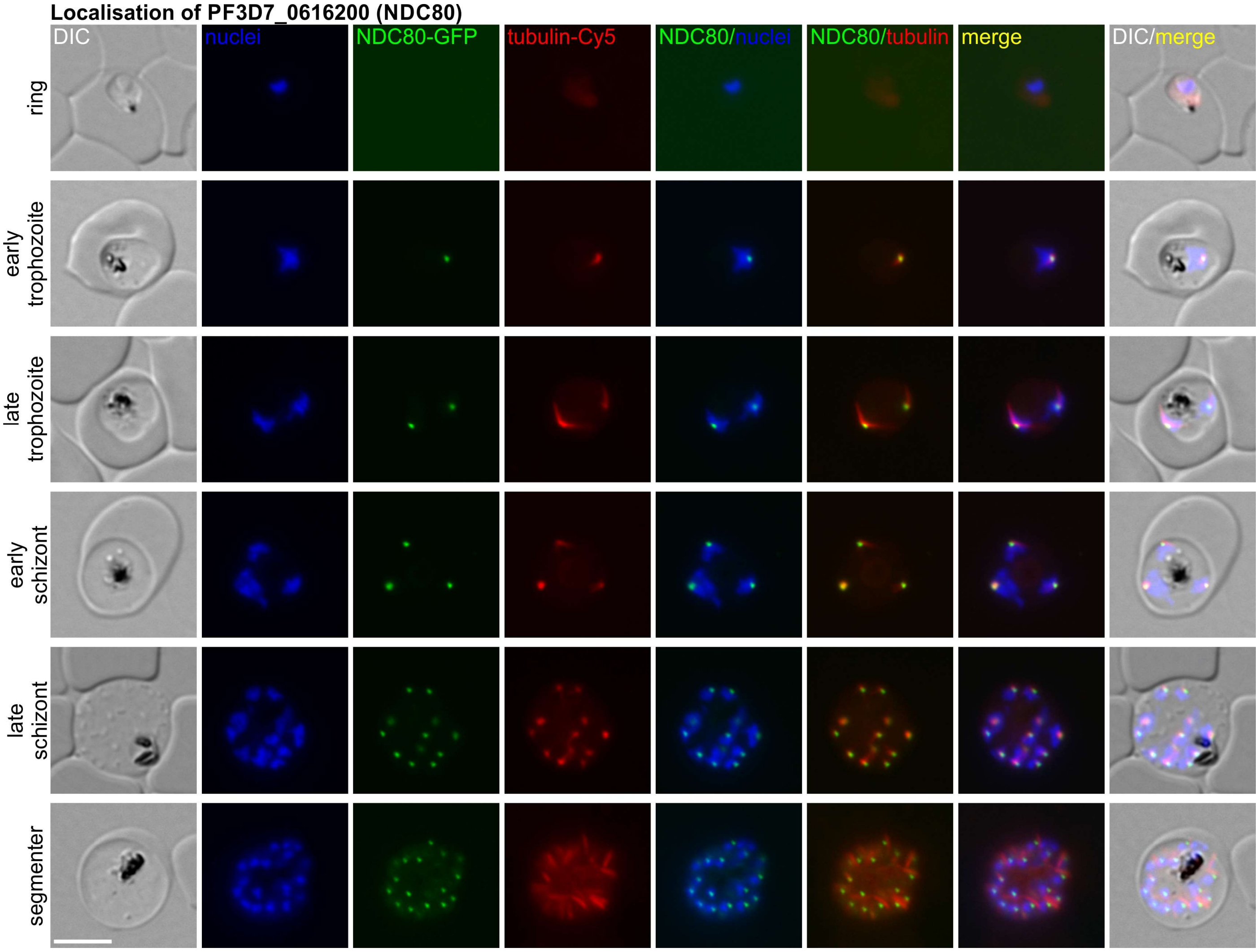
